# SubcellPert: Benchmarking Subcellular Perturbation Prediction

**DOI:** 10.64898/2026.10.11.758305

**Authors:** Jinghao Wang, Hanqun Cao, Xilin Dang, Tianhui Zhou, Yiyang Zhao, Chang-Yu Hsieh, Chunbin Gu, Pheng Ann Heng

## Abstract

Perturbations change both RNA abundance and intracellular organization, but gene-count predictors omit the positions needed to model spatial responses. To address this limitation, we introduce ***SubcellPert***, a benchmark for perturbation-response prediction at single-RNA-molecule resolution. It comprises 14 imaging datasets with 13.88 million cells and 3.65 billion RNA molecules. Given source observations and a directed condition change, models generate target RNA populations in normalized two-dimensional coordinates. Two independently trained tasks separate abundance from localization: *Oracle* predicts coordinates given exact target counts; *Joint* predicts both. Eight baselines span source resampling, classical transport, and frozen foundation-model adapters. Fixed perturbation and cellular-context hold-outs evaluate abundance error, target-gene coverage, and conditional spatial error. Through within-dataset analyses, we find context-dependent gains from spatial features in condition decoding. Interpreting spatial responses requires distinguishing changes within spatial clusters from population composition, and RNA fractions from enrichment relative to compartment geometry. Through comprehensive experiments, we find that Gaussian transport achieves the lowest Oracle spatial error on eight of eleven cellular-context OOD datasets. Audits show that better centroid accuracy can accompany worse radial distributions, and that 19 of 190 Joint adapter fits retain source counts after validation. SubcellPert opens a spatial frontier for virtual cell research, laying the groundwork for predictive models that capture both the molecular composition and intracellular organization of cellular responses. The code is available at https://github.com/king159/SubcellPert.

## 1 Introduction

Cells regulate both RNA abundance and transcript location, which helps determine where proteins can be produced. RNA localization contributes to migration (Wang et al., 2017), activity-dependent neuronal translation (Buxbaum et al., 2014), and stress-granule recruitment (Khong et al., 2017); drug treatment can also alter localization (Mah et al., 2024). Predicting a perturbation response therefore requires considering both how much RNA a cell contains and how it is organized.

Gene counts, the primary representation in current perturbation predictors and benchmarks (Lotfollahi et al., 2019; Roohani et al., 2024; Wei et al., 2026), describe how much RNA a cell contains but not how that RNA is organized: cells with identical counts can have different intracellular distributions. Molecule-resolved imaging now measures RNA identities and coordinates under perturbations (Chen et al., 2015; Binan et al., 2025), and subcellular methods characterize observed localization (Samacoits et al., 2018; Mah et al., 2024; Wang & Zhou, 2025). These measurements and analyses do not by themselves establish how to evaluate spatial responses predicted for held-out perturbations or cellular contexts. The challenge is to compare unpaired source and target populations while distinguishing errors in RNA abundance, gene coverage, and spatial distribution: count predictions determine how many coordinates exist and which genes can be spatially evaluated.

Here we introduce ***SubcellPert***, a benchmark for predicting subcellular responses from unpaired populations. Figure 1 summarizes its prediction inputs and outputs, Oracle and Joint settings, dataset holdouts, and baseline families. The collection contains 13.88 million cells and 3.65 billion RNA molecules across genetic interventions, treatments, and condition changes. Models generate target populations from source observations and a directed condition change. Independently trained *Oracle* supplies target counts to isolate localization; *Joint* predicts counts and coordinates, with coverage tracking omitted genes. Three datasets hold out interventions or combinations; eleven hold out responses in cellular contexts with source observations available.

**Figure 1:**
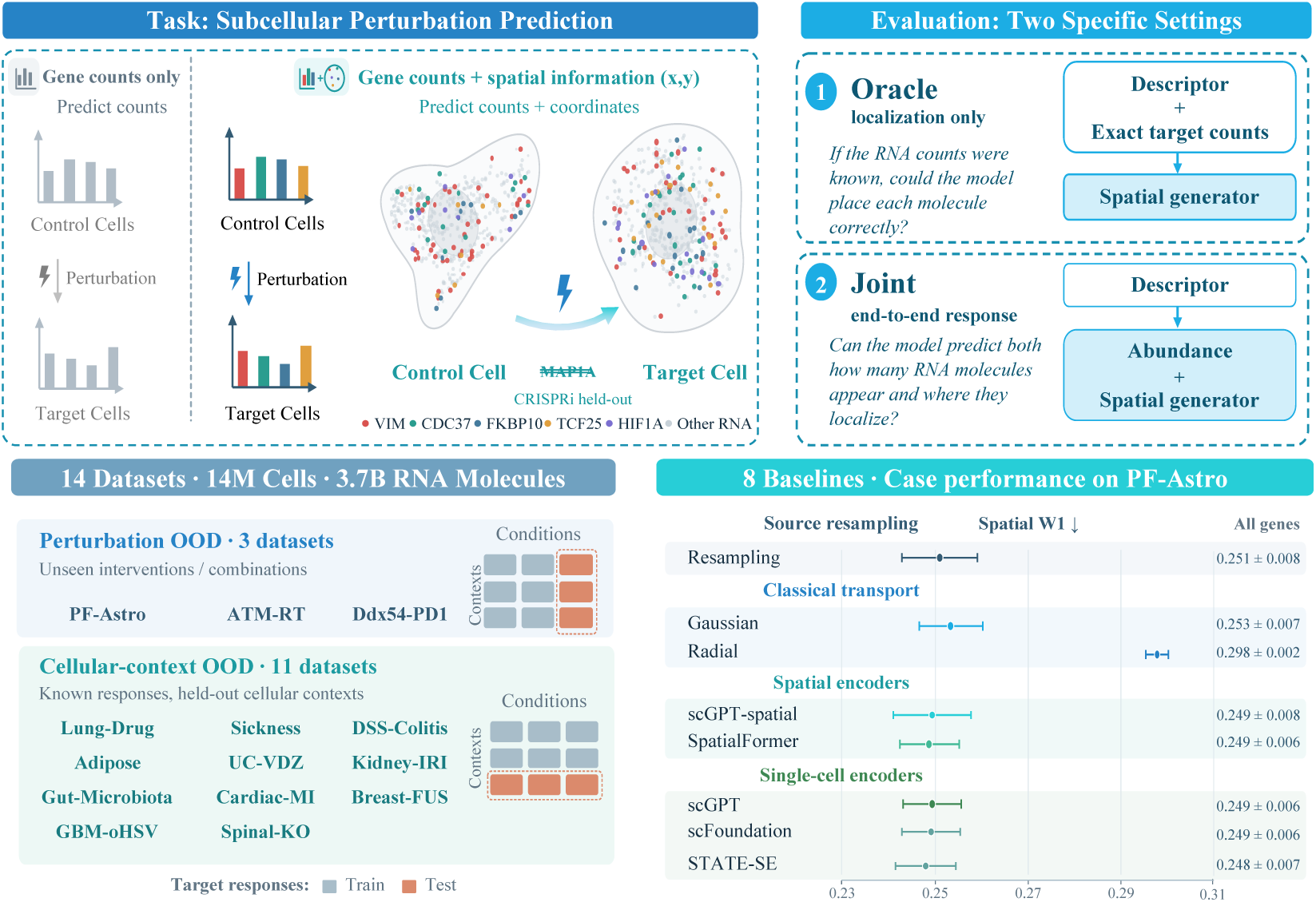
Overview of SubcellPert. **Top left:** Identical RNA abundance can conceal distinct RNA arrangements. RNA dot colors identify genes, with gray denoting other RNA; pale cell membranes and nuclei are schematic. MAP1A illustrates a held-out CRISPRi perturbation; control and target cells are unpaired. **Top right:** Oracle predicts coordinates given target counts; Joint predicts abundance and coordinates from source observations and a condition change. **Bottom left:** Fourteen datasets: three for Perturbation OOD and eleven for Cellular-context OOD. Matrix rows denote cellular contexts and columns denote conditions; blue and orange mark training and test target responses. Holding out columns illustrates unseen perturbations, while holding out rows illustrates unseen target responses in cellular contexts whose source observations remain available. **Bottom right:** PF-Astro Oracle all-gene spatial W1 (lower is better) for eight baselines: mean and sample standard deviation across three splits, each averaging 18 held-out responses.

To understand what spatial measurements contribute beyond gene counts and how observed RNA responses should be interpreted, we conduct descriptive analyses within each dataset. To assess how well existing methods predict these responses under held-out perturbations and cellular contexts, we evaluate eight baselines and audit their predicted counts and coordinates to understand the behaviors underlying their benchmark scores.

Our contributions are as follows:

- **A subcellular perturbation prediction benchmark.** To our knowledge, SubcellPert is the first benchmark for predicting gene-identified RNA point clouds after perturbation, supported by 14 curated imaging datasets.
- **Comprehensive evaluation of distinct sources of performance.** Our benchmark includes eight baselines, gene-panel comparisons, coverage-aware metrics, and separate Oracle and Joint tasks to isolate localization from abundance prediction.
- **Characterization of observed and predicted RNA responses.** We connect within-dataset analyses of RNA organization with audits of model predictions to examine how biological context informs spatial interpretation and how prediction behavior shapes benchmark performance.

## 2 Related work

### 2.1 Perturbation-response prediction

Single-cell response models predict post-intervention abundance. scGen (Lotfollahi et al., 2019) transfers a response in latent space, CPA (Lotfollahi et al., 2023) composes perturbation, dose, and covariate effects, and CellOT (Bunne et al., 2023) maps unpaired control and perturbed populations. GEARS (Roohani et al., 2024) and TxPert (Wenkel et al., 2026) use gene-relation graphs for unseen genetic or combinatorial effects; pretrained models such as scGPT (Cui et al., 2024) and scFoundation (Hao et al., 2024) provide transferable abundance representations.

Spatial response models add tissue context but retain an abundance target. CONCERT (Lin et al., 2025), for example, propagates perturbation effects through cellular niches while predicting cell-or spot-level abundance. Its coordinates are contextual inputs, not predicted intracellular RNA locations. The output distinction relevant here is a count matrix versus a population of gene-identified intracellular RNA coordinates.

### 2.2 Subcellular RNA modeling

Subcellular methods analyze observed molecule coordinates. The spatial-statistics framework of Samacoits et al. (2018) and Bento (Mah et al., 2024) characterize localization patterns, domains, and colocalization. PointFISH (Imbert et al., 2023) learns point-cloud representations, Focus (Lu et al., 2025) constructs RNA-neighborhood graphs, and ComSeg (Defard et al., 2024) uses molecule graphs for segmentation. FISHFactor (Walter et al., 2023) provides probabilistic spatial factors, and ELLA (Wang & Zhou, 2025) tests within-cell spatial variation. scGPT-spatial (Wang et al., 2025) and SpatialFormer (Wang et al., 2026) extend representation learning across subcellular and tissue scales. These methods embed, segment, or explain an observed measurement; they do not learn its intervention-conditioned future population.

## 3 SubcellPert

### 3.1 Problem Formulation

#### Representation and conditioning

Let *X_i_* = {(*g,* **r***_igk_*): *g* ∈ *G, k* = 1*…, n_ig_*} represent cell *i*, where *n_ig_* counts gene *g* in vocabulary *G* and **r***_igk_* ∈ ℝ^2^ is its normalized RNA coordinate. Source 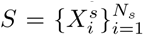 and target 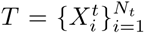 populations are unpaired. All RNA within selected cells are retained; the count projection *∁*(*X_i_*) = (*n_ig_*)*_g_*_∈G_ discards their arrangement. Coordinates are centered at the all-RNA centroid and scaled by the cell’s RMS radius, describing relative organization rather than absolute size or annotated compartments, as detailed in Appendix A.2. The supplied descriptor *d* identifies the target condition; the source condition fixes response direction. Its condition indicators, intervention combinations, or gene-relation features are constructed without held-out target abundance or geometry, as specified in Appendix A.3.

##### Response factorization

For an output-cell budget *B*, let N and R denote output counts and gene-identified coordinates. The response distribution factorizes as

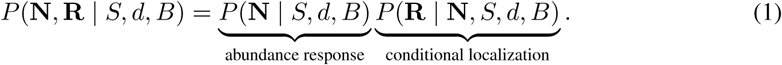

This separates errors in RNA quantity from placement. Table 1 contrasts the two settings and their access to target counts.

**Table 1:** Task comparisons between SubcellPert settings and previous tasks. RNA configurations *X* specify gene identities and intracellular coordinates. Both settings withhold target geometry and share a fixed output-cell budget; only Oracle receives target counts.

| Task | Observed input | Perturbation $d$ | Target counts at inference | Required prediction |
| --- | --- | --- | --- | --- |
| Perturbation prediction | Source counts $\mathbf{n}^s$ | ✓ | Hidden | Target counts $\hat{\mathbf{n}}^t$ |
| Subcellular Spatial Transcriptomics | Observed RNA configuration $X$ | × | Not applicable | Localization features / representations |
| SubcellPert *Oracle* | Source RNA configuration $X^s$ | ✓ | **Given** $\mathbf{n}^t$ | Target coordinates; counts fixed |
| SubcellPert *Joint* | Source RNA configuration $X^s$ | ✓ | **Hidden** | Target abundance **and** coordinates |

### 3.2 Prediction Settings and OOD Tasks

#### Oracle: conditional localization

Oracle receives the source population, the transition descriptor, and one exact target count vector per output cell:

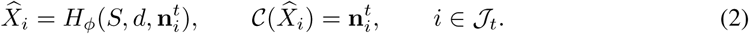

Abundance is a constraint rather than a prediction target, isolating localization from errors in abundance prediction. The target index specifies a count vector, not a matched source cell.

#### Joint: abundance and localization

Joint does not receive target counts. Given source observations, *d*, and a declared output-cell budget, it predicts a count vector and then generates the corresponding gene-identified coordinates:

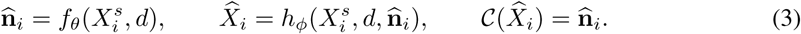

Both settings generate *B* = min(2000*, N_t_*) cells using the same preselected target indices *J_t_*. Joint receives only this budget as target metadata; source draws are independent of target abundance and geometry. The budget does not model proliferation. Neither setting receives target coordinates or target-derived normalization. Independent training means Oracle is not a strict upper bound on Joint. Zero-RNA outputs remain in abundance evaluation without fabricated geometry. Appendix A.1 details sampling and prediction-freezing rules.

#### Cellular-context OOD

It predicts a known response in a held-out cellular context. Treated cells from that context are excluded from fitting, while its source controls remain available.

#### Perturbation OOD

It predicts an unseen intervention or combination within an observed context.

### 3.3 Benchmark Construction

#### Retrieval and inclusion

Figure 2 shows how we screened OpenAlex and Europe PMC with manual supplementation, then reconstructed eligible releases. Admission required gene-resolved RNA coordinates, recoverable molecule-to-cell assignments, released condition and context annotations, auditable counts, and a supported source–target response within one experimental branch. Quality control follows source-specific molecule and cell filters, checks gene identifiers against the retained assay vocabulary, and compares reconstructed counts with released count objects where available. Structural checks verify finite coordinates, valid cell/gene indices, and agreement between molecule totals and gene counts; cells without eligible RNA are excluded. Before release, context–condition groups with fewer than 50 cells are removed, pooling eligible units within each dataset. Each evaluated response additionally requires at least 50 target and 50 matched source cells in the same context and anatomical reference where applicable. Retrieval provenance, quality-control details, and exclusion decisions appear in Appendix B.

**Figure 2:**
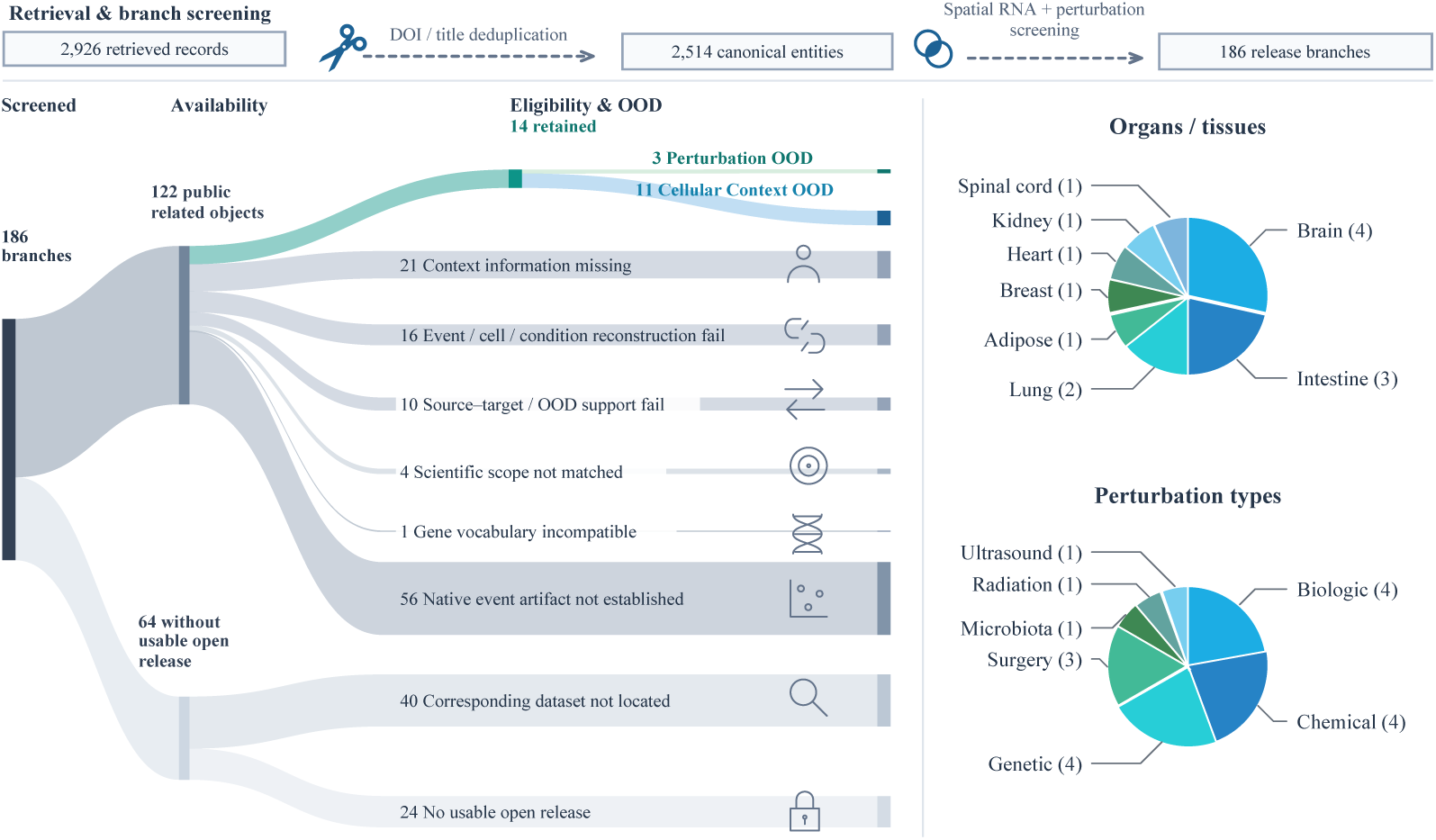
Dataset selection and benchmark diversity. Dashed arrows connect retrieval stages with different counting units; solid ribbons conserve 186 release branches. Of 122 branches passing the usable-open-release screen, 14 are retained: 11 Context OOD and three Perturbation OOD. Further details are in Appendix B.

#### Scale and diversity

Table 2 reports the 14 datasets, comprising 13.88 million nonempty cells and 3.65 billion RNA molecules with each assay’s gene panel retained. They span human and mouse tissues profiled by MERFISH, Xenium, and CosMx, with genetic, drug, immune, injury, microbiota, and weight-loss contrasts. PF-Astro (Binan et al., 2025) holds out single-gene perturbations (87 supported identities, 18 test identities per split); ATM-RT and Ddx54-PD1 hold out combined interventions in 2 × 2 designs. The other 11 datasets hold out cellular contexts. Individual experimental designs and interpretation limits are described separately in Appendix C. We provide some examples of each dataset in Appendix D.

**Table 2:** Released SubcellPert populations. It contains 13,880,723 cells and 3,653,986,724 eligible RNA across 14 retained releases.

| Release | # RNAs | # Cells | # Genes | Unit/group labels | # Perturbations/<br>Cellular Contexts |
| --- | --- | --- | --- | --- | --- |
| *Perturbation OOD* |  |  |  |  |  |
| PF-Astro | 10,670,839 | 11,454 | 485 | 3 wells | 87 |
| ATM-RT | 147,722,071 | 790,166 | 298 | 4 sections | 3 |
| Ddx54-PD1 | 494,265,984 | 1,155,233 | 5,006 | 4 sections | 3 |
| *Cellular-Context OOD* |  |  |  |  |  |
| Lung-Drug | 681,015,170 | 1,561,700 | 5,006 | 8 sections | 11 |
| Sickness | 1,101,298,121 | 1,765,443 | 941 | 16 animals | 103 |
| DSS-Colitis | 119,553,365 | 1,439,542 | 940 | 17 mice | 8 |
| Adipose | 1,907,755 | 13,972 | 477 | 4 paired donors | 7 |
| UC-VDZ | 19,552,710 | 80,562 | 1,000 | 7 patients | 28 |
| Kidney-IRI | 168,519,194 | 1,374,915 | 300 | 6 mice | 20 |
| Gut-Microbiota | 189,197,264 | 2,057,563 | 1,815 | 28 acquisition groups | 75 |
| Cardiac-MI | 61,134,068 | 225,023 | 475 | 7 slides | 67 |
| Breast-FUS | 102,102,171 | 880,565 | 1,000 | 2 conservative pools | 46 |
| GBM-oHSV | 520,974,197 | 2,401,940 | 480 | 8 patients | 11 |
| Spinal-KO | 36,073,815 | 122,645 | 347 | 8 sample animals | 158 |
| **Total** | **3,653,986,724** | **13,880,723** | — | — | — |

**Table 3:** All-gene cross-dataset mean ranks. Ranks use all eight methods and every dataset in the stated axis, with average ranks for ties.

| Baseline | Perturbation OOD |  |  | Cellular-context OOD |  |  |
| --- | --- | --- | --- | --- | --- | --- |
|  | O-W1 ↓ | J-MSE ↓ | J-PCC ↓ | O-W1 ↓ | J-MSE ↓ | J-PCC ↓ |
| Resampling | 5.00 | 1.17 | 4.50 | 6.73 | 3.36 | 7.18 |
| Gaussian | 4.67 | 3.17 | 4.83 | 2.36 | 1.95 | 2.95 |
| Radial | 8.00 | 3.17 | 4.83 | 7.45 | 1.95 | 2.95 |
| scGPT-spatial | 3.33 | 6.67 | 4.33 | 4.09 | 5.64 | 4.73 |
| SpatialFormer | 3.67 | 6.67 | 3.00 | 4.18 | 5.73 | 3.64 |
| scGPT | 4.33 | 6.00 | 3.67 | 3.91 | 6.27 | 5.82 |
| scFoundation | 3.67 | 4.83 | 6.17 | 4.00 | 5.18 | 4.36 |
| STATE-SE | 3.33 | 4.33 | 4.67 | 3.27 | 5.91 | 4.36 |

### 3.4 Baselines

We compare eight baselines, fitted independently in Oracle and Joint. **Source resampling** provides a parameter-free reference. **Classical transport** comprises Gaussian affine and radial quantile transport, which learn condition-dependent transformations from cell-level spatial summaries. The five **foundation-model baselines** use single-cell encoders scGPT (Cui et al., 2024), scFoundation (Hao et al., 2024), and STATE-SE (Adduri et al., 2025) (SE-600M), or spatial encoders scGPT-spatial (Wang et al., 2025) and SpatialFormer (Wang et al., 2026).

All five encoders remain frozen and provide contextual gene states to a shared GEARS-based adapter (Roohani et al., 2024), following scPerturBench (Wei et al., 2026). It combines source counts, RNA geometry, and condition descriptors to predict gene-specific affine transformations of RNA coordinates. Only the adapter and output heads are trained, for five epochs with a fixed optimizer batch size of 256. Baseline implementations, losses, and training details are provided in Appendix E.

### 3.5 Evaluation Protocol

Oracle is evaluated by spatial sliced 1-Wasserstein distance (W1). Joint also uses abundance MSE and source-delta Pearson correlation (PCC-delta), computed from population mean log(1 + count) including zeros. Spatial W1 averages 64 projected distances over target-present genes for Oracle and shared nonempty genes for Joint. Joint target-gene coverage (Cov.) is the fraction of target-present genes with predicted RNA; absent genes have no spatial error to score.

We score all-gene and top-100 abundance-ranked panels, fixing predictions before using held-out observations for scoring or ranking. All generated cells and RNA are evaluated. Twelve datasets have three frozen split seeds; ATM-RT and Ddx54-PD1 have one compositional holdout each. Scores weight responses, families, and splits equally at their respective aggregation levels. Split standard deviations measure partition variation, not biological replication. Appendix F gives the equations, weighting, and missing-value rules.

## 4 Experiments

### 4.1 Within-dataset Observations of RNA Organization

We characterize observed RNA responses within each dataset; Appendix G gives complete settings and further analyses. Each point is a context–condition comparison within one dataset, except condition decoding, which uses one point per context across its supported conditions.

#### Abundance and spatial changes

In Figure 3a, we compare relative abundance and spatial response magnitudes on matched genes in 1,628 supported population comparisons. A separate gene-level analysis gives Spearman correlations of 0.264 in Adipose adipocytes, 0.086 in GBM tumor cells, and 0.139 in GBM macrophages/monocytes. These weak, differently signed associations do not establish independence.

**Figure 3:**
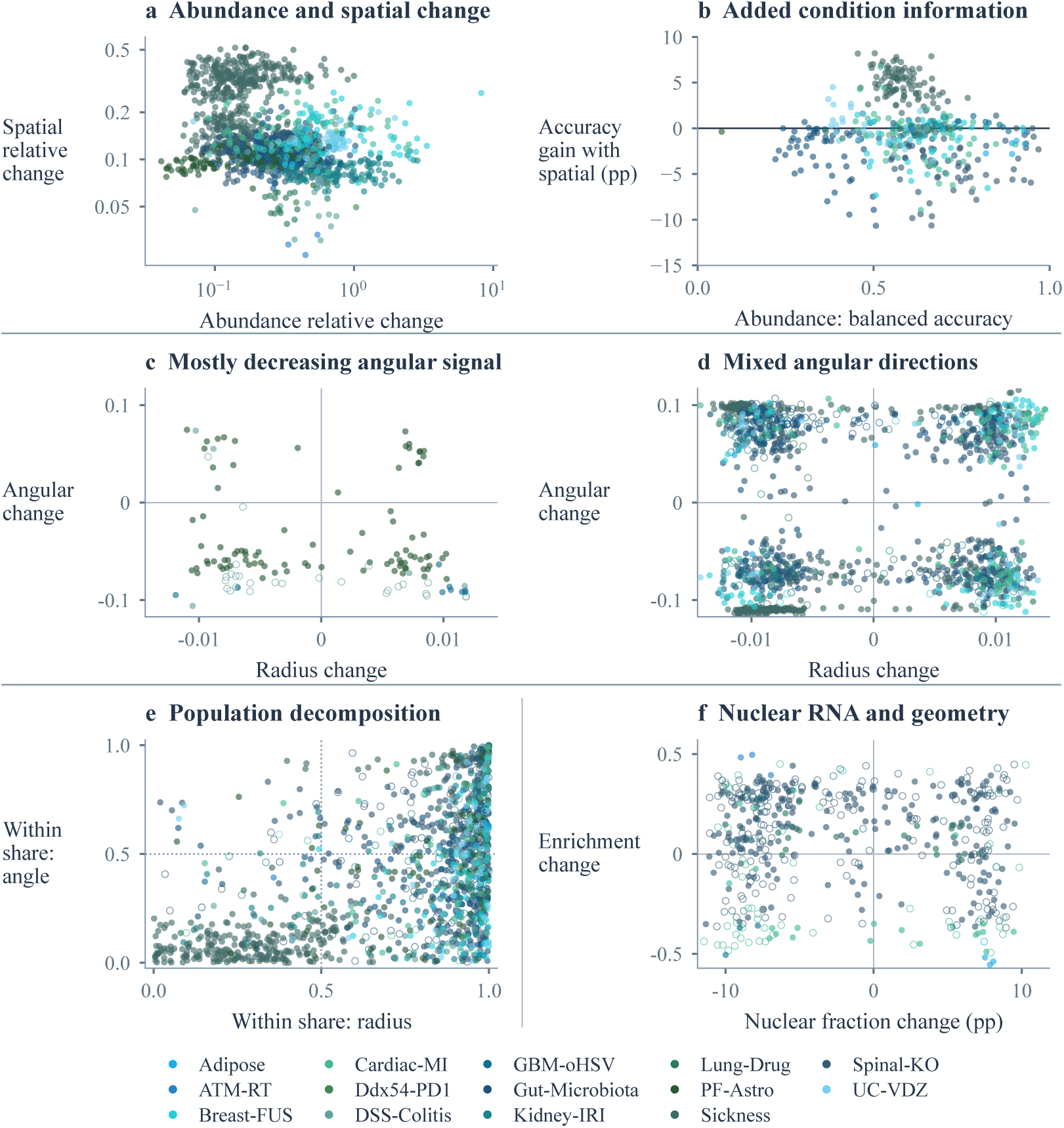
Within-dataset observations of RNA responses. **a:** Relative abundance and spatial changes on matched genes. **b:** Abundance-only balanced accuracy versus its gain after adding spatial principal components; points average three stratified cell splits, and positive gains favor spatial addition. **c–d:** Radial and angular changes in 1,622 comparisons from 12 datasets with enough comparisons for descriptive grouping. **e:** Within-cluster contribution shares, |*W| /*(*|W|* + *|C|*), for radius and angle; *C* denotes cluster composition. Dotted lines at 0.5 mark equal absolute contributions. Shares omit effect direction and net magnitude. **f:** Median gene-level changes in nuclear RNA fraction and enrichment normalized by the nuclear-to-cell projected-area ratio. Colors identify datasets; no pooled effect is estimated. Hollow circles flag median angular-valid cell fractions *<* 20% in **c–d**, minimum clustering-initialization ARI *<* 0.8 in **e**, or *<* 20 supported genes in **f**; they do not mark significance. Panel **a** uses log axes; **c–d** and **f** use symmetric-log axes.

#### Incremental condition information

In Figure 3b, we compare condition classification with and without spatial information using 64 principal components per branch and three stratified 80:20 cell splits per context. Sickness improves in 101/102 contexts, with a median balanced-accuracy gain of 4.85 percentage points; GBM-oHSV improves in 7/8, with a median gain of 0.67 points. Adipose and DSS-Colitis have negative mean gains in all six and eight contexts, respectively, while Spinal-KO has a median gain of 3.01 points. These results concern observed-condition classification, not prediction of unseen donors or perturbations.

#### Radial and angular patterns

In Figure 3c–d, we separate changes in normalized mean RNA radius from changes in angular concentration relative to other RNA. DSS-Colitis, GBM-oHSV, and PF-Astro have predominantly negative angular changes in panel c. Adipose and eight other datasets have mixed directions in panel d. Low angular observability in DSS-Colitis limits interpretation of its apparent direction.

#### Population composition and within-cluster changes

In Figure 3e, we separate changes in cluster proportions from changes within fixed spatial clusters and plot their relative absolute contributions. In Breast-FUS, within-cluster terms dominate radius changes in 66/68 comparisons, while composition terms dominate angular changes in 51/68. Adipose has within-cluster radius dominance in all six comparisons but an even split for angle. In Sickness, composition terms dominate 238/331 radius and 274/331 angular comparisons, with condition-dependent heterogeneity. These operational clusters do not establish biological states or cell trajectories.

#### Nuclear fraction and compartment geometry

In Figure 3f, we compare changes in nuclear RNA fraction with enrichment relative to nuclear area. Adipose has higher median fractions but lower median enrichment in four contexts; adipocyte enrichment is sensitive to one donor. ATM-RT shows the reverse directions in all three comparisons. GBM tumor cells and macrophages/monocytes decrease in both quantities. In Lung-Drug airway epithelium, the two drugs have opposite fraction changes but both increase enrichment; two Spinal-KO perturbations show the same contrast within the v1 context. Cardiac-MI spans all four sign combinations, with limited gene support in 54/76 comparisons. These descriptive differences concern fractions and projected geometry, not absolute nuclear counts or RNA transport.

### 4.2 Baseline Performance and Prediction Behavior

Eight baselines are evaluated in independently fitted Oracle and Joint settings: 608 runs over 14 datasets and 38 frozen split instances. Figure 4 summarizes performance; Appendix H gives complete scores. We relate these scores to observed responses and audit saved RNA counts and coordinates.

**Figure 4:**
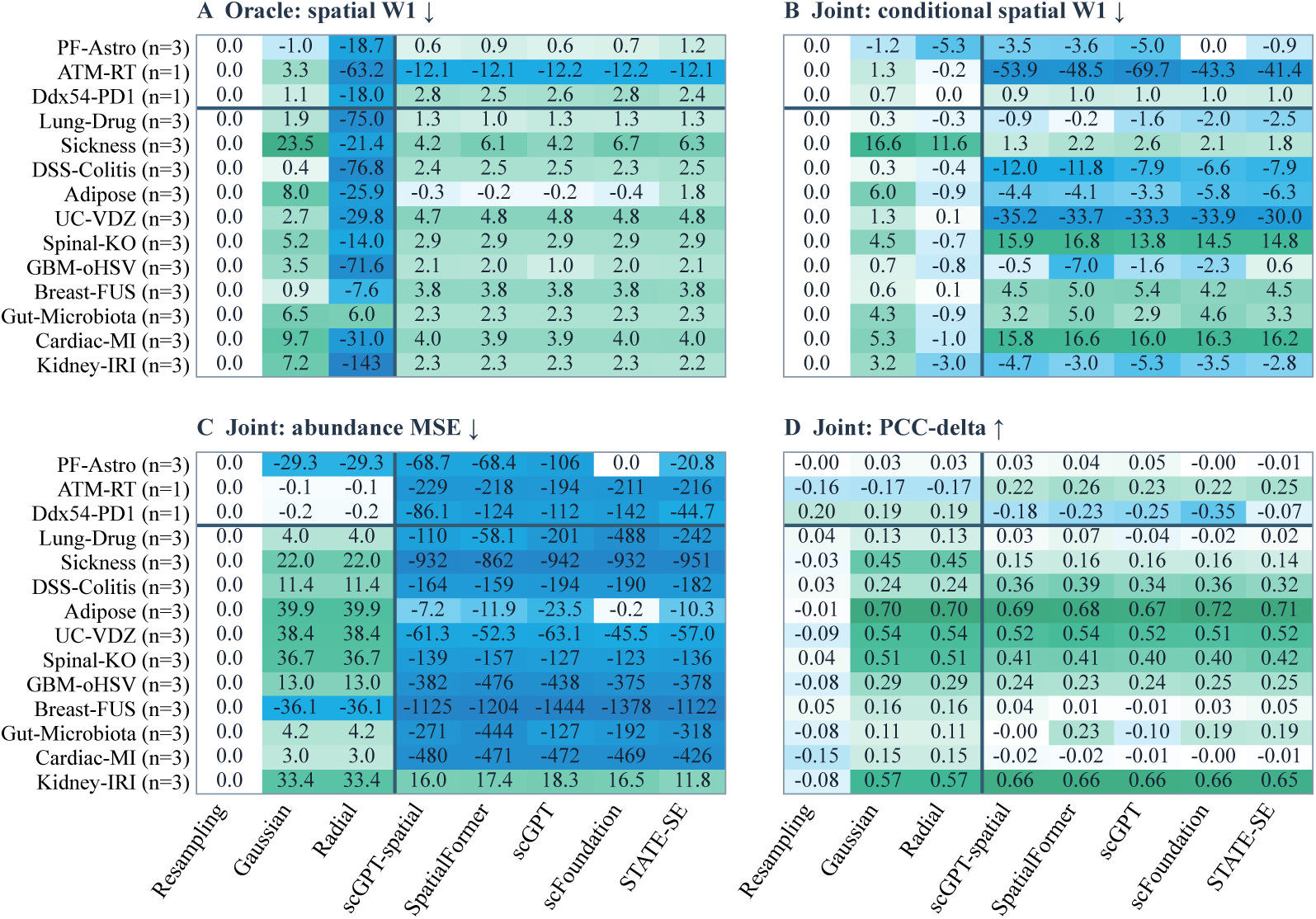
All-gene benchmark results. **A–C**: show percentage improvement over Resampling; **D** shows PCC-delta. Title arrows indicate the preferred direction of the raw metrics.

#### Spatial performance and a common contraction

Gaussian has the lowest Oracle W1 on eight of eleven Cellular-context OOD datasets. In Adipose, mean responses from other contexts improve abundance in 6/6 and radius in 4/6 comparisons; in Sickness, spatial features improve observed-condition decoding in 101/102 contexts but mean radius and angular responses from other contexts improve none of 202 comparisons (Figure 3 and Appendix G.7). Gaussian leads Oracle W1 in both (Tables 26 and 28); these tests do not identify why it succeeds. FM predictions contract in 667/685 estimable model–response cases, typically to a scale near 0.90. On ATM-RT, centroid MSE improves about 8% while radial W1 worsens about 34% relative to Resampling: mean-location accuracy can mask distributional error. Gaussian often preserves overall radius while adapting gene-specific means and covariances, without implying a whole-cell transform.

#### Abundance changes: magnitude and direction

Gaussian and Radial share identical abundance regression and therefore identical Joint counts in all 138 test responses. Among FM fits, validation selects epoch zero in 19/190 cases, leaving donor counts unchanged in all 127 associated responses. Other fits overshoot or misdirect changes: on ATM-RT, predicted change norms are 1.78–1.90 times observed norms, with directional cosines only 0.34–0.37. On Ddx54-PD1, all five FM change directions have negative cosine. These behaviors help explain the competitiveness of source-preserving references.

#### Shared predictions and evaluation scope

Within individual datasets, median pairwise FM abundance-change cosines are 0.68–0.99, and Oracle per-gene error-rank correlations are 0.978–1.000. Under the shared adapter, these similarities establish neither encoder-specific biological utility nor encoder irrelevance.

#### Gene panels and coverage

Of 61 positive FM–dataset Oracle gains, 31 reverse on the top-100 abundance-ranked panel (Appendix H.1). On Spinal-KO, SpatialFormer lowers conditional Joint W1 relative to Gaussian (0.312 vs. 0.358) but has lower target-gene coverage (76.5% vs. 93.5%) and higher abundance MSE (0.0372 vs. 0.0117; Table 30). Joint W1 scores localization among recovered genes; Figure 9 reports coverage across datasets. Rankings concern the frozen-encoder, five-epoch protocol; the two compositional tasks have one fixed split.

## 5 Conclusion

In this paper, we propose **SubcellPert**. It benchmarks abundance and spatial response prediction across 14 imaging datasets, enriching the previous perturbation-prediction literature with subcellular resolution. The collection contains 13.88 million cells and 3.65 billion RNA molecules across genetic interventions, treatments, and condition changes. Within-dataset observations distinguish population composition from within-cluster changes, show context-dependent decoding gains, and separate nuclear RNA fraction from area-normalized enrichment. These descriptive findings motivate separate abundance, spatial, and coverage evaluation without establishing shared mechanisms. Comprehensive experiments with eight baselines show that classical references remain competitive under the frozen-adapter protocol; prediction audits reveal spatial contraction, abundance-change misalignment, and exact source preservation.

SubcellPert opens a spatial axis for virtual cell research by making intracellular RNA organization an explicit prediction target. Its datasets, tasks, and complementary metrics provide a testbed for developing spatially informed models, testing their ability to capture changes in abundance and localization, and identifying which aspects of cellular responses remain unresolved. This supports progress toward virtual cell representations that capture both molecular composition and internal organization.

## Ethics statement

SubcellPert reuses released imaging datasets without new human or animal experiments or donor re-identification. Source publications and accessions document provenance. Its population-level predictions and descriptive associations support methodological evaluation, not patient-level treatment effects or causal claims; interpretation depends on sampling, segmentation, and compartment geometry.

## Reproducibility statement

Appendices B and C document releases, curation, and reconstruction; Appendices A and E specify inputs, splits, models, and training. Appendix F defines metrics and aggregation. Further appendices give within-dataset analysis settings, support rules, and limitations. Split variation measures partition variability, not independent biological replication.

## Ai Use Statement

Generative AI tools assisted grammar checking and dataset auditing.

## Appendix guide

The appendices follow the order in which they are first referenced in the main text.

- Appendix A: Prediction inputs, target-access rules, representations, and frozen OOD splits.
- Appendix B: Dataset search, selection, molecule reconstruction, and eligibility.
- Appendix C: Individual dataset descriptions, native measurements, experimental designs, and interpretation limits.
- Appendix D: Observed RNA illustrations and their selection procedure.
- Appendix E: Baselines, pretrained resources, shared adapter, optimization, and checkpoint selection.
- Appendix F: Evaluation protocol, metric definitions, gene support, and aggregation.
- Appendix G: Within-dataset analyses of abundance–spatial associations, condition decoding, spatial patterns, population composition, and compartment geometry, followed by shared transformations, context transfer, recorded injury stages, and the auxiliary localization-only screen. Each subsection gives the settings and results of one analysis.
- Appendix H: Complete scores and target-gene coverage for all eight baselines, and gene-panel sensitivity supporting the benchmark experiments.

## A Task Definition, Representation, and Splits

### A.1 Prediction inputs and target access

Each observation is a complete gene-identified RNA point cloud. Both settings use the full assay gene axis and all RNA within selected cells. Oracle receives exact target cell-by-gene counts but not target geometry. Joint receives source observations, the condition descriptor, and the output-cell budget *B* = min(2000*, N_t_*), without target counts or geometry. Target-derived coordinate normalization is unavailable to both models.

The same target-cell selection is fixed before generation for both settings. Oracle samples one source cell uniformly from the permitted source pool for each supplied target count vector. Joint allocates its *B* source-cell draws as equally as possible across source units and samples with replacement within units. When *B* is smaller than the number of units, a fixed seed selects the units. These assignments are computational samples, not matched biological cell pairs; repeated source cells are not additional biological replicates.

Each generated cell uses one sampled source cell for all genes. A gene absent from that source cell uses its all-RNA coordinate distribution as a prior. Joint uses the same source cell for abundance and spatial prediction, and empty outputs remain zero-count cells without fabricated coordinates. All predictions are fixed before held-out observations are used for scoring.

### A.2 Coordinates and complete-cell spatial features

Let **p***_ik_* be the physical coordinates of the *M_i_ >* 0 eligible RNA molecules of observed cell *i*, pooled over genes. We normalize coordinates by

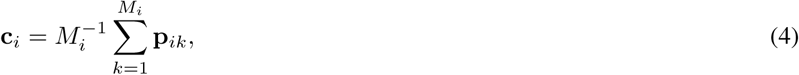

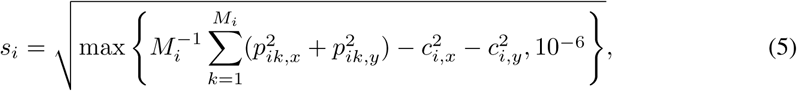

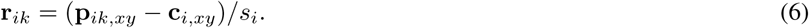

Thus the scale has a 10^−3^ floor in the source coordinate units. It uses the all-RNA centroid and RMS radius, not the segmentation centroid, a nuclear boundary, or a gene-specific scale. Each observed cell defines its own frame; there is no training-population fit of this geometric transformation. Only *xy* coordinates enter the model, including for releases with measured *z*. Target centers and scales are not supplied to either prediction interface. Cells with no eligible RNA have no defined frame and are excluded before population indexing. By contrast, a Joint model’s empty generated cell is retained as an all-zero count output, without fabricated coordinates.

For **r** = (*x, y*), define directions *D* = ((1, 0), (0, 1), (2^−1^*^/^*^2^, 2^−1^*^/^*^2^), (2^−1^*^/^*^2^, −2^−1^*^/^*^2^)) and frequencies *F* = (1, 2, 4). The fixed RNA feature map is

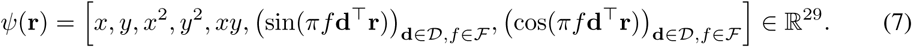

Direction is the outer index and frequency the inner index; all 12 sine terms precede all 12 cosine terms. For a gene with *n_ig_ >* 0, its spatial vector is 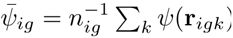; for *n_ig_* = 0 it is the zero vector, including the cosine slots. Concatenating [log(1 + *n_ig_*)*, ψ̅_ig_*] produces a 30*G*-dimensional feature vector. The training objective separates each gene’s 29 spatial entries from its log-count entry and compares the spatial entries directly; Joint separately supervises predicted log-counts using Equation (10). Every eligible molecule contributes. This fixed loss feature map is distinct from the learned point MLP used by the adapter.

### A.3 Condition descriptors

For cellular-context OOD, each known condition, including the reference, receives a one-hot vector in lexicographic order. The basis is shared across cell types, so a held-out context does not introduce an unseen condition token.

For perturbation OOD, PF-Astro represents each of 127 perturbation genes with a 127-dimensional Gene Ontology (GO) similarity vector and a 485-dimensional vector of source abundance correlations. GO similarities are Jaccard overlaps of the annotation sets used by GEARS; self-edges are zero, and up to 20 strictly positive neighbors are retained. These abundance correlations are computed over all 2,036 pooled control cells using Pearson correlation after library-size scaling to 10,000 and `log1p`. Absolute correlations are used for this descriptor, self-correlations are removed, and at most 20 neighbors with correlation at least 0.1 are retained. An unmeasured perturbation gene has a zero abundance-correlation block. Concatenating the two blocks and L2-normalizing gives 612 dimensions; the control descriptor is zero. No target-response abundance enters this construction. This descriptor is distinct from the adapter’s gene graph based on positive abundance correlations.

Table 6 lists the two-component multi-hot vectors used by ATM-RT an_√_d <u>D</u>dx54-PD1. The double factor is the sum of the two unit vectors, (1, 1); it is not divided by 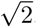. The source condition specifies each task’s direction. Held-out response abundance and geometry are not used to construct descriptors.

### A.4 Generalization splits

Twelve datasets use three split seeds and ATM-RT and Ddx54-PD1 each use one fixed split, giving 38 dataset–split instances. All methods and both settings share these partitions. One model learns the admitted training transitions within a dataset family and evaluates its held-out responses separately.

#### Response eligibility

A transition is admitted only when source and target each contain at least 50 cells in the same context and anatomical reference. This rule is applied before splitting and depends on population support, not spatial effect sizes or model performance.

#### Cellular-context OOD

Each split holds out a supported context label while retaining known response identities in the development contexts. Source controls remain available in the held-out context, but its response cells are excluded from both training and validation. Development response cells are partitioned into training and validation sets; these are cell-level, not independent animal- or patient-level, partitions. Table 4 lists the held-out contexts, response-cell counts, and number of evaluated responses. A context can recur across split seeds; split variability is not biological replication.

**Table 4:** Response-cell partitions and held-out conditions or contexts. Source cells are excluded from these counts; responses denote distinct evaluation contrasts. Part 1/2.

| Dataset | Seed | Held-out target | Train | Validation | Test | Responses |
| --- | --- | --- | --- | --- | --- | --- |
| PF-Astro | 1 | 18 genes | 7,215 | 777 | 1,426 | 18 |
| PF-Astro | 2 | 18 genes | 7,427 | 577 | 1,414 | 18 |
| PF-Astro | 3 | 18 genes | 6,090 | 542 | 2,786 | 18 |
| ATM-RT | 1 | double factor | 477,376 | – | 44,937 | 1 |
| Ddx54-PD1 | 1 | double factor | 739,964 | – | 216,519 | 1 |
| Lung-Drug | 1 | Lymphatic EC | 571,174 | 142,794 | 17,775 | 2 |
| Lung-Drug | 2 | Myeloid | 530,416 | 132,604 | 68,723 | 2 |
| Lung-Drug | 3 | Submucosal Gland | 584,675 | 146,169 | 899 | 2 |
| Sickness | 1 | e-M8 | 1,044,317 | 261,080 | 2,452 | 5 |
| Sickness | 2 | e-N10 | 1,044,899 | 261,225 | 1,725 | 5 |
| Sickness | 3 | e-C6 | 1,040,269 | 260,068 | 7,512 | 5 |
| DSS-Colitis | 1 | ICC | 957,696 | 239,424 | 2,972 | 4 |
| DSS-Colitis | 2 | Adipose | 959,537 | 239,885 | 670 | 3 |
| DSS-Colitis | 3 | EntericNervous | 941,971 | 235,493 | 22,628 | 4 |
| Adipose | 1 | Myeloid | 6,146 | 1,537 | 589 | 1 |
| Adipose | 2 | APC | 5,559 | 1,390 | 1,323 | 1 |
| Adipose | 3 | Endothelial | 3,996 | 1,000 | 3,276 | 1 |
| UC-VDZ | 1 | 04-IEC_CT_colonocyte | 18,675 | 4,669 | 1,812 | 2 |
| UC-VDZ | 2 | 05B-MNP-M2_<br>macrophage | 20,077 | 5,020 | 59 | 1 |
| UC-VDZ | 3 | 05G-MNP_activated | 20,084 | 5,021 | 51 | 1 |

**Table 5:**
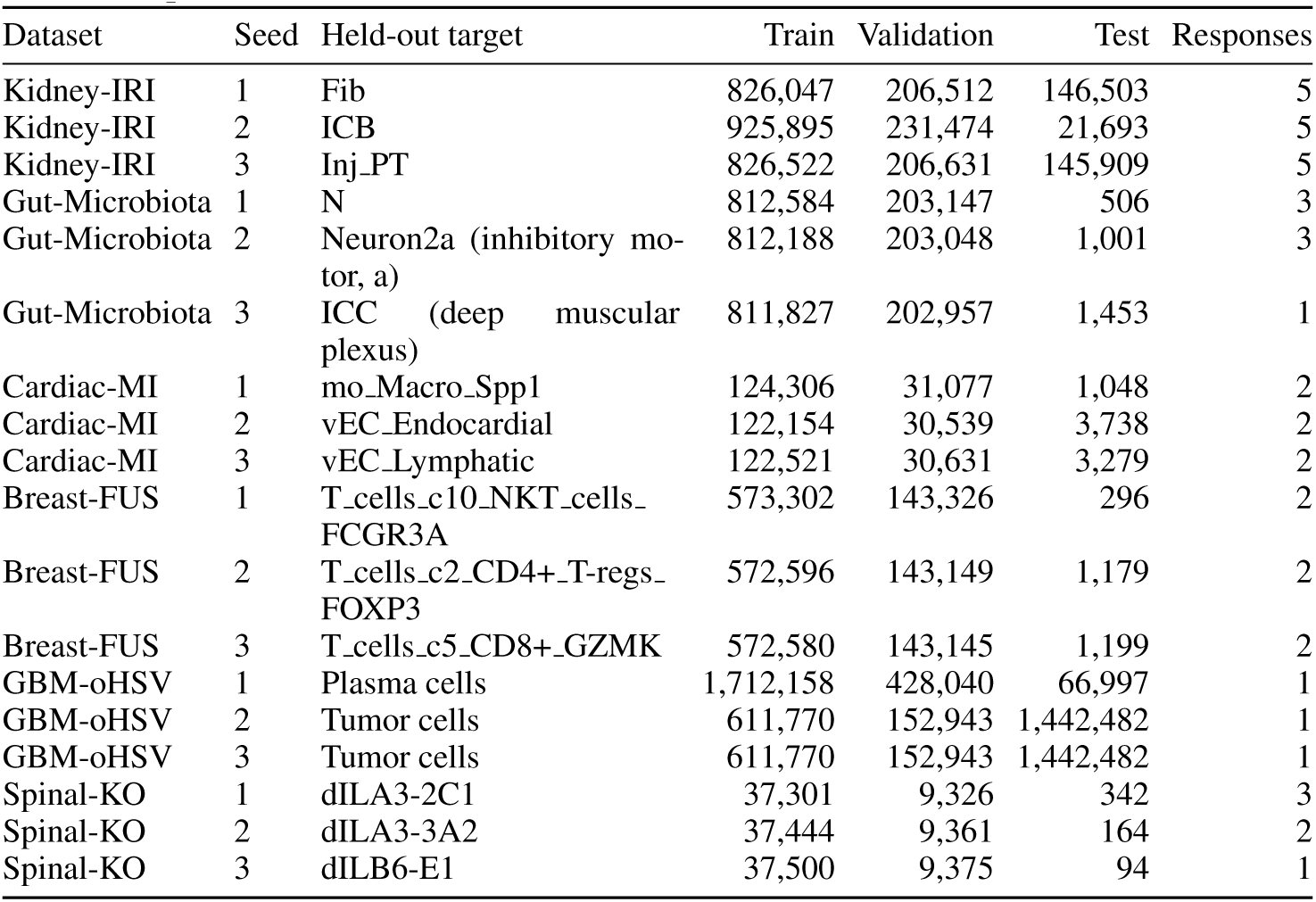
Response-cell partitions and held-out conditions or contexts. Source cells are excluded from these counts; responses denote distinct evaluation contrasts. Part 2/2.

#### PF-Astro perturbation holdouts

Of 127 stored perturbation identities, 87 satisfy response eligibility. For each seed, the first split assigns ⌊0.8 × 87⌋ = 69 identities to development and 18 to testing; the second assigns ⌊0.9 × 69⌋ = 62 to training and seven to validation. All response cells follow their perturbation’s partition. Controls remain permitted sources throughout. The descriptor space covers all 127 identities, but descriptor availability alone does not make a response eligible.

#### Compositional holdouts

For ATM-RT and Ddx54-PD1, Table 6 shows the single-factor training responses and double-factor test response, all relative to the untreated source. They have no validation partition and use the final epoch. The combined descriptor is available without observing its response cells; these tasks test composition, not an unseen intervention component.

**Table 6:** Registered factorial condition grids. Reference cells remain inputs; single-factor states are training responses; the double-factor state is the test response. The last column is the exact two-component descriptor.

| Dataset | Condition | Role | Descriptor |
| --- | --- | --- | --- |
| ATM-RT | ATM_intact | Reference | (0, 0) |
|  | ATM_lost | Train | (1, 0) |
|  | ATM_intact+RT | Train | (0, 1) |
|  | ATM_lost+RT | Test | (1, 1) |
| Ddx54-PD1 | WT+isotype | Reference | (0, 0) |
|  | Ddx54KD+isotype | Train | (1, 0) |
|  | WT+anti-PD1 | Train | (0, 1) |
|  | Ddx54KD+anti-PD1 | Test | (1, 1) |

## B Dataset Curation and Reconstruction

### B.1 Search strategy and eligibility

#### Search and release-level screening

We searched OpenAlex and Europe PMC using spatial-measurement terms, including MERFISH, seqFISH, Xenium, and CosMx, together with intervention or condition terms such as genetic perturbation, drug treatment, irradiation, immune challenge, injury, and recovery. We supplemented these results through publications, data-availability statements, and repository links. Candidate releases were traced to GEO, Dryad, Figshare, Brain Image Library, or author repositories and checked against their molecule tables, cell annotations, counts, and assignment files.

Bibliographic retrieval yielded 2,926 records and 2,514 entities after DOI/title reconciliation. Manual screening and supplementation produced 186 release branches from 184 source-work families. Publication records and release branches are different counting units: one publication can contain several assays or releases. The dashed retrieval links in Figure 2 therefore do not represent a single count-conserving sequence of datasets.

#### Eligibility

Admission required gene-resolved RNA coordinates, recoverable molecule-to-cell assignments, explicit conditions and context labels, auditable counts, compatible gene identifiers, and a supported source–target comparison for the selected OOD task. Count matrices or cell-centroid coordinates alone were insufficient. Cellular-context OOD additionally required comparable context labels across conditions; the two factorial Perturbation-OOD datasets retain explicitly identified mixed `all_cells` populations. Nuclear boundaries were not mandatory because the primary task predicts RNA point clouds. Requirements were checked for each release rather than inferred from the publication title.

#### Availability and exclusions

Of the 186 branches, 122 had usable access to public related objects and 64 did not. The latter comprise 40 branches whose corresponding dataset was not located and 24 without a usable open release. These 24 include CNP0006277, whose downloads required an access request, and syn51449054, which returned HTTP 403 with `GEO_RESTRICTION` from the screening environment. An access restriction does not establish that no public deposit exists.

Of the 122 accessible branches, 14 were retained and 108 excluded: 56 lacked suitable native molecule data, 16 lacked a resolved reconstruction or identity mapping, 21 lacked comparable cross-condition context labels, ten lacked response or split support, four were outside scope, and one lacked compatible gene identifiers. Each branch is counted once by its decisive exclusion reason. These decisions describe benchmark eligibility, not absence of biological response or an exhaustive inventory of all spatial datasets.

### B.2 Dataset diversity

The retained corpus contains 14 datasets, 13,880,723 nonempty cells, and 3,653,986,724 RNA observations. These are released population totals after cellular-context annotation and minimum-support filtering; they are not the sum of repeated train/validation/test views. The assays are MERFISH, Xenium, and CosMx, with retained gene panels ranging from 298 to 5,006 genes. Table 2 reports each population and panel. Panel size, molecule yield, segmentation, and label granularity vary substantially, so the common prediction interface does not imply that measurements from different datasets form one interchangeable biological population.

The datasets span eight organ/tissue categories. Cultured cells and tumor models are assigned by tissue of origin, and intestinal regions are grouped together. These are descriptive labels: for example, a lung-derived tumor model need not be a measurement of normal lung tissue. Perturbation categories overlap, giving 18 memberships across seven modalities. Biologics include antibodies, immune stimuli, and oncolytic virus; surgery includes bariatric-surgery-associated weight loss and ischemic injury. Category memberships count forms of experimental coverage, not independent interventions or biological replicates.

There are also distinct experimental designs within this collection. PF-Astro provides many single-gene perturbation identities in one cellular context. ATM-RT and Ddx54-PD1 provide one held-out combination each in complete 2 × 2 designs. The other eleven datasets support transfer of known condition responses to held-out cellular contexts, including animal challenges, injury stages, environmental comparisons, and clinical pre/post observations. A context may be a broad cell type, a finer subtype, an injury-associated state, or an unresolved author label; these are not a harmonized cross-dataset ontology.

The amount of RNA is also distinct from the amount of biological replication. Millions of cells can come from few animals, sections, or pooled clinical samples. We retain patient or animal identities where documented, distinguish slices and fields of view from those identities, and leave unverified mappings unresolved. The benchmark evaluates each response within its own dataset and panel. Its breadth tests models across different measurement and task settings; it does not establish a common cross-dataset biological effect.

### B.3 Molecule reconstruction and response eligibility

#### From native observations to complete cells

For each retained release, we preserve the link between a molecule’s gene, coordinates, cell assignment, sample, condition, and available context label. Cell identifiers are interpreted within their native sample, run, field of view, or region of interest before joining annotations; repeated local barcodes are not treated as the same cell. When explicit assignments are available, they provide the starting point. Where assignment must be reconstructed from released masks, the coordinate transform and mask-label mapping must also be established. We do not infer cell identities by matching an abundance profile to an unrelated assay or create molecule locations from a cell centroid.

Filtering follows the documented source-specific RNA and cell eligibility rules. These include endogenous-gene/probe selection, molecule quality, author cell-QC flags, and, for PF-Astro, the single-guide requirement. A targeted imaging panel is not subjected automatically to sequencing-specific mitochondrial or highly-variable-gene filters. The retained assay gene axis is preserved, including genes with zero count in an individual cell. After these filters, cells without any eligible RNA are excluded from the observed population. This final nonempty-cell rule is separate from upstream scientific selection and from later response-support checks.

Where author count objects are available, reconstructed cell–gene counts are compared on the corresponding gene axis and cell identities. Exact agreement supports the recorded assignments and filtering convention; it does not prove that segmentation is biologically perfect. We also distinguish a full matrix comparison from a comparison on the published columns available for cell matching, as in PF-Astro. The released cell-major representation stores complete eligible RNA point clouds and their sparse count vectors, rather than selecting a few molecules from a pooled cell population. Structural checks require consistent cell and gene pointers, finite coordinates, valid gene indices, and agreement between RNA payload lengths and summed counts. Reconstruction, structural validation, and task/split validation are separate checks.

#### Population, task, and split denominators

The upstream experiment, reconstructed source population, released population, and admitted prediction task need not contain the same cells. Before release, we remove context–condition groups with fewer than 50 cells, aggregating eligible units within each dataset using the benchmark’s cellular-context labels. This removes 674 groups containing 15,103 cells. UC-VDZ additionally excludes 411 cells without a matched cellular-context annotation; patient identities remain separate metadata. Together these filters remove 15,514 cells and 4,230,351 RNA observations from the reconstructed source populations. The totals in Table 2 describe the resulting release.

Adipose excludes Lean samples from its directed Obese-to-Weightloss population. PF-Astro first removes multi-guide cells, then removes 40 under-supported perturbations before release, leaving 11,454 cells across 87 perturbations and the control condition. For Lung-Drug, the released population retains untreated controls, but the evaluated task uses two drug add-on transitions from Bleomycin-treated sources. A task denominator therefore need not equal the released population size.

A prediction response additionally requires at least 50 source and 50 target cells in the same context, anatomical reference where applicable, and directed condition transition. This response check precedes the train/validation/test response partition and aggregates cells over eligible units. Neither support rule requires 50 cells in every individual animal or subsequent validation subset. A supported cell count is an engineering requirement for distribution evaluation, not a minimum number of independent biological replicates. A released target condition can meet the population threshold while lacking a supported matched source, so response eligibility remains a separate check.

Auxiliary control-identity examples used for model fitting can include source groups below the 50-cell response threshold; those groups are outside the filtered publication population.

Source observations may be reused for more than one compatible response or split. Such reuse does not create new cells, new donors, or additional independent tests. Task views retain their cell identities so that response cells assigned to validation or test can be checked against training membership. The detailed source-access and OOD rules are in Appendix A; the dataset descriptions below explain why those rules have different biological meanings across datasets.

#### Coordinates, boundaries, and compartment metadata

The common benchmark representation uses two-dimensional RNA coordinates, centered and scaled separately for each cell using its eligible RNA. Stored centers and scales permit reconstruction of native XY coordinates up to the released numerical precision. This RNA-based coordinate frame is not a membrane or nuclear boundary, and the compact XY payload is not a complete copy of every native image, z coordinate, or segmentation file. Normalization also changes the question from absolute tissue position to organization within the observed cell.

Native compartment information must therefore be checked separately from benchmark admission. Several retained Xenium releases provide nuclear-overlap annotations, cell/nuclear areas, or boundary files, and a CosMx molecule table can provide compartment labels. Their presence does not by itself establish that the labels and polygons match the cells, assignment version, and coordinate frame used in the benchmark. Kidney-IRI is a particularly clear example: Xenium segmentation objects cannot automatically be attached to the retained Baysor-assigned cells. A nuclear fraction needs a valid molecule-level compartment assignment; area-normalized enrichment additionally needs matched cell and nuclear areas; a boundary-distance statistic requires registered geometry. These requirements are not interchangeable. Missing fields in the compact benchmark are not evidence that the original release lacks them, and a public file listing is weaker evidence than inspection and registration of its contents.

## C Dataset descriptions

For each dataset, we describe the original biological design, the observations and annotations used in reconstruction, and the specific prediction question retained in the benchmark. Table 2 gives population sizes and panel sizes; those totals should be read together with the population/task distinction in Appendix B.3. Donor or animal identities recorded in each dataset are distinguished from acquisition groups and computational storage groups below.

### C.1 PF-ASTRO

Perturb-FISH combines MERFISH with pooled CRISPRi in human pluripotent-stem-cell-derived astrocytes (Binan et al., 2025). Its astrocyte release is available through the Brain Image Library. The material includes decoded RNA coordinates, mosaic-placement transforms, segmentation masks, perturbation assignments, and published cell count vectors. The retained observation panel has 485 genes. The single biological context is Astrocyte; different wells are not additional cell types or independent human donors.

Reconstruction places decoded observations into the released mosaic and uses the masks to recover cell assignment. The recorded identity audit matched all 14,926 published cell rows on the 277 count columns used for matching. This identity check has a narrower gene axis than the 485-gene observation panel. The benchmark then retains exactly one decoded guide copy per cell, leaving 12,748 cells before publication support filtering from wells sample1, sample2, and sample5. The 2,178 excluded cells with multiple guide copies are not treated as designed combinations. Non-targeting control subtypes are pooled within wells.

Of 127 reconstructed single-gene perturbation identities, 40 fall below the 50-cell threshold and are excluded before release. The released population contains 87 perturbations plus control, with 9,418 target cells and 2,036 controls. Each of the three frozen splits assigns 62/7/18 perturbation identities to training/validation/test. Thus the held-out variable is the targeted gene, while the cell context is known. Neither well membership nor the unpaired source and response populations provides an observed trajectory for an individual cell.

### C.2 ATM-RT

The Xenium dataset of tumoral ATM loss and focal radiation in a mouse diffuse midline glioma model (Mangoli et al., 2025) (GSE246584) provides four spatial samples in a complete 2 × 2 design: ATM intact, ATM lost, ATM intact with radiation, and ATM lost with radiation. The condition refers to the tumor model and exposure. It does not label every surrounding stromal or immune cell as having undergone a cell-autonomous ATM knockout.

Quality-filtered gene-targeting molecules are joined through sample-local cell identifiers. The reconstructed population contains 790,166 nonempty cells over 298 genes; the final nonempty-cell check removes 208 eligible annotated cells without retained RNA. The current task uses the label `all_cells`, which contains tumor and non-tumor cells. It is a mixed population, not an independently annotated homogeneous cell type.

The untreated ATM-intact state supplies the source. Training observes the two single-intervention responses; testing withholds their combined response. There is one compositional holdout and no separate validation split. The retained files give one spatial sample per factorial condition, so the large cell count does not provide replicated estimates across animals for that condition. This task examines prediction of a combined population response, which may include changes in cell composition as well as within-cell RNA organization.

### C.3 DDX54-PD1

The Xenium dataset of Ddx54 knockdown and anti-PD-1 in an LLC1 syngeneic mouse lung-cancer model (Gong et al., 2025) (GSE285341) also supplies a complete 2 × 2 design. The four released spatial blocks represent control or Ddx54-knockdown tumor cells combined with isotype or anti-PD-1 treatment. The local condition name WT denotes the control tumor arm, not a declaration that all cells in the section form one wild-type cell population.

Released assignments/masks and the physical-to-pixel transform establish the RNA-to-cell mapping and recorded count reconstruction. Removing 1,154 annotated cells without eligible RNA leaves 1,155,233 cells on the 5,006-gene panel. As in ATM-RT, the benchmark uses `all_cells`: tumor, immune, and stromal contributions are not separated into author-defined contexts in the active task. Native boundary files are additional source objects; their availability is separate from verification of the compact RNA representation.

Training uses the two single-intervention responses and testing uses Ddx54 knockdown plus anti-PD-1. This is generalization to a combination of seen components, not to an unseen single-gene perturbation. It has one test combination and no independent validation split. The four block identities do not establish a recoverable animal-level replication structure for each arm. Changes in immune infiltration or other population composition can therefore contribute to the observed response alongside intracellular changes.

### C.4 Lung-Drug

This Xenium lung dataset (Ziehr et al., 2026) (GSE325253) provides eight sections spanning untreated control, Bleomycin, Bleomycin plus AZD3965, and Bleomycin plus VB124. The author level-2 annotation has eleven contexts, including epithelial, vascular, fibroblast, immune, and other stromal populations. Assigned, quality-filtered molecules are joined to those labels by sample-local cell identifiers; the retained panel has 5,006 genes.

The current benchmark fits the two drug add-on transitions jointly, using Bleomycin-treated cells as source and the two combination arms as responses. Untreated controls remain in the reconstructed population but are not used to define a separate injury-prediction task in the reported benchmark. The reported response and model counts therefore cover the drug add-on task.

Each split withholds a cellular context while both known drug transitions are observed in other contexts. The test concerns context transfer, not unseen-drug or unseen-animal prediction. There are two section suffixes per condition, but the compact groups named replicate1 and replicate2 pool section identities and do not establish longitudinal pairing of the same animal across conditions. Native transcript, cell, and boundary files provide further information only after their sample and coordinate identities have been reconciled.

### C.5 Sickness

The Sickness MERFISH dataset (Sullivan et al., 2025; Moffitt et al., 2025) profiles the mouse hypothalamus under health and induced sickness states; its molecule release is available at Dryad. The reconstructed cohort has 16 animals and 105 source context labels, including neuronal subtypes and non-neuronal populations. The known challenges are DSS, LPS, PIC, PLA, and STAG, with PBS as reference. These labels describe immune or sickness-inducing conditions, not five genetic knockouts.

Animal-prefixed cell identities align molecule observations and cell annotations. Known nonblank genes with mir ≤1.0 reproduce the recorded raw-count matrices, and cells marked for exclusion by the authors are removed. The resulting panel contains 941 genes. Population support filtering leaves 1,765,443 cells and 103 contexts. Large differences in cell abundance across hypothalamic populations make context-specific source and response support more informative than the total number of measured cells.

The task learns the five known challenge responses together and withholds a cellular context in each split. PBS sources must belong to the same context; unsupported comparisons such as PLA/Monocytes are not evaluated. Animals are the biological groups, and challenge arms comprise different animals. The dataset therefore supports within-dataset comparison of observed condition populations and context transfer, without implying longitudinal tracking, causal separation of all systemic effects, or prediction of unseen challenges.

### C.6 Dss-Colitis

The DSS colitis MERFISH time course (Cadinu et al., 2024; Moffitt, 2024) is distributed through Dryad. The reconstructed population contains 58 cell-assigned slices from 17 recorded mouse groups and a 940-gene panel. Eight Tier-1 contexts represent adipose, endothelial, enteric nervous, epithelial, fibroblast, interstitial cells of Cajal, immune, and smooth-muscle populations. Here Adipose is a context label within the colon dataset, distinct from the Adipose clinical dataset below.

We preserve the released cell identities, Tier-1 annotations, and assigned RNA. The benchmark defines four D0-to-post-treatment transitions, at D3, D9, D21, and D35. A condition label identifies the sampled stage of one DSS exposure/recovery design; it does not denote a different drug at every day. The mice sampled at different stages do not form complete longitudinal pairs.

All supported transitions share a fit in each split, and the held-out variable is a cell context. Adipose has no D35 target observations, so that response is excluded without removing the entire dataset or relabeling missing cells. The task assesses transfer of known stages to a held-out context, not temporal extrapolation. D0 is the untreated reference; the design does not supply an untreated time-matched control at every later stage. Descriptions of stage changes should preserve this limitation and the distinction between slices and independent mice.

### C.7 Adipose

The human adipose dataset (Miranda et al., 2025) (GSE295862; SCP3117) includes obese, post-weight-loss, and lean observations. The benchmark selects the Obese-to-Weightloss comparison from four paired donors, yielding 13,972 released cells on a 477-gene panel after removing 35 under-supported source mast cells. The larger recovered source contains 25,795 cells; that upstream number includes observations outside the directed task. The seven level-1 contexts are adipose progenitor cells, adipocytes, endothelial, lymphoid, mast, mural, and myeloid cells.

Native event-to-cell assignments reconstruct the recorded counts, while sample identities preserve the donor pairing across the two conditions. Lean observations are not pooled into the source: they come from a different comparison and would change the interpretation of the response. The deposited cell and nuclear geometry is a separate source of compartment information whose registration is documented in Appendix G.5.

Each fit learns the known Obese-to-Weightloss transition from observed contexts and evaluates a held-out context, subject to the common response-support rule. The held-out context is not an unseen patient. Pairing enables within-donor comparisons, but the clinical transition combines surgery-associated weight loss, elapsed time, and other changes. It is not an isolated drug intervention or a measurement of the same individual cells before and after treatment.

### C.8 UC-VDZ

The ulcerative-colitis vedolizumab CosMx dataset (Kattah, 2024) has a versioned Figshare release. From a source with 19 patient identifiers, the benchmark retains seven complete longitudinal patients: four responders and three non-responders. Healthy and post-only patients are outside the directed paired task. The released population has 80,562 cells and a 1,000-gene panel, after removing 143 cells in under-supported context–condition groups and 411 cells without matched cellular-context annotations.

Molecule assignments, sample/field identities, and the deposited annotations are reconciled within patient. The source annotation has 29 author labels, including five labels containing `unassigned`; 28 labels remain after population support filtering. These labels should not be described as fully identified cell types. These labels are kept explicit rather than converted into newly inferred biological types. Patient is the biological grouping unit, whereas cell context is the OOD variable; storage grouping by patient must not be mistaken for a patient-holdout experiment.

Responder-specific and non-responder-specific pre/post transitions are fitted together, with supported responses in the held-out context evaluated separately. The response-support rule removes insufficiently populated comparisons rather than assuming that every one of the 29 labels supports both transitions. Crucially, response status is an eventual clinical outcome. Providing that status in a condition descriptor makes the task retrospective and outcome-conditioned, not a prospective prediction of whether a new patient will respond. Three complete non-responder pairs also give limited subgroup support.

### C.9 Kidney-IRI

The GSE269719 Xenium/Baysor release profiles kidney ischemia–reperfusion injury. Its twelve samples represent left and right kidneys from six mice and supply twenty annotated contexts on a 300-gene panel. The contexts include nephron segments, vascular and immune populations, and injury/repair-associated proximal-tubule states. A state label such as injured proximal tubule is an observed context, not a known one-to-one lineage match to a control cell.

Native assignments are checked against the author raw counts, and paired kidneys retain a shared mouse identity. The retained assignment is the Baysor assignment; other Xenium segmentation objects under the accession do not necessarily use the same cell definition. Additional cell or nuclear geometry must be registered to this assignment before it can support compartment measurements.

Day-14 Sham samples provide the reference for five post-injury stages: 4 hours, 12 hours, 2 days, 14 days, and 6 weeks. The Sham reference is not a day-0 baseline or an independently time-matched control for every stage. There is one mouse per stage, not two independent replicates obtained by counting kidneys separately. The benchmark transfers these known stage responses to held-out contexts; it does not establish temporal extrapolation or separate a stage effect from between-animal variability.

### C.10 Gut-Microbiota

The Gut-Microbiota MERFISH release compares specific-pathogen-free (SPF) and germ-free (GF) conditions in ileum, cecum, proximal colon, and distal colon. The source contains 105 slices in 28 acquisition groups and 78 context labels spanning epithelial, immune, stromal, vascular, and enteric nervous populations. Some labels further distinguish anatomical positions or cellular states. These are within-dataset annotations, not direct counterparts of every label in the other intestinal datasets.

Following the author preprocessing, we require quality score *>* 0.8 and assignment confidence *>* 0.5. The 106 genes removed by the source contamination filter are not restored, giving a retained panel of 1,815 genes. The recorded reconstruction matches the corresponding post-QC raw-count axis; the larger pre-filter gene axis describes a different stage of processing.

The filtered release contains 2,057,563 cells and 75 contexts. Four SPF-to-GF responses are defined within matching intestinal regions. Source and target must share both region and context, and sparse comparisons are excluded by the common support rule. The model learns known condition responses in other contexts; microbiota status and region are not held-out intervention identities. Acquisition groups and slices do not establish an independent-animal count. The comparison also does not represent immediate microbiota removal from the same tracked animals: developmental and environmental differences can contribute to the observed condition populations.

### C.11 Cardiac-mi

The GSE280376 Xenium release profiles myocardial injury and repair. Cardiac-MI is the retained benchmark identifier, but the deposited injury is cryoablation; the name should not be read as evidence of coronary-ligation-induced infarction. The admitted material comprises seven slides, a 475-gene panel, and 74 source context labels from the deposited clustering and cluster-name crosswalk; 225,023 cells and 67 contexts remain after publication support filtering.

Native cell identifiers link RNA observations to the author annotations. GSM9460460 is excluded because its deposited annotation duplicates another slide and cannot be reconciled to its native cells. This is an identity-mapping failure for that sample, not a reason to infer replacement cell types. For some other samples, GEO distributes the same cell table in both CSV and Parquet; these are alternative file encodings, not additional slides or replicates.

Day-7 Sham samples serve as source for day-7 and day-28 injury responses. Supported context-specific transitions share a fit, with a context withheld for evaluation. The day-28 comparison consequently uses the day-7 Sham reference. The design provides two injury stages rather than a densely sampled trajectory. Slides are acquisition groups unless an independent-animal mapping is documented, and neither the slide count nor the number of cells supplies that missing replication information.

### C.12 Breast-FUS

The GSE296517 CosMx release concerns breast-tumor observations under focused ultrasound (FUS), with or without gemcitabine (GEM), and untreated reference material. Four native runs contribute 880,565 released cells on a 1,000-gene panel after population support filtering. The deposited annotation supports 46 valid cellular contexts, alongside observations without a defined context.

The join uses run, field-of-view, and original cell identifiers. These keys resolve a swapped prefix in the merged annotation object; an apparently shared prefix alone would assign cells to the wrong source run. Cells labeled `undefined` do not define a prediction context and are removed from this release by the population support filter. Native CosMx compartment annotations are a separate input from the cell-type labels and require their own cell/molecule correspondence.

Pooled untreated sources support the known FUS and FUS+GEM responses in held-out contexts. A complete patient-to-field correspondence is unavailable for the retained mapping, so the clinical runs are conservatively pooled and storage groups are not counted as independent patients. The resulting task is neither patient-OOD nor a subtype-matched causal comparison. Because a separate, matched GEM-only arm is not supplied by this task, the combined response also does not identify an isolated gemcitabine interaction or synergy effect.

### C.13 GBM-oHSV

The GSE296577 Xenium release contains glioblastoma observations from eight patients, with eleven Tier-1 contexts and twenty native regions of interest (ROIs). After author QC and population support filtering, the release retains 2,401,940 cells and 520,974,197 RNA observations over 480 genes. One native ROI has no cells remaining after the final author QC; it does not contribute an additional nonempty population.

ROI-local identifiers are reconciled to the deposited annotations and raw-count assay. The recorded reconstruction gives unique cell matches and count agreement. Multiple ROIs belonging to one patient retain that patient’s identity, rather than being treated as additional independent donors. The compact point-cloud release does not, by itself, certify that every associated native boundary archive is complete or registered for compartment analysis.

The benchmark learns the known pre/post condition response and evaluates held-out cellular contexts. The observed transition combines sampling after oncolytic herpes simplex virus treatment with recurrence and elapsed time. It therefore describes a clinical condition change, not a treatment effect isolated from disease course. Patient pairing can organize within-patient comparisons, while the benchmark’s context holdout still leaves patient identities represented across the source and other observed contexts.

### C.14 Spinal-KO

The GSE291460 Xenium release examines E16.5 mouse spinal cord development under transcription-factor knockouts. It supplies four control and four knockout animals, a 347-gene panel, and 179 source context labels, including fine neuronal subtypes and developmental states. Removing 478 under-supported context–condition groups leaves 122,645 released cells and 158 contexts. This fine label resolution differs from the broad cell classes used by several other datasets.

Animal-specific namespaces precede barcode joins among molecule coordinates, cell-feature matrices, and the deposited coordinate-table labels. The reconstruction retains the final author-annotated cells and verifies count agreement, with control probes handled separately from the endogenous gene panel. Eight animals should not be counted as ten independent replicates by splitting anatomical sections.

The source reference is anatomically matched: brachial controls support Ptf1a, Zic3, and Zic14 knockout responses, whereas thoracic controls support Gsx1/2 knockout. Supported responses share a fit with a held-out cellular context; rare context–condition combinations are not rescued by pooling anatomically incompatible controls. The test uses known knockout identities in new contexts, not unseen knockouts or unseen animals. Four knockout animals across different conditions also do not provide four independent replicates of every individual knockout response.

## D Observed RNA Examples

Figures 5 to 7 show 56 observed cells across 14 datasets. Each row contains two source and two response cells from the seed-1 training population. We choose the context–transition pair with the largest minimum source/response cell count, then cells nearest each arm’s 25th and 75th RNA-count percentiles; ties are lexicographic. The three genes most abundant across the displayed source cells receive consistent colors across the row; other genes are gray. Every retained RNA is plotted, without rotation or subsampling.

**Figure 5:**
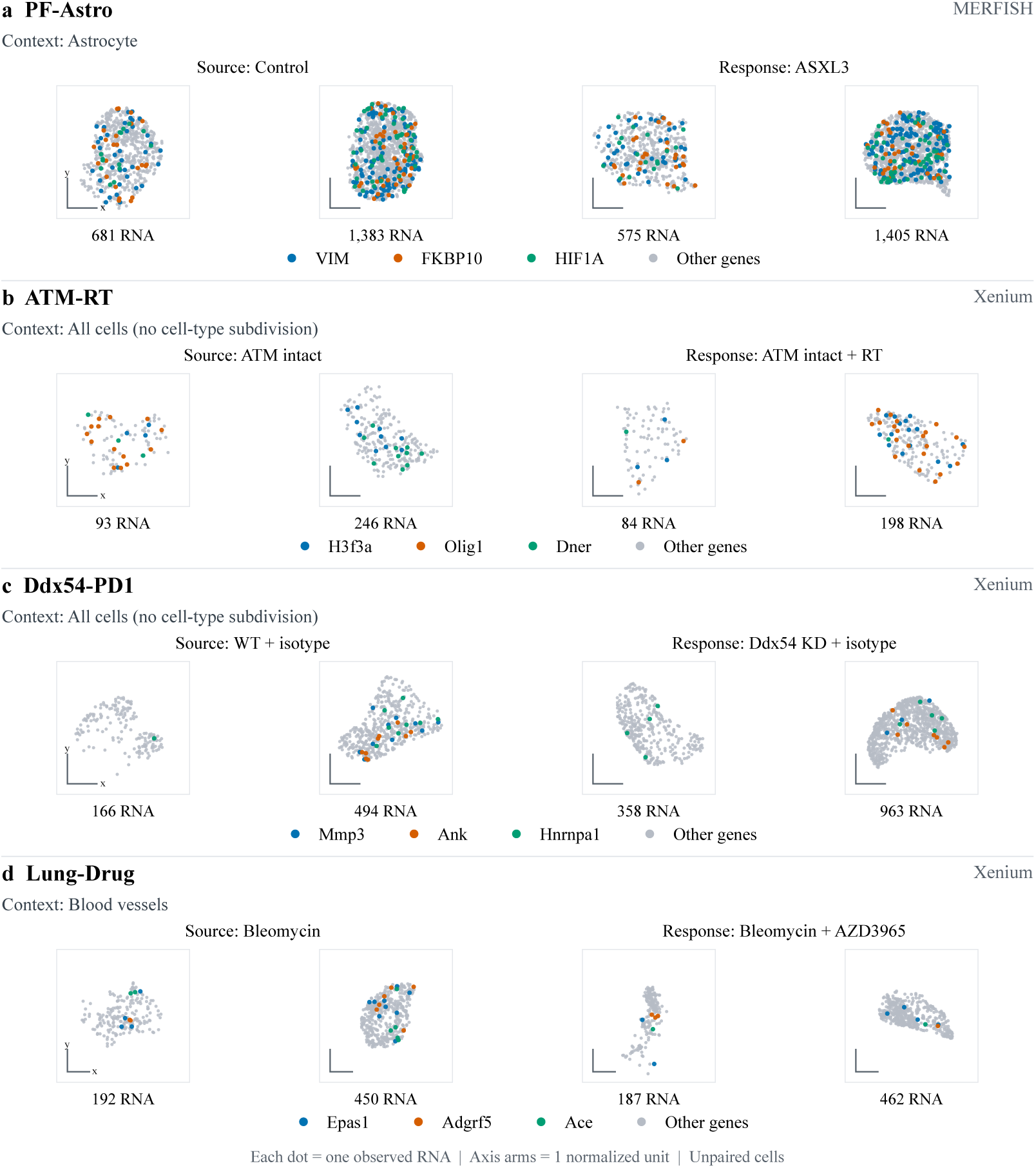
Observed RNA examples (1/3). PF-Astro, ATM-RT, Ddx54-PD1, and Lung-Drug. Each dot is one RNA molecule; counts below cells include all genes.

**Figure 6:**
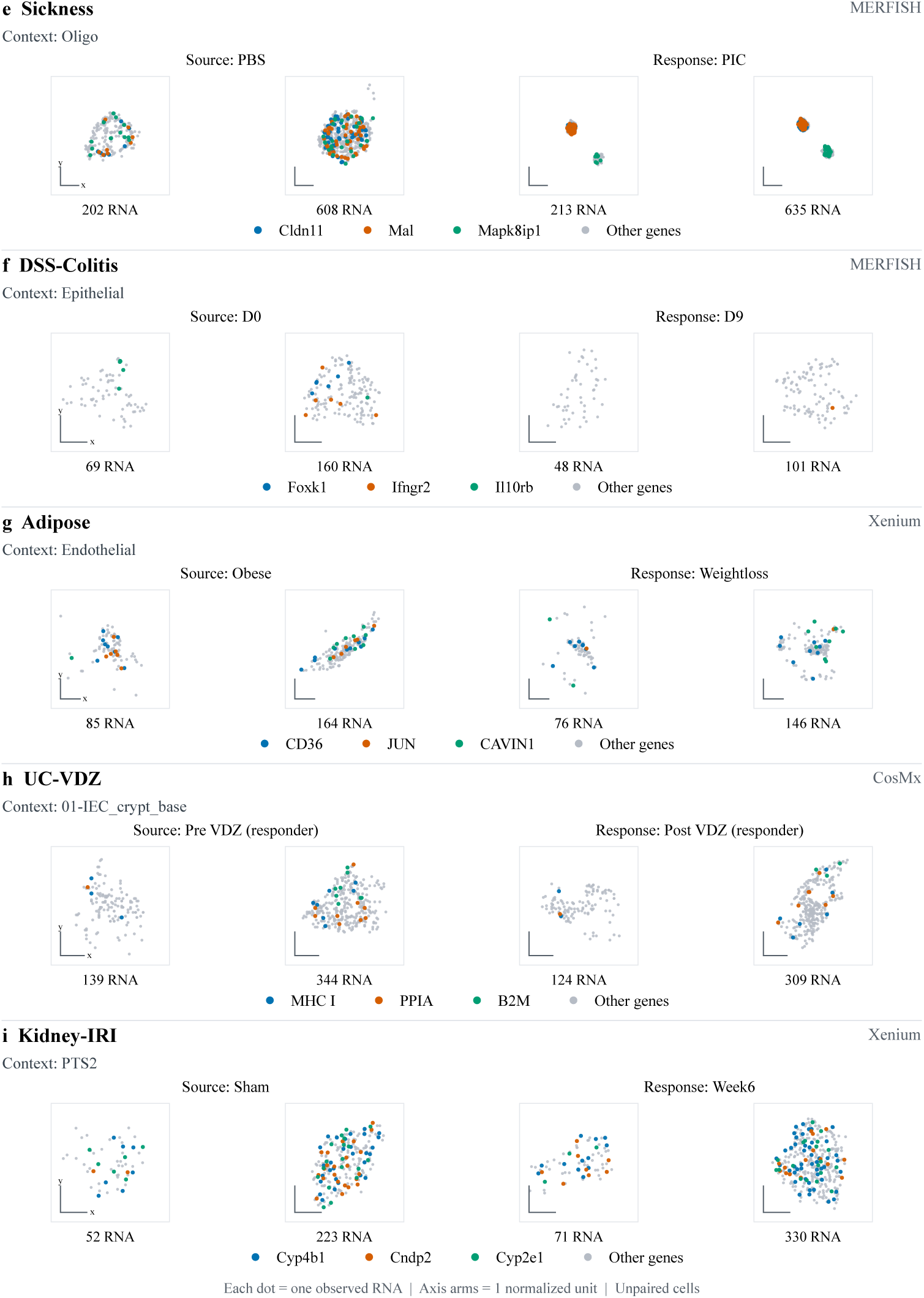
Observed RNA examples (2/3). Sickness, DSS-Colitis, Adipose, UC-VDZ, and Kidney-IRI, using the same selection and plotting conventions. UC-VDZ retains its retrospective responder condition label.

**Figure 7:**
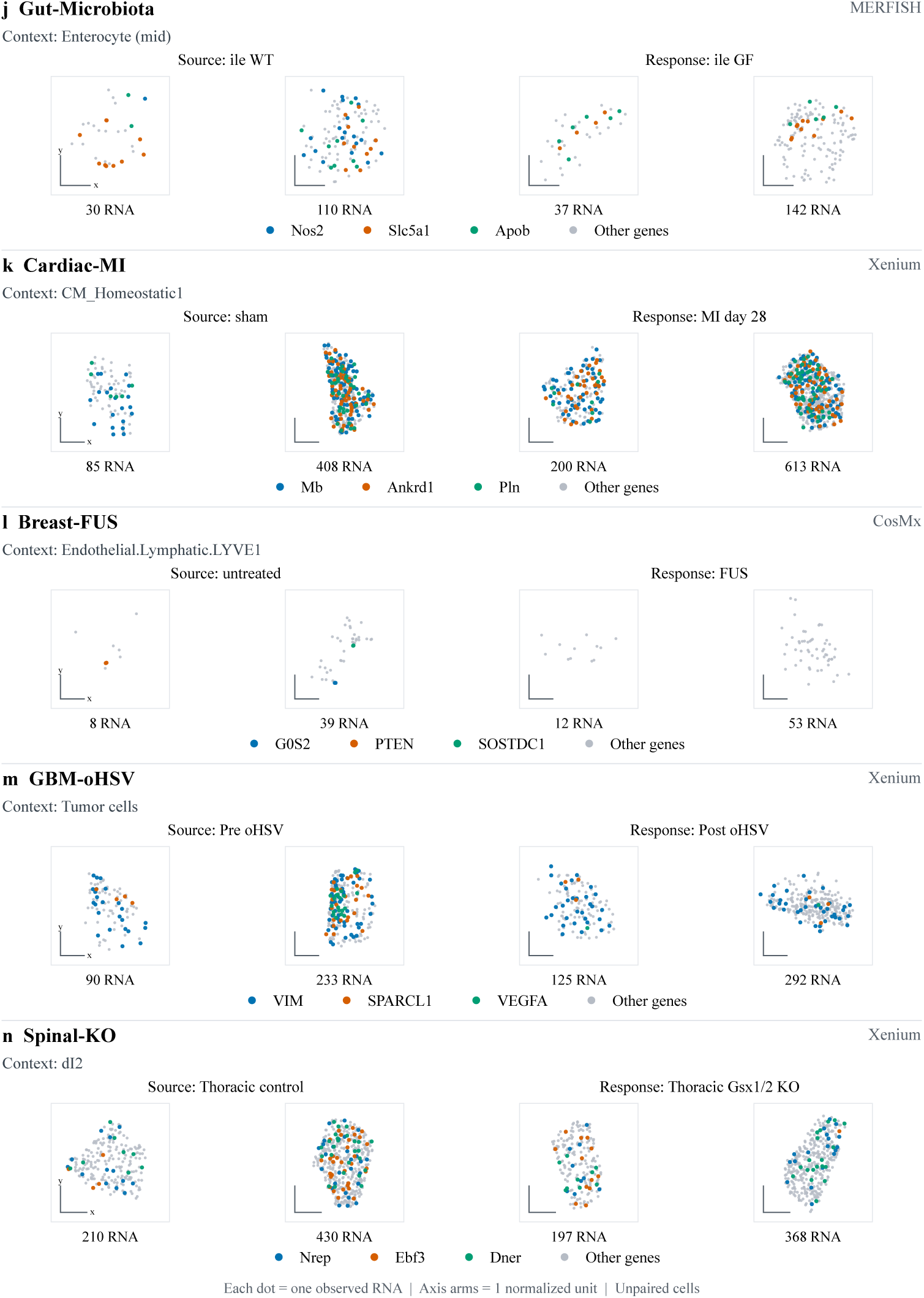
Observed RNA examples (3/3). Gut-Microbiota, Cardiac-MI, Breast-FUS, GBM-oHSV, and Spinal-KO. Source and response cells are independently selected, including for longitudinal datasets.

All cells in a row share axis limits in the normalized frame of Appendix A.2; each axis arm denotes one normalized unit. Frames are not cell membranes and do not preserve physical size or tissue position. These unpaired observations illustrate data structure, not model predictions or causal treatment effects.

## E Baselines and Training Protocol

### E.1 Source resampling and classical transport

Source resampling copies source-cell counts in Joint and resamples source-cell coordinates to satisfy the supplied counts in Oracle. Gaussian affine and radial quantile transport fit condition-dependent spatial regressions from complete-cell statistics, with ridge coefficient 10; radial maps use 16 empirical quantiles. Their Joint abundance regressions fit group-average log(1 + *n*) shifts, clipped to [−log 4, log 4], followed by nonnegative log-count flooring and integer rounding. Source-cell assignments are computational samples, not observed cell pairs.

### E.2 Pretrained encoders and gene alignment

All five encoders remain frozen and provide source-cell gene representations to the shared adapter. Table 7 summarizes their representation widths and token budgets. scGPT (Cui et al., 2024) and scGPT-spatial (Wang et al., 2025) use contextual gene-token states before cell pooling; SpatialFormer (Wang et al., 2026) uses its final transformer layer before gene-mean pooling. Token identities align features to assay genes, rather than positions in an abundance-ranked sequence. We use human-pretrained scGPT, scGPT-spatial v1, SpatialFormer’s step-176000 checkpoint, and the released scFoundation gene-mode weights.

**Table 7:** Pretrained gene representations. Width excludes the availability indicator; token budgets exclude special tokens. scFoundation uses matched assay genes at their official vocabulary positions.

| Encoder | Gene width | Gene-token budget |
| --- | --- | --- |
| scGPT-spatial | 512 | 1199 |
| SpatialFormer | 512 | 1200 |
| scGPT | 512 | 1199 |
| scFoundation | 512 | matched assay |
| STATE-SE | 2048 | 2047 |

For these encoders and scFoundation, alignment uses exact symbols followed by unambiguous case-insensitive matches, without orthology inference. scGPT variants retain positive-count matched genes, selecting the highest counts with deterministic ties when exceeding 1,199 genes plus CLS. SpatialFormer admits 1,200 gene tokens, five prefix tokens, and SEP. Zero-count, unmatched, and truncated genes have unavailable states but remain on the downstream assay gene axis.

scFoundation (Hao et al., 2024) uses gene-mode decoder states for matched assay genes at their official positions in the 19,264-gene vocabulary, including zero-count genes and two library-depth tokens. A cell without matched RNA has unavailable features. Per-cell stabilization of its Performer key kernel preserves the singleton computation independently of batch composition.

STATE-SE uses the SE-600M checkpoint and protein-derived gene inputs (Adduri et al., 2025). Human genes use exact symbols; mouse genes use unambiguous one-to-one MGI mouse–human homologs. Mapped genes are ranked by source count with symbol-based ties, retaining at most 2,047 unique genes plus CLS. Counts are transformed as 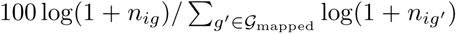, with zero for CLS. Matched zero-count genes may occupy remaining positions in a nonempty library; short panels are not padded with duplicate genes. Attention dropout is disabled and the native State Transition module is unused.

Source-derived normalization statistics are estimated only from unique training-source cells and fixed thereafter. scGPT-spatial supplements its reference means with source-derived nonzero and per-gene fallback means; quantile-bin ties are deterministic. SpatialFormer ranks counts divided by its pretrained gene medians, using one when a median is absent or invalid. Its species, tissue, and assay prefixes are listed in Table 9; these are encoder choices, not biological reannotations (for example, Human Skin is the adipose tissue proxy). Healthy/Disease prefixes depend only on the permitted source condition.

**Table 8:**
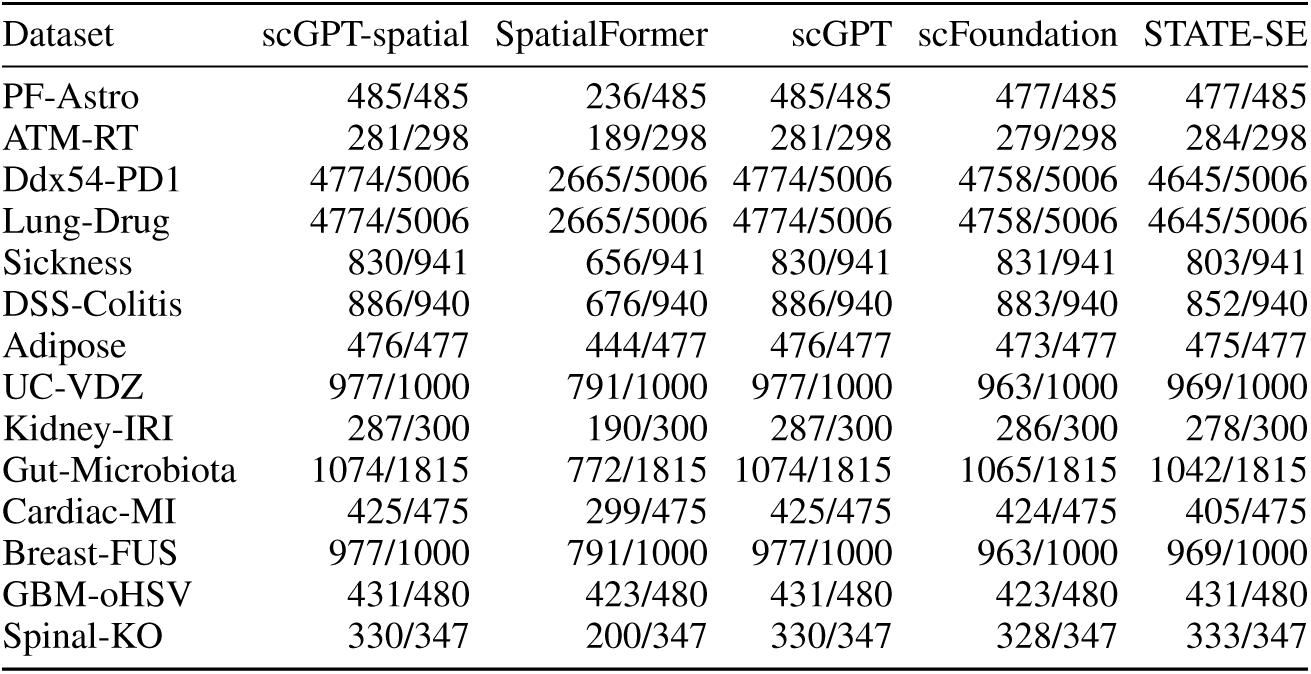
Assay-panel vocabulary coverage for all fourteen datasets and five encoders (matched/panel). The first four use exact then unambiguous case-insensitive matching; STATE-SE uses exact human symbols or one-to-one mouse–human homology. This is input vocabulary coverage, not per-cell token availability or Joint output-gene coverage.

| Dataset | scGPT-spatial | SpatialFormer | scGPT | scFoundation | STATE-SE |
| --- | --- | --- | --- | --- | --- |
| PF-Astro | 485/485 | 236/485 | 485/485 | 477/485 | 477/485 |
| ATM-RT | 281/298 | 189/298 | 281/298 | 279/298 | 284/298 |
| Ddx54-PD1 | 4774/5006 | 2665/5006 | 4774/5006 | 4758/5006 | 4645/5006 |
| Lung-Drug | 4774/5006 | 2665/5006 | 4774/5006 | 4758/5006 | 4645/5006 |
| Sickness | 830/941 | 656/941 | 830/941 | 831/941 | 803/941 |
| DSS-Colitis | 886/940 | 676/940 | 886/940 | 883/940 | 852/940 |
| Adipose | 476/477 | 444/477 | 476/477 | 473/477 | 475/477 |
| UC-VDZ | 977/1000 | 791/1000 | 977/1000 | 963/1000 | 969/1000 |
| Kidney-IRI | 287/300 | 190/300 | 287/300 | 286/300 | 278/300 |
| Gut-Microbiota | 1074/1815 | 772/1815 | 1074/1815 | 1065/1815 | 1042/1815 |
| Cardiac-MI | 425/475 | 299/475 | 425/475 | 424/475 | 405/475 |
| Breast-FUS | 977/1000 | 791/1000 | 977/1000 | 963/1000 | 969/1000 |
| GBM-oHSV | 431/480 | 423/480 | 431/480 | 423/480 | 431/480 |
| Spinal-KO | 330/347 | 200/347 | 330/347 | 328/347 | 333/347 |

**Table 9:** SpatialFormer species, tissue, and assay prefixes. Proxy tokens select available pretrained representations; they do not redefine dataset biology.

| Dataset | Species | Tissue token | Assay token |
| --- | --- | --- | --- |
| PF-Astro | Human | Brain | Merfish |
| ATM-RT | Mouse | Brain | Xenium |
| Ddx54-PD1 | Mouse | Lung | Xenium |
| Lung-Drug | Mouse | Lung | Xenium |
| Sickness | Mouse | Brain | Merfish |
| DSS-Colitis | Mouse | Colon | Merfish |
| Adipose | Human | Skin | Xenium |
| UC-VDZ | Human | Colon | CosMx |
| Kidney-IRI | Mouse | Kidney | Xenium |
| Gut-Microbiota | Mouse | Colon | Merfish |
| Cardiac-MI | Mouse | Heart | Xenium |
| Breast-FUS | Human | Breast | CosMx |
| GBM-oHSV | Human | Brain | Xenium |
| Spinal-KO | Mouse | Brain | Xenium |

Table 8 reports panel-to-vocabulary compatibility, which differs from cell-level token availability and from Joint output-gene coverage. An unavailable state is zero with an availability indicator of zero; all assay counts and RNA remain available to the other model branches.

### E.3 Shared GEARS adapter

The four 512-dimensional encoders and the 2,048-dimensional STATE-SE encoder each supply a gene-availability indicator. A learned projection maps their outputs to a common 512-dimensional space, combined with gene identity, source log-counts, and per-cell/gene RNA features from a 2 → 32 → 16 point MLP. The gene graph uses positive correlations between gene abundances in unique training-source cells. A condition graph uses cosine similarity between the descriptors in Appendix A.3, replacing the genetic-only interface of GEARS (Roohani et al., 2024). Both graphs retain up to 20 positive neighbors per node and use one graph-convolution layer. Held-out descriptors, but not their response observations, are available.

Recovery MLPs produce per-gene states and scalar readouts, followed by a *G* → 512→ 512 cross-gene MLP. The spatial head combines gene states, the cross-gene summary, scalar readouts, and conditioning log-counts through a 1026 → 512 → 6 MLP:

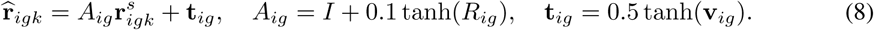

The transform is initialized to identity and applied to source cells sampled under the protocol in Appendix A.1.

Oracle has no abundance head. Joint adds a gene-specific readout conditioned on the source gene states and cross-gene summary:

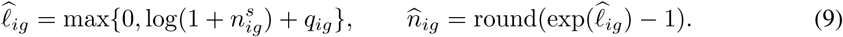

The count residual is unbounded; nonnegative flooring and rounding give absolute molecule counts, including zero. Spatial training conditions on observed development counts, whereas Joint inference uses predicted counts. The abundance head never receives target counts. Singleton training inputs use running BatchNorm statistics.

### E.4 Optimization and Model Selection

Each dataset–split–family model learns all admitted training transitions, with condition/context groups kept separate in the loss. Oracle and Joint are fitted independently; individual test responses do not trigger retraining.

#### Optimization and batch size

Oracle and Joint are fitted independently for five epochs with frozen foundation weights. The adapter and output heads use Adam (Kingma & Ba, 2015) with initial learning rate 5 ×10^−4^ and weight decay 5 ×10^−4^. StepLR halves the learning rate after each epoch. All neural runs use a fixed physical batch size of 256 and accumulation factor one. Gradient values are clipped to [−1, 1] after each backward pass. Frozen encoders compute features online from source cells. Their microbatch size is 256, except for STATE-SE on Lung-Drug and Ddx54-PD1, where it is 64 to reduce encoder memory use without changing the optimizer batch size. Experiments use NVIDIA A100 accelerators. Geometry sampling is seeded per generated cell, independently of inference batch partitioning.

#### Sampling and checkpoint selection

A fixed assignment seed of 42 draws one source cell with replacement per development target, supplemented by deduplicated control-identity examples. For a response example, the source cell and perturbation descriptor are the predictor inputs and the observed response cell supplies the supervision. In a control-identity example, the control cell itself supplies both the source and target. This follows the upstream construction of one random control input per perturbation-response cell, extended to each task’s permitted source pool. Each epoch globally shuffles complete-cell examples and drops an incomplete final microbatch. Source-cell assignments are computational choices, not biological cell pairs. Independent validation selects minimum all-gene spatial W1 for Oracle and minimum top-20-DEG abundance MSE for Joint, breaking ties in favor of the earliest epoch. Initialization (epoch 0) also participates in validation selection. Validation DEGs are ranked using validation source/target populations only; the top-20 restriction applies to checkpoint selection, while the training objective uses the condition-specific gene support defined below. Outer test responses never enter model selection. ATM-RT and Ddx54-PD1 have no validation split and use the final epoch.

#### Training objective

All five Joint foundation-model variants use the scFoundation/GEARS abundance loss (Roohani et al., 2024; Hao et al., 2024; Wei et al., 2026). For each condition/context group *c* represented in a microbatch, let *B_c_* be its computational training examples and *G_c_* contain genes with nonzero counts in at least one cell of that group’s training response population. Zero observations in individual cells are retained for these genes. Control-identity groups retain all genes. With raw-count log1p values *ℓ* and the mean allowed training-source log-count *ℓ̅^s,c^*, Joint’s abundance term is

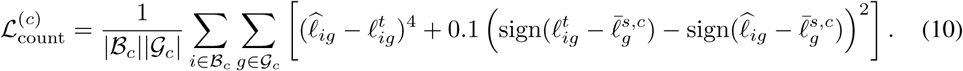

This matches the upstream fourth-power abundance error, direction-term coefficient passed by GEARS, and equal weighting of represented condition groups. Gene support and control means are computed from training data only. The direction term records disagreement in up- or down-regulation relative to the source mean. We retain the upstream hard sign operation and its zero autograd derivative; the abundance prediction receives its training gradient from the fourth-power error. Raw-count log1p, rather than library-normalized counts, permits generation of absolute RNA counts.

For spatial supervision, each cell/gene has 29 statistics computed from all its RNA coordinates: the moments and fixed Fourier features *ψ̅* defined in Appendix A.2. With the same groups and gene support, both modes use

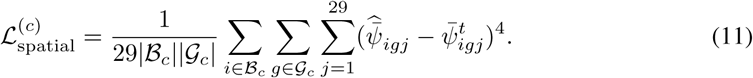

Let *C_B_* be the condition/context groups represented in the microbatch. The complete objectives are

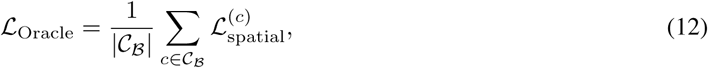

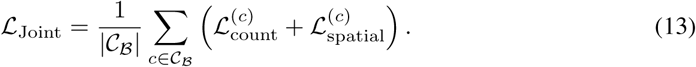

The spatial objective is our RNA-coordinate extension; scPerturBench does not supply this spatial task or loss. Oracle has no abundance prediction objective. Both modes condition spatial training on observed development counts. Joint uses predicted integer counts at inference, whereas Oracle continues to use the supplied counts. The source/target assignments remain random computational inputs, not biological cell correspondences.

The classical baselines are fitted analytically and do not use this neural optimization schedule. The fixed spatial training features differ from the 64 projection directions used by the evaluation metric.

### E.5 Trianing hardware and software

All neural training uses PyTorch 2.14 on A100 clusters. The overall training time is about 1,500 GPU hours.

## F Evaluation protocol and metrics

The evaluation asks three separate questions: are the RNA counts correct, are target genes represented, and are their predicted locations correct? Abundance metrics address the first question, coverage the second, and spatial W1 the third. Oracle receives exact target counts and is evaluated only for localization. Joint predicts both counts and coordinates, so all three questions apply. This appendix specifies the metric definitions, sampling and gene-support rules, and aggregation. Complete model scores and gene-panel sensitivity appear in Appendix H.

### F.1 Population metrics and gene support

#### Abundance: mean error and response correlation

Within each source, predicted, or target population, recorded units receive equal weight and cells receive equal weight within units. Let ***µ****_s_*, ***µ****_p_*, and ***µ****_t_* be the resulting means of raw-count log(1 + **n**), including zeros. For a gene panel *P*,

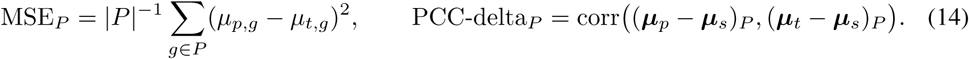

MSE measures the distance from predicted to observed target abundance; lower is better. PCC-delta measures whether predicted and observed changes from the source agree across genes; higher is better. A correlated response can still have the wrong magnitude, so the two scores answer different questions. These are population summaries rather than paired-cell residuals, as in count-based perturbation evaluation (Wei et al., 2026; Wu et al., 2025). PCC-delta is undefined when the product of centered-vector norms is at most 10^−12^.

#### Spatial W1: comparing locations for each gene

For each gene, we project the predicted and observed RNA coordinates onto 64 directions and average the exact weighted one-dimensional Wasserstein distances (Bonneel et al., 2015; Peyré & Cuturi, 2019). Supported units receive equal weight, gene-positive cells receive equal weight within units, and molecules receive equal weight within cells. For coordinate distributions *µ_p,g_* and *µ_t,g_*,

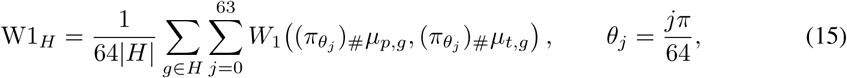

where *π_θ_*(**r**) = **r**^⊤^(cos *θ,* sin *θ*). Lower W1 means closer coordinate distributions. Oracle averages over target-present genes. Joint averages only over genes with RNA in both prediction and target, because a missing predicted gene has no coordinate distribution to compare.

Source references and targets are selected independently without replacement, using seed 42 and unit-stratified allocations, with at most 2,000 cells per population. Both settings use the same selected targets and generate *B* = min(2000*, N_t_*) cells. Every generated cell and all its RNA are scored.

#### Coverage: which target genes can receive a spatial score

Let 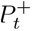 contain target-present panel genes and 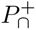 those present in both prediction and target. Coverage is 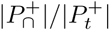, calculated per response before aggregation. It measures gene presence: one predicted RNA is enough to cover a gene. It does not measure recovery of the correct molecule count.

A missing target gene contributes zero predicted counts to abundance evaluation but has no defined spatial distance. A spurious predicted gene contributes abundance error but has no target geometry. Consequently, a low conditional Joint W1 can coexist with poor counts or missing genes. We report it alongside abundance metrics and coverage, without inventing locations or spatial penalties for missing genes. Strict full-target W1 requires predicted geometry for every target-present gene. An empty prediction has zero coverage when target RNA is present and undefined conditional W1; a panel without target-present genes has undefined coverage.

#### Gene panels: all genes and the top-100 abundance-ranked genes

Both panels score the same saved predictions. The top-100 panel is ranked only after predictions are fixed: observed source and target counts are scaled per cell to 10,000, transformed by log(1 + *n*), and ranked by absolute Welch statistic (Welch, 1947), with lexicographic ties. The first min(100*, G*) genes are retained. No significance threshold is required, and predictions do not select the genes. The table shorthand “top-100 DEGs” therefore denotes a ranking panel, not confirmed donor-level differential abundance or spatial-response genes. This count normalization is used for ranking, not for the reported abundance predictions.

### F.2 Aggregation, improvements, and ranks

Scores are averaged with equal weights over responses within family, families within dataset/split, and splits within dataset. Cross-dataset summaries weight datasets equally and keep the two OOD axes separate. Undefined values are not replaced by zero; a summary requiring complete scores remains undefined if an input is missing.

Twelve datasets use three split seeds and report a mean and sample standard deviation with denominator *n−*1. This variation describes different partitions, not independent biological replication or a confidence interval. ATM-RT and Ddx54-PD1 each have one fixed split, so no split-variation error bar is available.

For a lower-is-better loss *L*, the split-paired percentage improvement of method *m* over Resampling on dataset *d* is

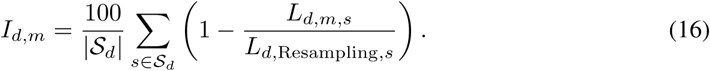

Each ratio compares the same setting, split, and responses. Positive improvement means lower error than Resampling; negative improvement means higher error. Reference errors at most 10^−12^ make the percentage undefined. The heatmap uses this improvement for its error panels and raw PCC-delta for its correlation panel, so higher displayed values are better throughout. The raw-metric arrows still point downward for W1/MSE and upward for PCC-delta.

Performance tables instead report raw split-averaged scores. Mean ranks order methods by these raw scores within each dataset, assign average ranks to ties, and then average equally across datasets within each OOD axis. Lower mean rank is better. We rank Oracle W1 and Joint MSE/PCC-delta only when all eight methods have valid scores throughout the axis. We do not combine metrics into an overall score or rank conditional Joint W1 by itself. Oracle and Joint also use different gene support, so their W1 values do not form a direct common-support comparison.

## G Within-dataset analyses of observed RNA responses

This appendix gives the settings and results of each analysis of observed RNA populations. All comparisons are made within a dataset and cellular context; no biological effects are pooled across datasets. Shared controls, genes, and experimental units make comparison counts different from numbers of independent biological replicates. These exploratory analyses describe associations, transfer under specified predictors, or population composition.

### G.1 Abundance–spatial response associations

#### G.1.1 Settings

We ask whether larger changes in RNA abundance accompany larger spatial changes within a dataset. The main analysis compares source and target populations within one cellular context, using the same supported genes in both conditions. Let *a* concatenate mean counts per cell across these genes and let *s* concatenate their fixed 128-dimensional random Fourier distribution embeddings. We plot

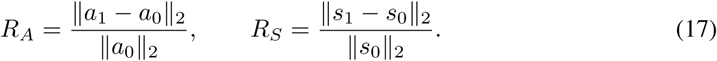

Cell summaries are averaged within units, and eligible units receive equal weight within each condition. Each arm requires at least one unit with 20 cells, and each comparison requires at least five common supported genes. Spatial coordinates use the all-RNA centroid/RMS frame. The spatial quantity measures a change in a distribution embedding, not molecular displacement; its scale depends on that representation. Gene sets may differ between comparisons. Dataset colors indicate provenance, and no pooled cross-dataset slope or correlation is fitted.

A separate gene-level analysis asks whether genes with larger absolute abundance log-fold changes also have larger spatial changes within one context. It correlates the absolute log_2_ ratio of target to source mean counts with the norm of the difference between mean spatial embeddings. Counts include zero-count cells and are averaged equally across donors. Spatial summaries weight gene-positive cells equally within donors and donors equally within conditions. Eligibility requires at least 100 cells per unit–context and, for each gene, at least 20 positive cells and 100 molecules in every donor–condition population. This descriptive gene-support rule is separate from the condition-level support and training-only feature preprocessing used for cell-level decoding in Appendix G.2.

### G.1.2 Results

Of 2,152 candidate population comparisons, 469 fail arm-level cell support and 55 fail common-gene support. The remaining 1,628 comparisons across 14 datasets appear in Figure 3a. The stricter every-unit support analysis is retained separately. The scatter describes variation in response magnitudes across context–condition comparisons; it does not test a common biological relationship across datasets.

Table 10 reports all ten gene-level correlations. The point estimates include 0.264 in Adipose adipocytes, 0.086 in GBM tumor cells, and −0.139 in GBM macrophages/monocytes. Their weak and mixed signs support reporting abundance and spatial responses separately. A negative correlation concerns response magnitudes across genes; it does not indicate movement toward the nucleus. GBM Low count/Necrotic remains a descriptive quality-control stratum.

**Table 10:** Within-dataset association between absolute abundance log-fold change and spatial response magnitude. Each correlation is computed separately within one cellular context; no cross-dataset pooling or correlation-difference test is performed. All supported contexts are shown, including the descriptive QC stratum.

| Dataset | Context | Pairs | Genes | Spearman $\rho$ |
| --- | --- | --- | --- | --- |
| Adipose | Endothelial | 4 | 68 | +0.146 |
| Adipose | Adipocytes | 4 | 49 | +0.264 |
| Adipose | APC | 4 | 30 | +0.106 |
| Adipose | Mural | 4 | 18 | +0.164 |
| GBM-oHSV | Tumor cells | 8 | 190 | +0.086 |
| GBM-oHSV | Macro./Mono. | 8 | 49 | -0.139 |
| GBM-oHSV | Vasculature | 7 | 54 | -0.195 |
| GBM-oHSV | Neutrophils | 6 | 36 | +0.125 |
| GBM-oHSV | T cells | 5 | 24 | +0.250 |
| GBM-oHSV | Low count/Necrotic (QC) | 4 | 9 | -0.617 |

The saved pair-bootstrap resamples complete donors jointly for both measures. Its percentile ranges are diagnostic rather than validated 95% confidence intervals. No test of differences between cell-type correlations is claimed, and weak correlation does not establish independence or abundance-independent regulation.

## G.2 Incremental spatial information for condition decoding

### G.2.1 Settings

We test whether spatial summaries improve classification of the observed condition after retaining an identical abundance representation and covariates. Each dataset–context has one multiclass classifier covering all its supported conditions. This task predicts a condition label for an unseen cell drawn from the same collection; it does not generate a target RNA cloud or test a new perturbation.

#### Support and cell splits

Before PCA or classification, a condition must have at least 100 cells within the context, and a context must retain at least two conditions. For each condition with *n* cells, a fixed random split holds out *n/*5 cells and uses the remainder for training. This gives at least 20 test cells per retained condition. This analysis-specific requirement is stricter than the 50-cell population-release threshold because each class must support a separate test set. Conditions below the decoding threshold are excluded from this analysis, not interpreted as biologically absent. We use three fixed split seeds and identical cell identities in the two compared models. Splitting is stratified by condition at the cell level; donor, section, or acquisition-group identities are not held out.

#### Features and training-only preprocessing

For cell *i* and gene *g*, the abundance input is 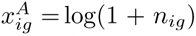, retaining observed zero counts. The spatial input concatenates the five per-gene descriptors in Appendix G.3: mean radius, finite-count-corrected first and second angular harmonic powers, and their differences from the other-gene RNA background. These summaries use the all-RNA centroid/RMS coordinate frame. They are not full RNA coordinates and do not include nuclear fractions or compartment labels. The shared covariates are log-one-plus total RNA count, detected-gene fraction, and the fraction of missing spatial features.

Feature means and variances use finite training entries only. A feature requires at least two finite training values, standard deviation above 10^−12^, and range above 10^−12^. Missing entries are imputed with the training mean, and retained features are standardized using training statistics. Separate abundance and spatial PCAs retain up to 64 components, limited by the number of training cells, retained features, and numerical rank. Covariance eigendecomposition is exact with at most 2,048 retained features; larger matrices use a fixed-seed randomized covariance subspace with up to 84 dimensions and four iterations. PC scores are rescaled to unit training variance. The 32-component sensitivity uses the exact first 32 columns of these same fitted projections. Covariates are standardized without PCA. Held-out cells never determine feature selection, imputation, scaling, or loadings.

The compared inputs are

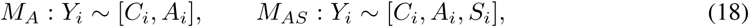

where *C_i_* contains the shared covariates and *A_i_, S_i_* are the abundance and spatial PC scores. The abundance columns, covariates, and their transformations are exactly identical in both models for a given split and component setting. Thus the contrast tests incremental information relative to a compressed abundance representation, not information conditional on the entire count profile.

#### Classifier and metrics

Class-balanced multinomial logistic regression minimizes

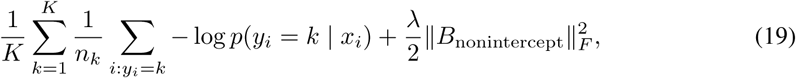

where *K* is the number of retained conditions and *n_k_* their training-cell counts. All nonintercept coefficients are penalized. Fitting uses L-BFGS-B with at most 2,000 iterations. The combined model starts from the fitted abundance coefficients and zero spatial coefficients; the penalized training objective must not increase. This nesting does not guarantee improved held-out classification.

The primary setting is up to 64 PCs per branch and *λ* = 1. The 32-PC setting and *λ ∈ {*0.001, 0.01, 0.1, 1, 10} provide sensitivity analyses. No configuration is selected for having the highest test score. Balanced accuracy is mean recall across retained conditions; the plotted gain is 100(BA*_AS_−*BA*_A_*) in percentage points. Each point in Figure 3b averages the same three splits within one context. When counting positive gains, differences within 10^−12^ in balanced-accuracy units are numerical ties. Ordinary accuracy, macro-F1, and class-balanced log loss are also retained. A uniform-probability reference has balanced accuracy 1*/K* and class-balanced log loss log *K*. Different class counts and reused cells prevent interpreting dataset medians as a common task difficulty scale or independent biological replications.

### G.2.2 Results

The support rules retain 426 contexts in all 14 datasets and 13,813,925 cells. Every abundance and spatial branch retains all 64 requested components. Across three cell splits, two component settings, and five penalties, all 25,560 individual classifiers converge. All 1,278 primary split comparisons are usable; no context is removed after inspecting its score. Table 11 reports the primary results and the matched 32-PC sensitivity.

**Table 11:** Cell-level condition decoding by dataset. Each context uses three identical stratified cell splits for abundance-only and spatially augmented classifiers. Both component settings use *λ* = 1. Gains are medians of context-level mean balanced-accuracy differences, in percentage points. The two count columns refer to 64 PCs: a positive three-split mean and improvement in every split, respectively, treating differences within 10^−12^ balanced-accuracy units as numerical ties. These counts are descriptive, not significance tests.

| Dataset | Contexts | Mean > 0 | All splits > 0 | Gain: 64 PCs | Gain: 32 PCs |
| --- | --- | --- | --- | --- | --- |
| Adipose | 6 | 0 | 0 | -1.672 | -1.540 |
| ATM-RT | 1 | 0 | 0 | -0.115 | -0.123 |
| Breast-FUS | 46 | 19 | 8 | -0.901 | +0.033 |
| Cardiac-MI | 50 | 18 | 7 | -0.664 | -0.283 |
| Ddx54-PD1 | 1 | 0 | 0 | -0.120 | -0.174 |
| DSS-Colitis | 8 | 0 | 0 | -0.091 | -0.027 |
| GBM-oHSV | 8 | 7 | 5 | +0.666 | +0.591 |
| Gut-Microbiota | 66 | 5 | 2 | -1.240 | -0.589 |
| Kidney-IRI | 20 | 13 | 6 | +0.036 | +0.107 |
| Lung-Drug | 11 | 2 | 2 | -0.105 | -0.050 |
| PF-Astro | 1 | 0 | 0 | -0.371 | -0.261 |
| Sickness | 102 | 101 | 99 | +4.851 | +5.581 |
| Spinal-KO | 83 | 9 | 1 | -3.006 | -1.859 |
| UC-VDZ | 23 | 14 | 6 | +0.357 | +0.401 |

#### Primary results by dataset

Sickness has positive three-split mean gains in 101/102 contexts and gains in every split in 99 contexts; its median gain is 4.85 percentage points. GBM-oHSV improves in 7/8 contexts, including five with improvement in all splits, with a median gain of 0.67 points. UC-VDZ improves in 14/23 contexts with a median gain of 0.36 points. Kidney-IRI improves in 13/20, but its median gain is only 0.04 points.

Adipose and DSS-Colitis have negative mean balanced-accuracy gains in all six and eight contexts, respectively; their medians are −1.67 and −0.09 points. In Breast-FUS, Cardiac-MI, Gut-Microbiota, Lung-Drug, and Spinal-KO, gains are positive in 19/46, 18/50, 5/66, 2/11, and 9/83 contexts, respectively, with negative dataset medians. The single supported context in each of ATM-RT, Ddx54-PD1, and PF-Astro also has a negative mean gain. These are separate within-dataset summaries, not estimates of a pooled spatial-information effect.

Loss and classification recall need not move together. For example, adding spatial features improves mean class-balanced log loss in four of six Adipose contexts and seven of eight DSS-Colitis contexts, despite negative balanced-accuracy gains in every context of both datasets. Reduced log loss therefore does not imply improved argmax classification, and we retain the fixed primary metric rather than choosing whichever score improves.

Five primary split configurations in four Breast-FUS myeloid contexts have class-balanced test log loss above 10. Their smallest test classes contain 33–97 cells, so they pass the condition-support rule. The affected angular features have only two or three finite training values and training standard deviations of approximately 6.0 ×10^−8^–4.2 ×10^−7^. Standardizing ordinary held-out values by these small scales produces spatial PC magnitudes of approximately 3.25 × 10^4^–1.29 × 10^5^, compared with training maxima of 10.19–22.42. A broader scan finds amplified held-out projections beyond these five high-loss configurations, predominantly in Breast-FUS. This is a stability limitation of sparse-feature preprocessing, not an optimizer-convergence failure. All affected configurations remain in the summaries, and their balanced accuracies pass independent reconstruction. Negative gains therefore do not establish an absence of spatial information.

#### Component and penalty sensitivity

With 32 PCs, Sickness has a median gain of 5.58 points and GBM-oHSV 0.59 points. Increasing to 64 PCs raises the separate Sickness abundance and combined model-score medians from 54.75% to 56.83% and from 60.69% to 62.25%, while the median paired gain falls to 4.85 points. A smaller incremental gain can therefore accompany improved scores for both models. The medians in Table 11 describe paired gains and are not calculated by subtracting the separate score medians.

Table 12 retains all five penalties for the 64-PC representation. Sickness has positive median gains throughout the grid, ranging from 4.85 to 6.76 points, and GBM-oHSV from 0.23 to 0.67 points. Adipose, DSS-Colitis, Gut-Microbiota, PF-Astro, and Spinal-KO have negative medians throughout. Other datasets can change median sign with regularization, including ATM-RT, Ddx54-PD1, and UC-VDZ. These sensitivities do not replace the fixed *λ* = 1 primary results with a test-selected configuration.

**Table 12:** Regularization sensitivity of condition decoding. Values are median context-level balanced-accuracy gains in percentage points with 64 PCs per branch. Columns retain the full fixed penalty grid; *λ* = 1 is primary. Each context first averages its three fixed cell splits.

| Dataset | $\lambda = 0.001$ | 0.01 | 0.1 | 1 | 10 |
| --- | --- | --- | --- | --- | --- |
| Adipose | −1.113 | −1.186 | −1.237 | −1.672 | −1.894 |
| ATM-RT | +0.328 | +0.326 | +0.331 | −0.115 | −0.373 |
| Breast-FUS | −1.156 | −1.095 | −1.419 | −0.901 | +0.011 |
| Cardiac-MI | −1.428 | −1.718 | −2.089 | −0.664 | +0.049 |
| Ddx54-PD1 | +0.070 | +0.081 | −0.016 | −0.120 | −0.062 |
| DSS-Colitis | −0.018 | −0.043 | −0.117 | −0.091 | −0.109 |
| GBM-oHSV | +0.320 | +0.227 | +0.545 | +0.666 | +0.658 |
| Gut-Microbiota | −1.798 | −1.996 | −2.311 | −1.240 | −0.750 |
| Kidney-IRI | −0.062 | +0.037 | +0.041 | +0.036 | +0.122 |
| Lung-Drug | −0.026 | +0.037 | −0.086 | −0.105 | −0.216 |
| PF-Astro | −0.208 | −0.771 | −0.414 | −0.371 | −0.300 |
| Sickness | +6.762 | +6.363 | +5.484 | +4.851 | +4.867 |
| Spinal-KO | −3.834 | −3.280 | −3.130 | −3.006 | −2.198 |
| UC-VDZ | −1.162 | −1.177 | −0.550 | +0.357 | +0.640 |

#### Reconstruction audit

All cell identities, condition labels, and three splits match the frozen support-filtered inputs, and every test condition has at least 20 cells. Independent test-metric calculations agree within 1.8 × 10^−15^ for the 64-PC primary fits and 8.9 × 10^−16^ for the 32-PC primary fits. Checks also reconstruct saved logits from coefficients and projections at every penalty, verify identical abundance inputs in each paired comparison, and confirm that adding spatial components or expanding the 32-PC models to 64 PCs decreases the penalized training objective. These numerical checks do not guarantee a gain on held-out cells or establish biological reliability.

#### Biological and technical interpretation

These results describe predictive utility for new cells within the observed collection. They do not establish unseen-donor or OOD response-generation performance, biological independence, or causality. Cells from the same donors and acquisition groups can occur in both training and test sets. Adipose condition differences include time and collection-method changes. In GBM, the 12 retained ROIs with retrieved acquisition summaries separate pre/post conditions by run/slide; metadata for other ROIs are incomplete. A classifier can therefore use measurement or sampling differences as well as biological changes. Adding components can strengthen the abundance baseline as well as the combined model, so a smaller incremental gain need not mean that the combined classifier became less accurate. The component and penalty sensitivities are descriptive; they are not tests of the absence of spatial information.

### G.3 Radial and angular response patterns

#### G.3.1 Settings

We distinguish changes in distance from the normalized RNA center from changes in angular concentration around that center. The cached spatial descriptors are mean radius, finite-count-corrected first and second angular harmonic powers, and each power’s difference from the other-gene RNA background. For a nonzero normalized coordinate, let **v** = (*x, y*)*/* ∥(*x, y*) ∥. The two harmonic vectors are *h*_1_ = (*v_x_, v_y_*) and 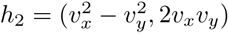; angular terms are set to zero when the radius is at most 10^−12^. For a gene with *n >* 1 RNA,

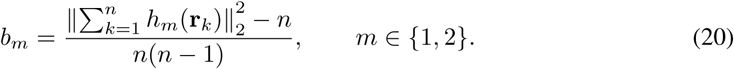

The relative power subtracts the corresponding value for other-gene RNA in the same cell; insufficient background counts yield a missing value. These descriptors use the all-RNA centroid/RMS frame, not a nuclear or membrane boundary.

For each context–condition comparison, the plotted coordinates are median signed target-minus-source effects over supported genes: normalized mean radius horizontally and relative first-harmonic power vertically. The two descriptors can have different supported gene sets. A dataset needs at least five comparisons for a display group; at least 75% negative angular medians defines the predominantly negative group, and other eligible datasets enter the mixed-direction group. These groups were chosen descriptively after inspecting the results. Symmetric-log axes use a linear threshold of 10^−3^ and retain zero and sign. Hollow markers indicate a median angular-valid cell fraction below 0.2.

### G.3.2 Results

Figure 3c shows 131 comparisons in the predominantly negative angular group: DSS-Colitis, GBM-oHSV, and PF-Astro. Adipose, Breast-FUS, Cardiac-MI, Gut-Microbiota, Kidney-IRI, Lung-Drug, Sickness, Spinal-KO, and UC-VDZ have mixed directions under this rule; their 1,491 comparisons appear in panel d. These two panels therefore cover 1,622 comparisons in 12 datasets. ATM-RT and Ddx54-PD1 each have only three comparisons; their six points remain in the analysis records without a group assignment. Insufficient comparisons are not classified as absence of a pattern.

The displayed radius and angular effects describe different aspects of RNA organization. Low angular observability, particularly in DSS-Colitis, limits interpretation of a predominant direction. Neither the display groups nor relative angular power establish a shared mechanism or movement toward a specific compartment.

## G.4 Population composition and within-cluster changes

### G.4.1 Settings

We ask how much of a change in a mean spatial descriptor is associated with changing cluster proportions and how much occurs within fixed clusters. Each dataset–context is fitted separately using the five descriptors defined in Appendix G.3. The expanded analysis uses a fixed sample of at most 500 cells per unit–context–condition, selected without replacement using seeds derived from seed 101 and the group identifiers. All RNA of a selected cell contributes to its descriptors. Genes require at least 20 positive cells and 100 molecules in every training-source unit; at least three eligible genes and three observed gene features per retained cell are required.

When several source units exist, the first half of their sorted identifiers supplies training controls, and remaining source units and target units outside the training set supply evaluation. A separately recorded repair uses source-only training units when the original split leaves no target units. Training and evaluation units remain disjoint in that repaired branch. With only one source unit, the output is explicitly a fitted-control description without an independent source holdout. Unverified biological-unit identities remain a separate limitation.

Training-control medians impute missing descriptors. An unadjusted branch retains the descriptors; a count-adjusted branch removes their fitted dependence on log-one-plus total normalization-molecule count, using training controls and equal unit weights. Training-control scaling and two-cluster K-means are then fitted with equal unit weights and ten starts. The seed-7123 assignment is frozen for evaluation; seeds 101, 211, 307, 401, and 503 measure initialization agreement. Two clusters is a fixed descriptive choice. Cluster labels are not aligned across datasets, contexts, or branches.

For condition *a* and cluster *s*, let *p_as_* be its proportion and *m_as_* its feature mass, both averaged equally over evaluation units. Define *µ_as_* = *m_as_/p_as_* for *p_as_ >* 0. The population mean is *x̅_a_* = *_s_ p_as_µ_as_*, and its change decomposes exactly as

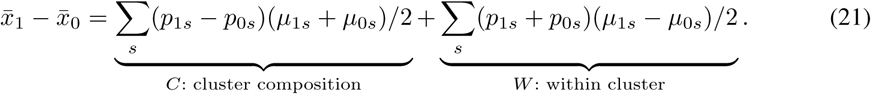

Thus *C* measures the contribution from changing proportions, and *W* the contribution from changing means within clusters. Opposite signs indicate cancellation. An absent cluster makes its mean undefined; it is not assigned a zero mean.

For the compact display, each comparison has a within-cluster contribution share

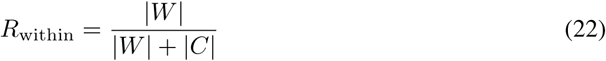

for radius and relative first-harmonic power separately. A share above 0.5 indicates that the within-cluster term has greater absolute magnitude; a share below 0.5 indicates a larger composition term. Zero total absolute contribution is undefined and is excluded. The horizontal and vertical coordinates in Figure 3e are the radius and angular shares on linear axes. This representation combines the comparisons in one panel but omits their effect signs and net magnitudes. Hollow markers indicate minimum initialization ARI below 0.8.

For descriptive grouping, a dataset needs at least five defined comparisons. At least two-thirds with |*W| > |C|* defines within-cluster dominance, and at least two-thirds with |*C|≥|W|* defines composition dominance; otherwise the contributions are mixed. These thresholds summarize the observed comparisons and are not significance tests.

### G.4.2 Results

The analysis contains 1,601 computed comparisons, of which 1,412 have defined shares for both displayed descriptors in the unadjusted branch. Table 13 gives each dataset’s supported comparison counts. In Breast-FUS, 66/68 radius changes have larger within-cluster contributions, whereas 51/68 angular changes have larger composition contributions. DSS-Colitis, GBM-oHSV, and Lung-Drug also meet the descriptive thresholds for within-cluster radius dominance and composition dominance for angle.

**Table 13:**
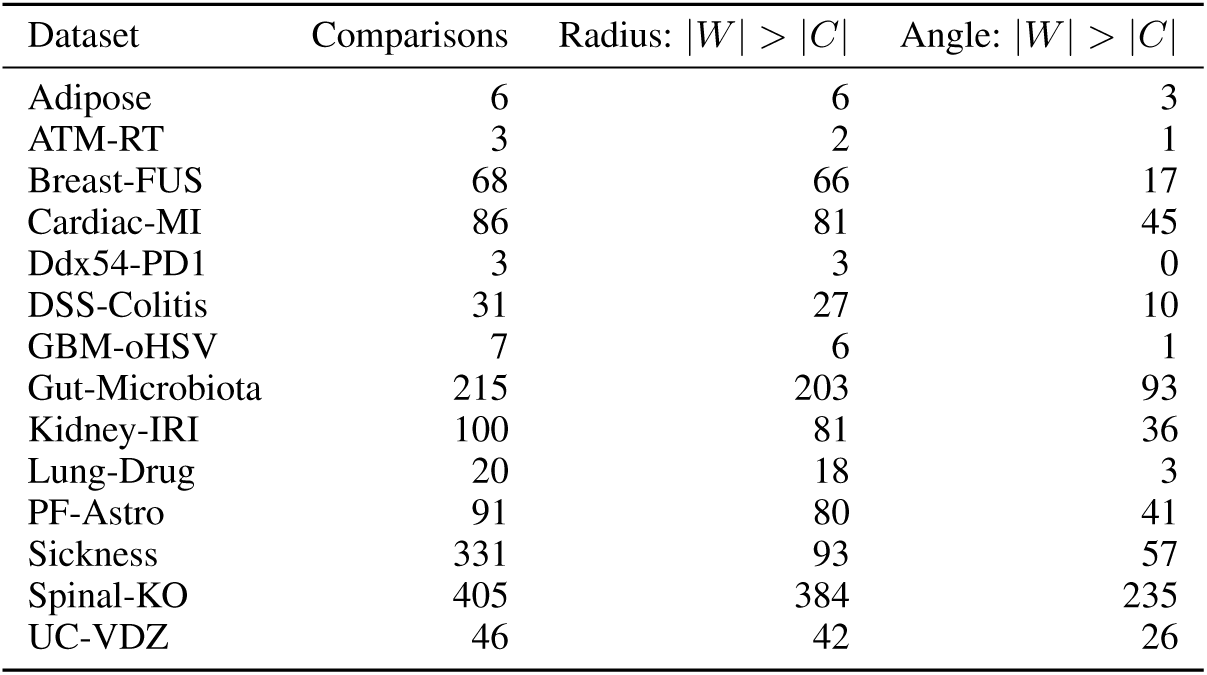
Within-cluster contributions by dataset. Counts refer to comparisons with defined radius and angular contribution shares. The last two columns count |*W| > |C|* for the indicated descriptor; remaining comparisons have at least as large a composition contribution. Comparisons can share controls and <u>units.</u>

Adipose has within-cluster radius dominance in all six comparisons, with three angular comparisons on each side of 0.5. Cardiac-MI, Gut-Microbiota, Kidney-IRI, PF-Astro, Spinal-KO, and UC-VDZ each meet the radius threshold but have mixed angular contributions. In Sickness, composition terms dominate 238/331 radius and 274/331 angular comparisons; DSS/STAG and LPS/PIC/PLA need not share the same radius pattern. Its decomposition is defined in 331 of 510 comparisons. ATM-RT and Ddx54-PD1 each have three comparisons and remain unclassified by the five-comparison grouping rule, while their defined shares remain visible in the compact plot.

The count-adjusted branch retains the same joint display memberships at the two-thirds threshold. Membership checks at 60%, two-thirds, 75%, and 80% are retained rather than selecting the threshold with the clearest separation. These groups summarize observed comparisons; they do not identify biological states or trace cells across conditions.

#### Paired-donor checks

Table 14 gives a paired Adipose/GBM analysis using all eligible Adipose cells and a fixed sample of at most 500 GBM cells per donor–condition–context. The first half of the frozen donor list supplied training controls; there were two held-out pairs per Adipose context and two to four per GBM context. All 100 feature–context–branch decompositions were defined, with maximum identity error below 3 × 10^−16^ and minimum initialization ARI 0.903 across 20 context–branch fits.

**Table 14:** Within-dataset decomposition of the mean normalized RNA radius. U: unadjusted; C: training-control count-adjusted. Values are multiplied by 10^3^ for readability; they are not physical distances. Δ = *O* + *W*, where *O* is the occupancy term and *W* the within-cluster term. *n* denotes held-out donor pairs; ARI is the minimum initialization agreement. Rows are separate descriptive analyses, not independent replications.

| Dataset | Context | Branch | $n$ | ARI | $10^3\Delta$ | $10^3O$ | $10^3W$ |
| --- | --- | --- | --- | --- | --- | --- | --- |
| Adipose | Endothelial | U | 2 | 0.990 | +10.375 | +0.004 | +10.371 |
| Adipose | Endothelial | C | 2 | 0.974 | +14.530 | -0.026 | +14.555 |
| Adipose | Adipocytes | U | 2 | 1.000 | +10.128 | +0.019 | +10.109 |
| Adipose | Adipocytes | C | 2 | 1.000 | +16.575 | +0.243 | +16.332 |
| Adipose | APC | U | 2 | 0.992 | +3.902 | +0.621 | +3.281 |
| Adipose | APC | C | 2 | 0.953 | +3.990 | +0.517 | +3.472 |
| Adipose | Mural | U | 2 | 0.929 | +14.978 | -0.616 | +15.594 |
| Adipose | Mural | C | 2 | 0.968 | +14.706 | -0.160 | +14.866 |
| GBM-oHSV | Tumor cells | U | 4 | 0.998 | +5.706 | -0.216 | +5.921 |
| GBM-oHSV | Tumor cells | C | 4 | 0.990 | +4.476 | -0.162 | +4.638 |
| GBM-oHSV | Macro./Mono. | U | 4 | 0.981 | +7.015 | -0.814 | +7.828 |
| GBM-oHSV | Macro./Mono. | C | 4 | 0.930 | +9.196 | -0.590 | +9.785 |
| GBM-oHSV | Vasculature | U | 4 | 0.991 | +1.206 | -0.469 | +1.675 |
| GBM-oHSV | Vasculature | C | 4 | 0.967 | -3.161 | -0.253 | -2.908 |
| GBM-oHSV | Neutrophils | U | 3 | 0.987 | -6.189 | -0.536 | -5.653 |
| GBM-oHSV | Neutrophils | C | 3 | 0.998 | -8.612 | -0.300 | -8.312 |
| GBM-oHSV | T cells | U | 3 | 0.987 | +0.971 | -2.769 | +3.740 |
| GBM-oHSV | T cells | C | 3 | 0.903 | +4.802 | -2.746 | +7.548 |
| GBM-oHSV | Low count/Necrotic (QC) | U | 2 | 0.986 | -0.604 | -0.800 | +0.196 |
| GBM-oHSV | Low count/Necrotic (QC) | C | 2 | 1.000 | -3.678 | -1.390 | -2.288 |

Within-cluster radius terms exceed proportion terms in all four Adipose contexts in both branches. For Endothelial, the unadjusted total, proportion, and within-cluster terms are 0.010375, 0.000004, and 0.010371; adjusted values are 0.014530, −0.000026, and 0.014555. In GBM tumor cells, the unadjusted values are 0.005706, −0.000216, and 0.005921, showing partial cancellation. GBM Vasculature changes sign after count adjustment, from 0.001206 to −0.003161, and is not treated as a robust directional result. These normalized quantities are not physical distances. Exact decomposition and stable initialization do not establish biological-state identity, causal mediation, or donor generalization.

## G.5 Nuclear RNA fractions and compartment geometry

### G.5.1 Settings

We compare the fraction of a gene’s RNA assigned to the nucleus with its enrichment relative to nuclear area. This analysis requires native nuclear assignments and matched cell and nuclear geometry; RNA coordinates alone do not supply those annotations. Six datasets support the reported comparisons: Adipose, ATM-RT, Cardiac-MI, GBM-oHSV, Lung-Drug, and Spinal-KO. Adipose uses registered cell and nuclear polygons. The other five use native molecule-level nuclear flags joined to native projected areas, with count and coordinate checks against the reconstructed RNA populations.

For cell *i* and gene *g* with retained count *n_ig_ >* 0, define

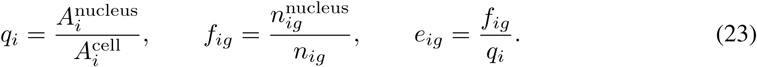

Here *f* is the observed nuclear RNA fraction, whereas *e* compares that fraction with the fraction of projected cell area occupied by the nucleus. The same gene-positive cells with usable projected areas contribute to both quantities. We average cells equally within each unit–condition–context, then average units equally within conditions and subtract source from target. Complete paired donors therefore receive equal weight in their paired difference. The mean of cell-level ratios is not replaced by a ratio of population means. Geometry summaries of *q* use all geometry-usable cells. These are two-dimensional area-normalized measurements, not three-dimensional concentrations.

For the native-flag datasets, every included unit–condition–context must contain at least 20 cells. Each displayed gene must cover all such units in both arms, with at least 20 geometry-usable gene-positive cells and 100 RNA molecules per unit. For Adipose, complete four-pair coverage requires at least 100 geometry-usable cells per donor–condition–context, plus the same per-gene thresholds of 20 positive cells and 100 molecules. A context with a missing eligible unit is not rescued by pooling the remaining units. Unsupported comparisons are omitted rather than treated as unchanged.

Each point in Figure 3f is the median across supported genes of their target-minus-source effects in *f* and *e*. Fraction changes are shown in percentage points; enrichment changes are dimensionless. The two medians use the same gene panel within a point, but panels can differ across comparisons. A median enrichment change cannot be reconstructed by dividing median fraction and area changes. Hollow circles identify fewer than 20 supported genes, not fewer than 20 cells or a failed significance test. No pooled cross-dataset estimate or formal discovery claim is made.

#### Adipose registration and boundary checks

Native cell and nuclear polygons are matched to 14,007 reconstructed source cells from four paired donors. Registration requires an overall inside-cell molecule fraction of at least 0.95, at least 0.99 of cells having half their RNA inside, and a median polygon-to-native-area ratio between 0.8 and 1.2. Per-cell geometry also requires an inside-cell fraction of at least 0.8. Of the matched cells, 14,000 pass these checks and seven are excluded; approximately 97.95% of assigned RNA lies inside its corresponding cell polygon. These checks establish coordinate registration, not biological correctness of every boundary. Primary estimates preserve the published molecule assignments. Sensitivity checks retain only RNA inside the cell polygon or erode that polygon by 0.25 and 0.5 native-coordinate units; these units are not relabeled as micrometers without verified scale information.

### G.5.2 Results

There are 519 supported context–condition comparisons in the six datasets. Table 15 reports their support separately; comparisons sharing controls, genes, or units are not independent biological replicates.

**Table 15:** Support for nuclear RNA and geometry comparisons. Gene ranges are per context–condition comparison after complete unit coverage and per-unit support checks. The last column counts comparisons plotted with hollow circles because fewer than 20 genes remain. Comparison counts are not biological-replicate counts.

| Dataset | Contexts | Comparisons | Genes | < 20 genes |
| --- | --- | --- | --- | --- |
| Adipose | 4 | 4 | 18–68 | 1 |
| ATM-RT | 1 | 3 | 298 | 0 |
| Cardiac-MI | 45 | 76 | 1–124 | 54 |
| GBM-oHSV | 9 | 9 | 1–190 | 6 |
| Lung-Drug | 10 | 20 | 23–3,714 | 0 |
| Spinal-KO | 155 | 407 | 3–234 | 185 |

#### Adipose

Across the four paired donors, 165 gene–context entries meet complete support in APC, adipocytes, endothelial cells, and mural cells. Nuclear fractions increase in 138 entries. The four context medians increase by 2.92–3.94 percentage points, while median enrichment decreases by 0.276–0.854. Table 16 gives the full supported background. This contrast is sensitive to donor composition in adipocytes: omitting donor SP02 −changes median enrichment from −0.593 to +0.141. The other three contexts retain negative median enrichment in each leave-one-donor-out check. SP01–SP04 are donor identifiers, not conditions or timepoints. These checks describe heterogeneity and do not create independent replications.

**Table 16:**
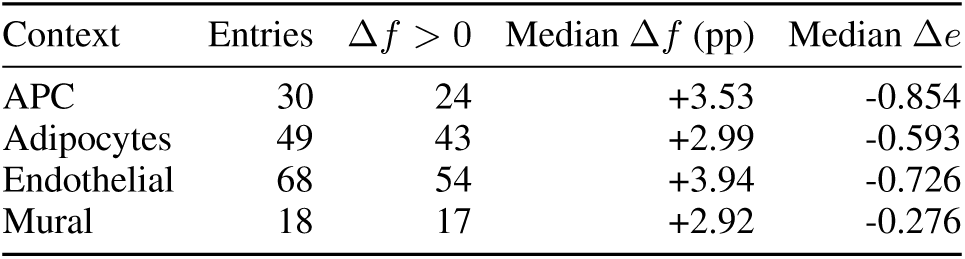
Complete-coverage compartment background. All 165 gene–context entries satisfying the engineering coverage criterion in all four donor pairs are retained. Nuclear fraction and enrichment are distinct estimands; genes and contexts reuse the same donors.

The nuclear-to-cell area ratio increases in all seven annotated contexts within each donor, as shown in Figure 8. A larger nuclear area fraction can therefore coexist with a larger nuclear RNA fraction but lower area-normalized enrichment.

**Figure 8:**
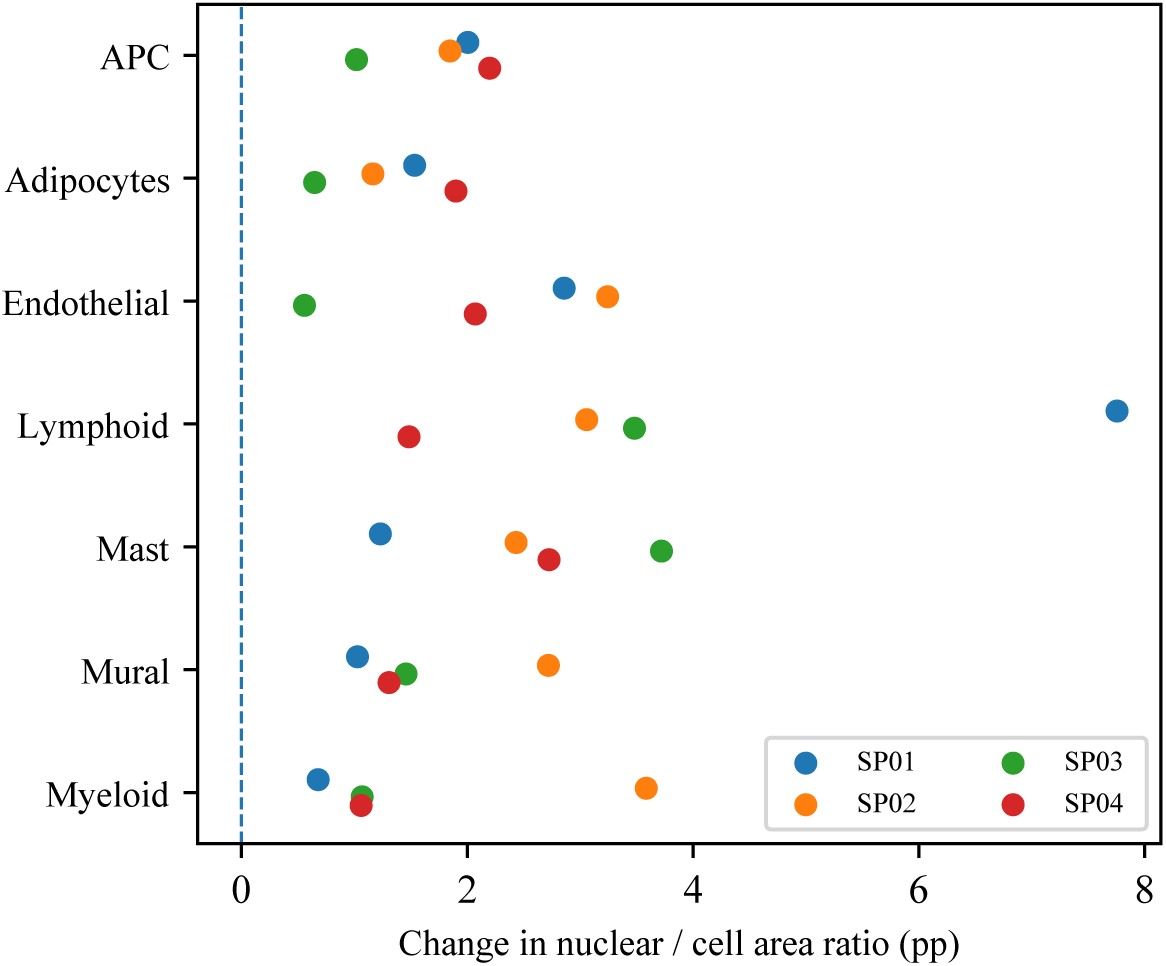
Adipose compartment geometry. Points show within-donor changes in the mean cell-level nuclear-to-cell projected-area ratio, Weightloss minus Obese, in percentage points. SP01–SP04 identify four paired donors. The seven contexts reuse these donors, so the 28 differences are not independent replicates. All changes are positive in these samples; sampling and segmentation remain possible explanations.

**Figure 9:**
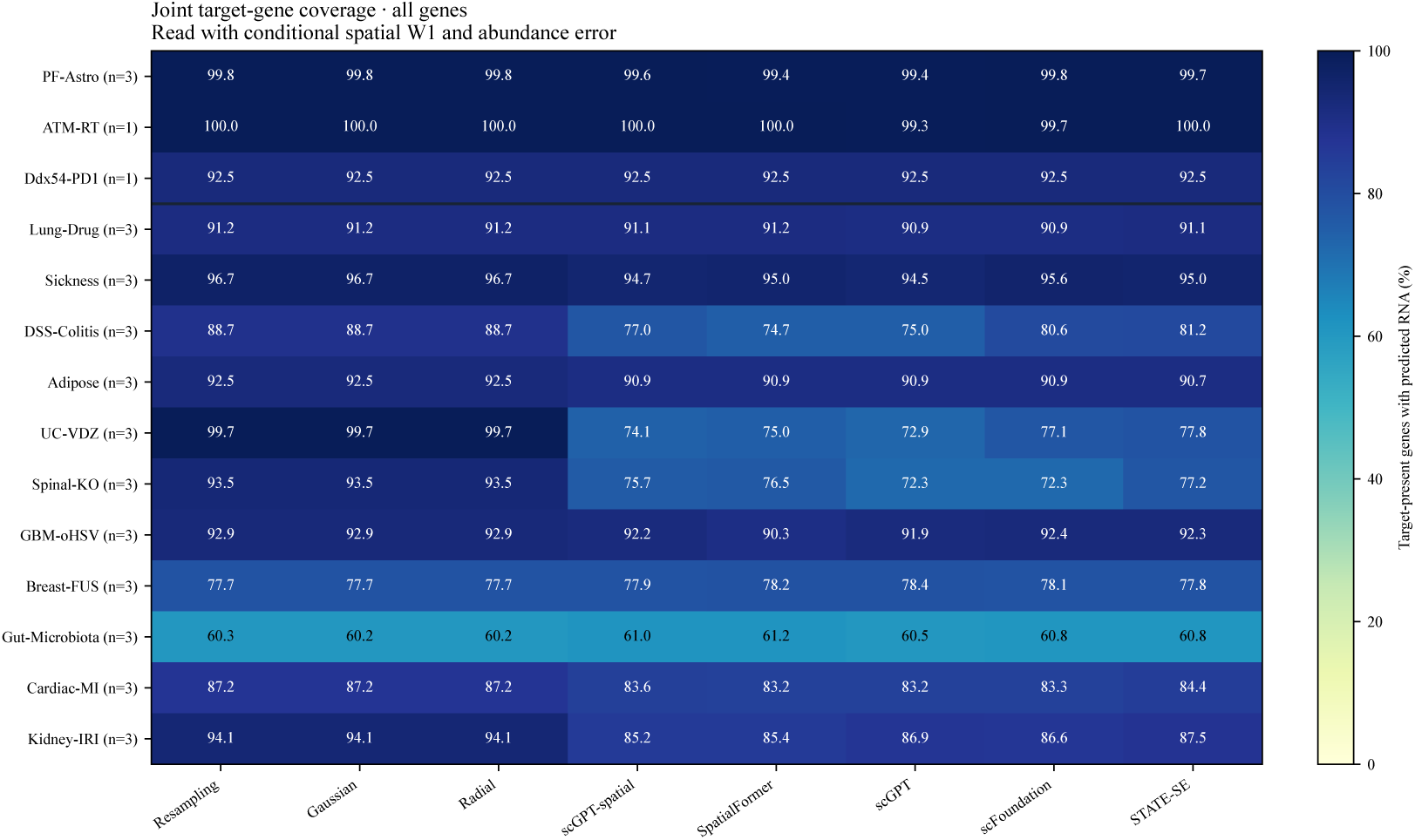
Joint target-gene coverage. Response/family/split-averaged percentage of target-present genes with predicted RNA, using identical targets across methods. A rounded mean near 100% need not imply complete coverage in every response, as required for strict full-target W1. The horizontal line separates Perturbation OOD from Cellular-context OOD.

Table 17 gives gene-level examples selected for descriptive illustration. Nuclear fraction increases by 4.49 percentage points for CAVIN1 in adipocytes and 11.35 points for DPT in APC, but decreases by 7.56 points for LITAF in endothelial cells. Their displayed directions persist across donors and under containment and cell-boundary erosion. These checks do not validate nuclear boundaries or remove sampling, detection, or collection-method effects. Adipose samples were collected surgically before weight loss and by needle biopsy afterward, so collection method and time remain entangled with condition.

**Table 17:** Post hoc compartment examples. Nuclear fractions are averaged equally over cells with nonzero counts for the corresponding gene within donor and then over four paired donors. Changes and donor ranges are percentage points (pp). Ranges are not confidence intervals; examples are not multiplicity-controlled discoveries.

| Context | Gene | Obese (%) | Weightloss (%) | $\Delta$ (pp) | Donor range |
| --- | --- | --- | --- | --- | --- |
| Adipocytes | *CAVIN1* | 9.62 | 14.11 | +4.49 | [+2.50, +5.79] |
| Endothelial | *LITAF* | 39.69 | 32.13 | -7.56 | [-10.04, -5.11] |
| APC | *DPT* | 11.04 | 22.39 | +11.35 | [+8.12, +16.33] |

#### ATM-RT

All three comparisons retain 298 genes. Relative to ATM-intact cells, the median nuclear fraction decreases by 2.26–6.06 percentage points in the radiation, ATM-loss, and combined conditions, while median enrichment increases by 0.179–0.481. Each condition has one section, so the three contrasts do not establish reproducibility across independent biological replicates.

#### GBM-oHSV

Tumor cells retain 190 genes and show median decreases of 3.46 percentage points in nuclear fraction and 0.126 in enrichment. Macrophages/monocytes retain 49 genes and decrease by 6.25 percentage points and 0.140, respectively. Both comparisons cover eight complete patient pairs, and both median directions persist when each patient is omitted in turn, despite heterogeneous patient-level effects. Six of the nine displayed contexts have fewer than 20 supported genes, which limits interpretation of those other context medians.

#### Lung-Drug

The 20 comparisons cover ten contexts. To compare the two drug responses on identical support, airway-epithelium summaries use their 3,022 common genes: adding AZD3965 to bleomycin decreases median nuclear fraction by 1.14 percentage points and increases enrichment by 0.026, whereas adding VB124 increases the fraction by 0.40 points and enrichment by 0.017. Thus the fraction directions differ within this context even though both enrichment changes are positive. Stored acquisition groups are not assumed to be independent biological replicates.

#### Spinal-KO

The 407 comparisons cover 155 contexts, with 185 comparisons supported by fewer than 20 genes. A matched-panel illustration within the v1 context uses 205 common genes: Brach zic14ko increases median nuclear fraction by 0.76 percentage points and enrichment by 0.033, whereas Brach zic3ko decreases the fraction by 1.24 points but increases enrichment by 0.079. Each knockout has one target unit; multiple contexts or genes do not replace biological replication.

#### Cardiac-MI

The 76 comparisons cover 45 contexts and all four combinations of fraction and enrichment signs: 21 have higher fraction and lower enrichment, 17 have lower fraction and higher enrichment, five increase in both, and 33 decrease in both. Interpretation is limited by projected-area coverage: only 67.6% of cells have valid areas overall, with condition-dependent coverage, and 54 comparisons have fewer than 20 supported genes. These support limitations preclude a single directional summary of Cardiac-MI.

#### Interpretation and audit scope

Independent checks reconstruct native counts, compartment identities, unit means, signed effects, support flags, and the plotted aggregation. These numerical checks do not establish biological-unit independence or remove segmentation and sampling effects. A larger nuclear RNA fraction does not necessarily imply more nuclear RNA molecules, and enrichment is conditional on the available projected geometry. None of these descriptive results identifies nuclear export, retention, or active trafficking, or shows that the compartment effects explain gains in condition decoding.

## G.6 Shared spatial transformations

### G.6.1 Settings

We test whether a simple transformation shared across genes predicts spatial responses in held-out groups and genes. Identity is compared with a radial map, a similarity map, and an affine map. Matching source and target unit IDs remain in the same fold; up to four folds leave at least two training groups and one evaluation group per arm. Two gene folds separate fitting genes from evaluation genes. Gene eligibility is determined from training support; observed test support determines only which predictions can be scored. Acquisition groups without established biological independence remain computational evaluation groups.

Maps minimize an equal-gene, regularized discrepancy between source and target random Fourier summaries, using 64 fitting features, bounded transformations, and at most 300 optimization steps. Evaluation uses 128 features and two fixed sketches, averaging squared distribution-summary error equally across evaluated genes and groups. Gain is identity error minus transformed-source error, so positive values favor the learned map. All coordinates retain the all-RNA centroid/RMS reference.

### G.6.2 Results

Of 2,152 candidates, 463 comparisons yield evaluable outputs; 1,633 lack group-holdout support and 56 lack common-gene support. Table 18 distinguishes positive mean gain from gain that stays positive in every evaluated fold and both sketches. Radial maps improve mean error in four of five Adipose comparisons, but only one improves throughout evaluation. Shared affine maps have positive mean gain in three of five GBM-oHSV comparisons, while none improves throughout; all 200 Sickness mean affine gains are nonpositive.

**Table 18:** Shared-map diagnostics within each dataset. Entries give the number of comparisons with positive mean held-out gain / positive gain in every evaluated fold and both evaluation sketches. Gains compare each shared map with identity. These counts are descriptive and reuse groups, genes, and controls.

| Dataset | Comparisons | Radial | Similarity | Affine |
| --- | --- | --- | --- | --- |
| Adipose | 5 | 4 / 1 | 1 / 0 | 1 / 0 |
| DSS-Colitis | 14 | 8 / 3 | 7 / 2 | 4 / 1 |
| GBM-oHSV | 5 | 2 / 0 | 0 / 0 | 3 / 0 |
| Gut-Microbiota | 175 | 58 / 16 | 34 / 11 | 27 / 9 |
| PF-Astro | 41 | 13 / 2 | 6 / 1 | 6 / 2 |
| Sickness | 200 | 6 / 0 | 1 / 0 | 0 / 0 |
| UC-VDZ | 23 | 8 / 2 | 3 / 1 | 4 / 1 |

The tested shared maps therefore provide uneven transfer. Failure of these particular maps does not identify the intrinsic dimension of spatial responses or prove that every more complex model would generalize better.

## G.7 Response transfer across cellular contexts

### G.7.1 Settings

We test whether the mean response in other cellular contexts predicts a held-out context within the same dataset and condition contrast. Unit holdouts are synchronized across contexts. A gene must have a training response in at least two other contexts and meet training-source support in the query context. The primary predictor never fits the query context’s target response. Abundance responses use log(1 + mean count); normalized mean radius and relative angular descriptors provide separate spatial responses. Each descriptor is evaluated against a no-change prediction in its own units. A separate amplitude-fitted diagnostic uses query-context training targets and is not the primary transfer comparison.

### G.7.2 Results

There are 450 evaluable comparisons; 1,497 candidates lack group support, 164 lack common observed support, and 41 lack other contexts. Table 19 reports positive mean gains and gains that remain positive in every fold. In Adipose, abundance transfer improves all six mean comparisons, radius transfer improves four, and angular gains are less consistent across folds. UC-VDZ also has positive abundance and radius cases. In Sickness, neither radius nor relative angular transfer improves mean error in any of the 202 evaluated comparisons.

**Table 19:** Other-context response transfer within each dataset. Entries give positive mean gain / positive gain in every evaluated fold, relative to predicting no change. Count, radius, and angular responses are evaluated separately in their native units.

| Dataset | Comparisons | Abundance | Radius | Angular |
| --- | --- | --- | --- | --- |
| Adipose | 6 | 6 / 4 | 4 / 2 | 4 / 0 |
| DSS-Colitis | 14 | 7 / 2 | 2 / 0 | 1 / 0 |
| GBM-oHSV | 7 | 3 / 2 | 3 / 3 | 5 / 2 |
| Gut-Microbiota | 196 | 97 / 33 | 35 / 12 | 53 / 10 |
| Sickness | 202 | 20 / 0 | 0 / 0 | 0 / 0 |
| UC-VDZ | 25 | 23 / 12 | 18 / 8 | 11 / 4 |

Thus a common mean response transfers in some settings but is not a general predictor of spatial changes. This conclusion concerns the specified predictor; it does not rule out every shared response program. Split-half agreement also varies by descriptor, so limited response reliability can reduce apparent transfer independently of model capacity.

## G.8 Responses across recorded injury stages

### G.8.1 Settings

We summarize source-to-stage abundance and spatial-descriptor changes in datasets with recorded stage labels. Each comparison remains within a dataset and cellular context. DSS-Colitis has recorded days 3, 9, 21, and 35 without time-matched controls. Kidney-IRI has one animal per stage and a day-14 sham reference. Cardiac-MI compares days 7 and 28 with day-7 sham samples, and independent animal identities are not established for every stored group. These observations sample different cell populations at different stages; the benchmark itself does not train or evaluate temporal extrapolation.

### G.8.2 Results

The audited outputs contain 31 nonempty context–stage comparisons in DSS-Colitis, 100 in Kidney-IRI, and 98 in Cardiac-MI. An additional empty Cardiac-MI output remains in the execution record and is not counted as an observed comparison. These outputs describe abundance and spatial changes at recorded stages, but the available controls and replication do not identify a lag between the two responses. We therefore make no claim about lead–lag relationships, recovery order, individual-cell trajectories, or temporal prediction.

## G.9 Auxiliary analysis: Localization-only screening

### G.9.1 Settings

#### Scope and units

Feature extraction covers the reconstructed source population of 13,896,237 cells and 3,658,217,075 RNA molecules without cell or RNA subsampling; the supported comparisons are selected using the criteria below. This extraction scope differs from the filtered release inventory in Table 2. Registered directed transitions and contexts were deduplicated across split seeds, yielding 125 dataset–transition pairs, 1,448 transition–context combinations, and 1,399,492 gene-level comparisons. PF-Astro contributes 87 supported contrasts; Lung-Drug uses the drug contrasts. UC-VDZ observations are grouped by patient in this analysis, unlike the cell-type holdouts used for prediction. Thus these 1,448 combinations describe the scan’s grouping, not the number of held-out prediction tasks. Genes, controls, and experimental units recur across comparisons, so these counts do not represent independent experiments. The exploratory analysis includes benchmark holdouts and is separate from model evaluation.

#### Spatial support

Each condition–context population requires at least 50 cells, and each included unit or acquisition group requires at least 20 cells. A gene requires at least 20 cells with three or more transcripts and at least one other-gene transcript in every compared group. We match on exact gene count and [2 log_2_ *N_i_*], where *N_i_* is total RNA in cell *i*. Each shared stratum requires at least two cells per group; its mass is the minimum occupancy across groups, with total shared mass at least 10. Normalized stratum masses give every compared group the same count/depth distribution. Paired data are matched within each complete unit pair; unpaired data require common strata across all included units. Unit summaries are weighted equally within each arm.

#### Radial effect

Coordinates are centered and scaled using each cell’s all-RNA centroid and root-mean-square radius. All transcripts of an eligible gene contribute to its cell-level radial cumulative distribution function (CDF). Let *F_s_* and *F_t_* denote the matched, unit-averaged source and target CDFs. We report

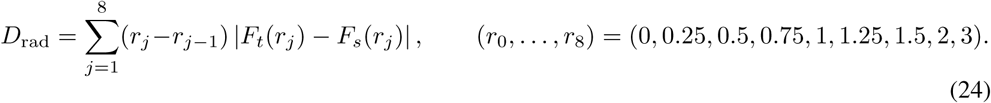

This fixed-grid radial discrepancy measures relative radial organization; it omits angular changes and differs from the full-coordinate sliced Wasserstein metric used for model evaluation. The multi-metric analysis additionally evaluates exact full-support radial W1 (including the tail beyond radius 3), mean radius, centroid offset, dispersion, anisotropy, and Gaussian-kernel MMD approximations (Gretton et al., 2012) on four rotation-invariant cell features at bandwidths 0.25, 0.5, and 1.0. The MMD approximations use random Fourier features (Rahimi & Recht, 2007) with 32 fixed frequencies per bandwidth and seed 20260918; they are not exact MMD on full two-dimensional RNA point clouds. The eight statistics other than exact radial W1 also have gene-minus-other-RNA background-adjusted versions, giving 17 statistics in total. Coordinates remain cell-normalized, not annotated organelle coordinates.

#### Screening criteria and statistical inference. Abundance criteria

For each unit, mean counts, mean log(1 + count), and detection fractions include zero-count cells. Denote the arm differences in the latter two quantities by Δ*ℓ* and Δ*d*. We scan fold margins *f∈{*1.05, 1.10, 1.20}, with *f* = 1.10 designated as primary before this reanalysis. Small point-estimate effects require the raw arm mean-count ratio, without pseudocounts, to lie in [1*/f, f*], |Δ*ℓ|≤*ln *f*, and |Δ*d|≤*0.1. These descriptive bounds do not establish equivalence. The stricter test intersects two one-sided equivalence tests (Schuirmann, 1987) on unit mean-log1p counts, log-transformed unit raw mean counts, and unit detection fractions. Their margins are ln *f*, ln *f*, and 0.1, respectively; the joint equivalence *p* value is the largest of the three. Paired or Welch *t* tests (Welch, 1947) use biological units rather than cells and retain their parametric assumptions.

#### Inference at the biological-unit level

Tests require identified independent units and at least three complete pairs or three units per arm. Adipose and GBM-oHSV supply paired tests; DSS-Colitis and Sickness supply unpaired tests. UC-VDZ has one patient per scan context, Kidney-IRI one mouse per timepoint, and Spinal-KO one target animal per KO. The remaining datasets supply descriptive acquisition or pooled-reference groups without established independent biological-unit support. Spatial tests enumerate paired sign flips or unpaired unit-label allocations preserving arm sizes (up to 100,000 enumerations). Validity depends on unit independence and the relevant symmetry or exchangeability assumptions. Each statistic is normalized by its permutation-invariant null mean and maxT calibration accounts for the 17 statistics within a comparison (Winkler et al., 2014). The localization-only conjunction takes the larger of the spatial omnibus and joint equivalence *p* values; a failed raw-ratio bound sets it to one. BH (Benjamini & Hochberg, 1995) and BY (Benjamini & Yekutieli, 2001) adjustments are retained within datasets and globally over all valid comparison–margin combinations, without filtering the testing family to selected candidates. The primary criterion is global BY *q <* 0.05 at *f* = 1.10, supplemented by unit stability and background-adjusted support. Paired stability uses signed/vector agreement across pairs; unpaired leave-one-unit-out stability is an internal diagnostic, not independent replication. Exact radial W1 has no direction, so a signed CDF proxy is used for that stability check.

### G.9.2 Results

Table 20 gives all dataset summaries. Of 98,904 spatially supported comparisons, 21,081 have sufficient biological units for inference. The within-comparison spatial test has *p <* 0.05 in 334 cases; these are not across-comparison FDR discoveries. At margins 1.05, 1.10, and 1.20, respectively, 56, 124, and 187 of those cases meet the small-abundance point-estimate bounds. No comparison passes joint abundance equivalence at nominal *p <* 0.05 for margins 1.05 or 1.10. At 1.20, 135 pass equivalence, but none overlaps the 334 spatial cases. Thus the nominal conjunction is already empty before global correction; the absence of final discoveries cannot be attributed solely to BY conservatism. Table 21 reports all three margins.

**Table 20:** Multi-metric analysis of observed localization. Counts are gene–contrast–context comparisons, not independent experiments. Supported means sufficient RNA and matching support; tested additionally requires independent biological units. Spatial denotes within-comparison maxT *p <* 0.05, before across-comparison correction. Small+spatial additionally meets the point-estimate bounds at *f* = 1.10. Final discoveries use the primary global BY and stability/background criteria. Dashes denote unavailable inference, not evidence of no effect.

| Dataset | Scanned | Supported | Tested | Spatial | Small+spatial | Final |
| --- | --- | --- | --- | --- | --- | --- |
| PF-Astro | 42,195 | 1,726 | — | — | — | — |
| ATM-RT | 894 | 882 | — | — | — | — |
| Ddx54-PD1 | 15,018 | 6,737 | — | — | — | — |
| Lung-Drug | 110,132 | 20,670 | — | — | — | — |
| Sickness | 479,910 | 51,929 | 20,052 | 271 | 113 | 0 |
| DSS-Colitis | 29,140 | 1,500 | 843 | 44 | 5 | 0 |
| Adipose | 2,862 | 88 | 88 | 0 | 0 | 0 |
| UC-VDZ | 7,000 | 1,050 | — | — | — | — |
| Spinal-KO | 79,116 | 5,635 | — | — | — | — |
| GBM-oHSV | 4,800 | 100 | 98 | 19 | 6 | 0 |
| Breast-FUS | 90,000 | 252 | — | — | — | — |
| Gut-Microbiota | 462,825 | 2,695 | — | — | — | — |
| Cardiac-MI | 45,600 | 382 | — | — | — | — |
| Kidney-IRI | 30,000 | 5,258 | — | — | — | — |
| Total | 1,399,492 | 98,904 | 21,081 | 334 | 124 | 0 |

**Table 21:** Sensitivity to the abundance-equivalence margin. Small counts all spatially supported comparisons satisfying the descriptive abundance bounds. The spatial, equivalence, and conjunction columns use nominal *p <* 0.05 among 21,081 biologically testable comparisons; small+spatial intersects the descriptive bounds with the spatial test. Global BY reports corrected conjunctions. All margins share the same spatial tests; the primary margin is 1.10.

| Fold | Small | Spatial | Small+spatial | Equivalence | Conjunction | Global BY |
| --- | --- | --- | --- | --- | --- | --- |
| 1.05 | 15,735 | 334 | 56 | 0 | 0 | 0 |
| 1.10 | 31,273 | 334 | 124 | 0 | 0 | 0 |
| 1.20 | 53,791 | 334 | 187 | 135 | 0 | 0 |

**Table 22:** Benchmark results on PF-Astro.

| Baseline | O-W1 ↓ | J-W1 ↓ | Cov. ↑ | MSE ↓ | PCC-delta ↑ |
| --- | --- | --- | --- | --- | --- |
| *All genes* |  |  |  |  |  |
| Resampling | 0.251 ± 0.0081 | **0.254 ± 0.0082** | **0.998 ± 0.00042** | **0.00646 ± 0.00025** | −0.00043 ± 0.032 |
| Gaussian | 0.253 ± 0.0069 | 0.257 ± 0.0063 | **0.998 ± 0.00042** | 0.00834 ± 0.00054 | 0.033 ± 0.021 |
| Radial | 0.298 ± 0.0025 | 0.267 ± 0.0069 | **0.998 ± 0.00042** | 0.00834 ± 0.00054 | 0.033 ± 0.021 |
| scGPT-spatial | 0.249 ± 0.0084 | 0.263 ± 0.012 | 0.996 ± 0.0029 | 0.0109 ± 0.004 | 0.0347 ± 0.0059 |
| SpatialFormer | 0.249 ± 0.0063 | 0.263 ± 0.0063 | 0.994 ± 0.0017 | 0.0109 ± 0.0015 | 0.0385 ± 0.016 |
| scGPT | 0.249 ± 0.0063 | 0.267 ± 0.013 | 0.994 ± 0.0026 | 0.0133 ± 0.0032 | **0.0482 ± 0.022** |
| scFoundation | 0.249 ± 0.0063 | **0.254 ± 0.0082** | **0.998 ± 0.00042** | **0.00646 ± 0.00025** | −0.00043 ± 0.032 |
| STATE-SE | **0.248 ± 0.0066** | 0.256 ± 0.01 | 0.997 ± 0.002 | 0.00778 ± 0.0022 | −0.00753 ± 0.02 |
| *Top-100 DEGs* |  |  |  |  |  |
| Resampling | 0.262 ± 0.0084 | **0.257 ± 0.0086** | **0.999 ± 0.00065** | **0.0116 ± 0.00066** | −0.0245 ± 0.043 |
| Gaussian | 0.262 ± 0.0077 | 0.258 ± 0.0076 | **0.999 ± 0.00065** | 0.0124 ± 0.00041 | 0.0124 ± 0.026 |
| Radial | 0.286 ± 0.0072 | 0.27 ± 0.0081 | **0.999 ± 0.00065** | 0.0124 ± 0.00041 | 0.0124 ± 0.026 |
| scGPT-spatial | 0.26 ± 0.0094 | 0.263 ± 0.012 | 0.997 ± 0.0033 | 0.0127 ± 0.0016 | 0.0646 ± 0.037 |
| SpatialFormer | 0.259 ± 0.007 | 0.261 ± 0.011 | 0.995 ± 0.0018 | 0.0129 ± 0.00091 | 0.0514 ± 0.023 |
| scGPT | 0.26 ± 0.0071 | 0.265 ± 0.014 | 0.996 ± 0.0028 | 0.0142 ± 0.0005 | **0.0683 ± 0.055** |
| scFoundation | 0.259 ± 0.007 | **0.257 ± 0.0086** | **0.999 ± 0.00065** | **0.0116 ± 0.00066** | −0.0245 ± 0.043 |
| STATE-SE | **0.258 ± 0.0071** | 0.257 ± 0.0086 | 0.997 ± 0.0027 | 0.0125 ± 0.0013 | −0.0423 ± 0.014 |

**Table 23:**
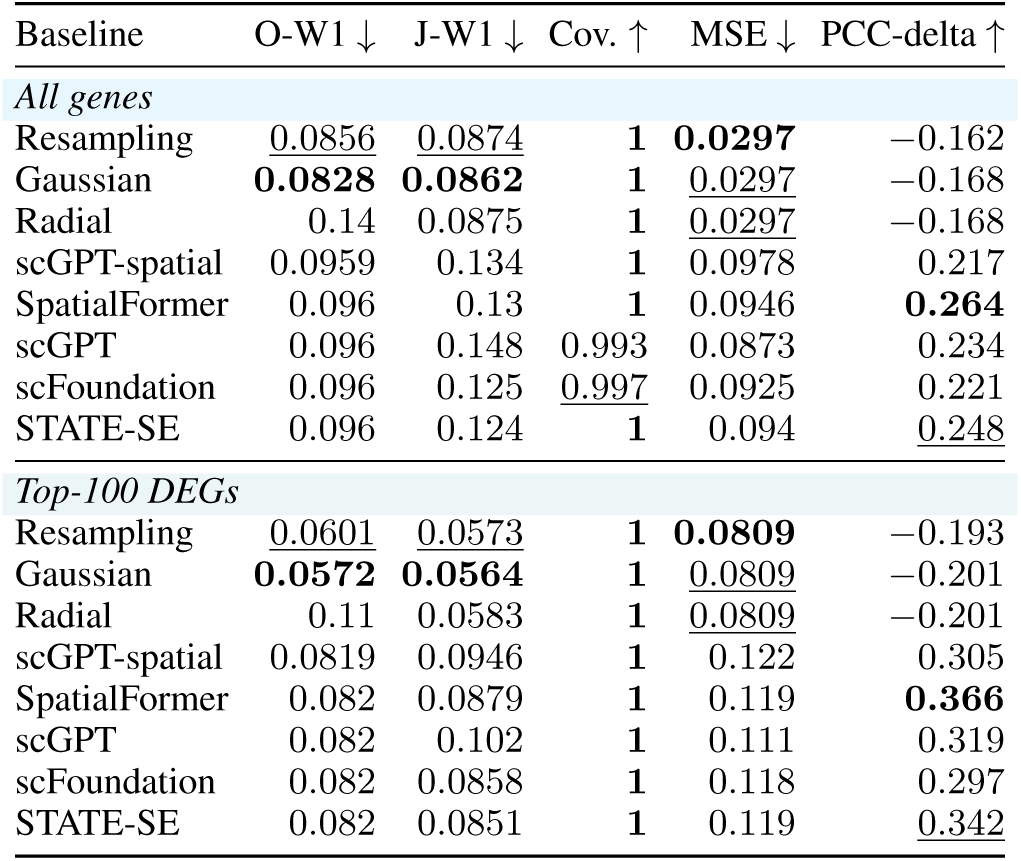
Benchmark results on ATM-RT.

| Baseline | O-W1 ↓ | J-W1 ↓ | Cov. ↑ | MSE ↓ | PCC-delta ↑ |
| --- | --- | --- | --- | --- | --- |
| *All genes* |  |  |  |  |  |
| Resampling | 0.0856 | 0.0874 | **1** | **0.0297** | −0.162 |
| Gaussian | **0.0828** | **0.0862** | **1** | 0.0297 | −0.168 |
| Radial | 0.14 | 0.0875 | **1** | 0.0297 | −0.168 |
| scGPT-spatial | 0.0959 | 0.134 | **1** | 0.0978 | 0.217 |
| SpatialFormer | 0.096 | 0.13 | **1** | 0.0946 | **0.264** |
| scGPT | 0.096 | 0.148 | 0.993 | 0.0873 | 0.234 |
| scFoundation | 0.096 | 0.125 | 0.997 | 0.0925 | 0.221 |
| STATE-SE | 0.096 | 0.124 | **1** | 0.094 | 0.248 |
| *Top-100 DEGs* |  |  |  |  |  |
| Resampling | 0.0601 | 0.0573 | **1** | **0.0809** | −0.193 |
| Gaussian | **0.0572** | **0.0564** | **1** | 0.0809 | −0.201 |
| Radial | 0.11 | 0.0583 | **1** | 0.0809 | −0.201 |
| scGPT-spatial | 0.0819 | 0.0946 | **1** | 0.122 | 0.305 |
| SpatialFormer | 0.082 | 0.0879 | **1** | 0.119 | **0.366** |
| scGPT | 0.082 | 0.102 | **1** | 0.111 | 0.319 |
| scFoundation | 0.082 | 0.0858 | **1** | 0.118 | 0.297 |
| STATE-SE | 0.082 | 0.0851 | **1** | 0.119 | 0.342 |

**Table 24:** Benchmark results on Ddx54-PD1.

| Baseline | O-W1 ↓ | J-W1 ↓ | Cov. ↑ | MSE ↓ | PCC-delta ↑ |
| --- | --- | --- | --- | --- | --- |
| *All genes* |  |  |  |  |  |
| Resampling | 0.305 | 0.297 | **0.925** | **0.00318** | **0.197** |
| Gaussian | 0.301 | 0.295 | **0.925** | <u>0.00318</u> | <u>0.188</u> |
| Radial | 0.36 | 0.297 | **0.925** | <u>0.00318</u> | <u>0.188</u> |
| scGPT-spatial | <u>0.296</u> | 0.294 | **0.925** | 0.00591 | −0.184 |
| SpatialFormer | 0.297 | **0.294** | **0.925** | 0.00713 | −0.227 |
| scGPT | 0.297 | 0.294 | **0.925** | 0.00674 | −0.247 |
| scFoundation | **0.296** | 0.294 | **0.925** | 0.00769 | −0.345 |
| STATE-SE | 0.298 | <u>0.294</u> | **0.925** | 0.0046 | −0.0685 |
| *Top-100 DEGs* |  |  |  |  |  |
| Resampling | <u>0.0767</u> | **0.0635** | **1** | **0.0877** | <u>0.0662</u> |
| Gaussian | **0.0763** | 0.064 | **1** | <u>0.0879</u> | 0.0233 |
| Radial | 0.107 | <u>0.0635</u> | **1** | <u>0.0879</u> | 0.0233 |
| scGPT-spatial | 0.0897 | 0.0805 | **1** | 0.177 | −0.0523 |
| SpatialFormer | 0.0919 | 0.0796 | **1** | 0.193 | −0.0174 |
| scGPT | 0.0919 | 0.0807 | **1** | 0.189 | **0.0721** |
| scFoundation | 0.0899 | 0.0799 | **1** | 0.225 | −0.27 |
| STATE-SE | 0.0914 | 0.0781 | **1** | 0.144 | −0.127 |

**Table 25:**
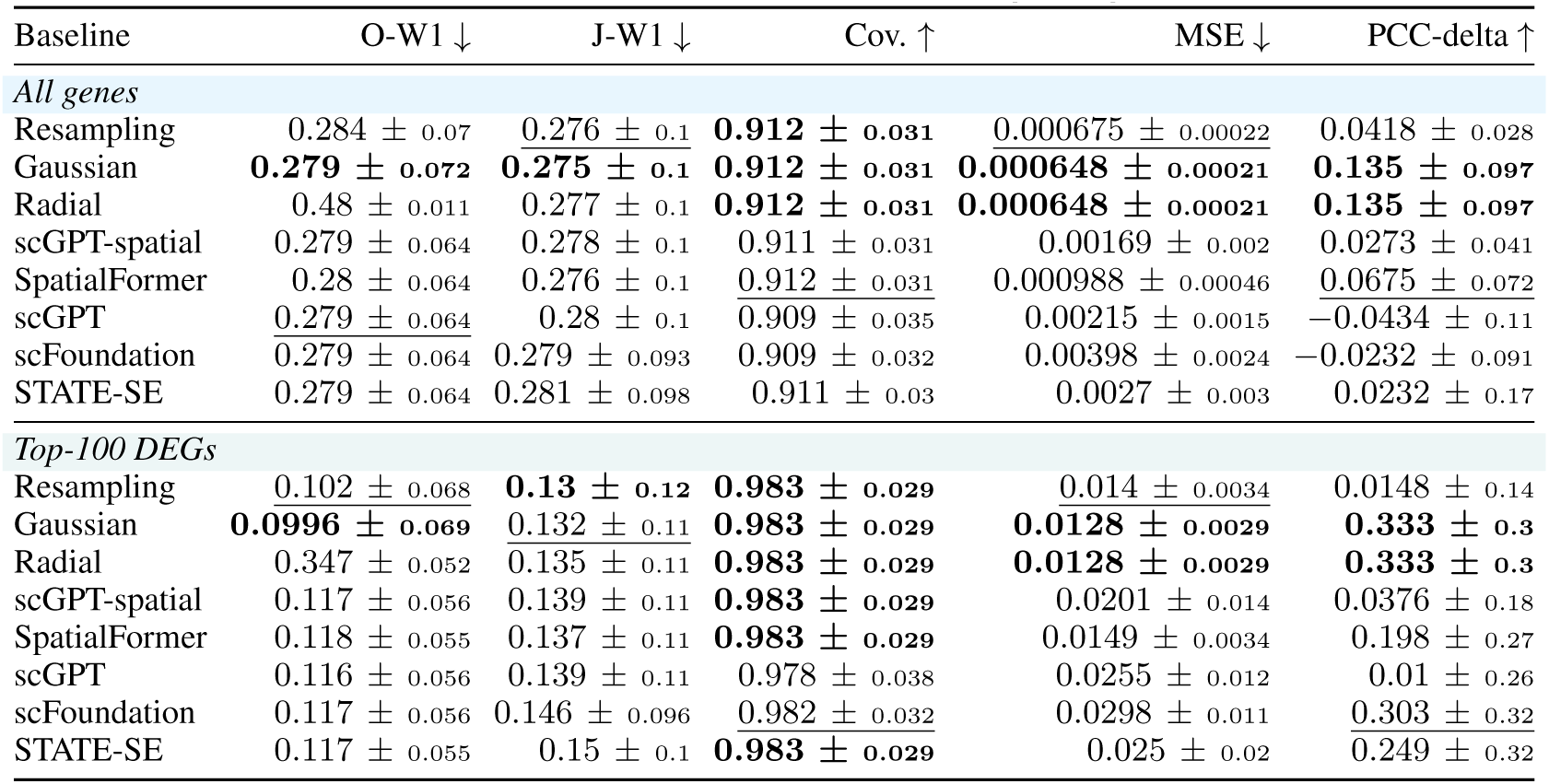
Benchmark results on Lung-Drug.

#### Interpretation

Screening yields depend on gene panels, common support, and thresholds; they cannot be interpreted as biological relocalization prevalence. Small abundance effects or scalar-mean equivalence leave room for changes in the full count distribution. Limited replication also constrains inference: with four Adipose donor pairs, the minimum attainable symmetric sign-flip *p* value is 2*/*2^4^ = 0.125. Normalization, cell morphology, segmentation, subpopulation composition, and acquisition can influence radial effects. These observations require independent confirmation and do not establish causal relocation, information conditional on the full count profile, or predictability in unseen conditions.

## H Complete benchmark results

The following tables report all-gene and top-100-DEG results for every dataset and baseline, using the aggregation in Appendix F. Values are means and sample standard deviations over three split seeds, except for the fixed ATM-RT and Ddx54-PD1 holdouts. A held-out context may recur across seeds; the error bars describe split variability, not biological replication. Within each gene panel and metric column, bold and underlined entries mark the best and second-best distinct unrounded means, respectively; exact ties share a marker. These column-wise markers do not define an overall Joint rank.

Joint W1 entries are conditional on shared nonempty genes and must be interpreted alongside abundance metrics and target-gene coverage. Only six all-gene dataset–method summaries retain every target-present gene in every response and split, all on ATM-RT. This limits spatial-score interpretation even when conditional W1 improves over Resampling.

The mean-rank summary is reported at the end of the main experimental section in Table 3.

**Table 26:** Benchmark results on Sickness.

| Baseline | O-W1 ↓ | J-W1 ↓ | Cov. ↑ | MSE ↓ | PCC-delta ↑ |
| --- | --- | --- | --- | --- | --- |
| *All genes* |  |  |  |  |  |
| Resampling | 0.34 ± 0.068 | 0.354 ± 0.058 | **0.967 ± 0.032** | 0.00436 ± 0.0018 | -0.0279 ± 0.037 |
| Gaussian | **0.264 ± 0.075** | **0.298 ± 0.071** | **0.967 ± 0.032** | **0.00338 ± 0.0013** | **0.452 ± 0.036** |
| Radial | 0.403 ± 0.0092 | 0.315 ± 0.072 | **0.967 ± 0.032** | **0.00338 ± 0.0013** | **0.452 ± 0.036** |
| scGPT-spatial | 0.326 ± 0.063 | 0.349 ± 0.051 | 0.947 ± 0.043 | 0.0407 ± 0.00069 | 0.15 ± 0.055 |
| SpatialFormer | 0.32 ± 0.067 | 0.346 ± 0.05 | 0.95 ± 0.043 | 0.0383 ± 0.0034 | 0.157 ± 0.062 |
| scGPT | 0.326 ± 0.065 | 0.345 ± 0.051 | 0.945 ± 0.047 | 0.041 ± 0.002 | 0.156 ± 0.058 |
| scFoundation | 0.319 ± 0.076 | 0.346 ± 0.048 | 0.956 ± 0.039 | 0.0408 ± 0.00031 | 0.156 ± 0.06 |
| STATE-SE | 0.319 ± 0.066 | 0.347 ± 0.051 | 0.95 ± 0.047 | 0.0413 ± 0.0023 | 0.144 ± 0.05 |
| *Top-100 DEGs* |  |  |  |  |  |
| Resampling | 0.22 ± 0.037 | 0.235 ± 0.045 | **0.988 ± 0.014** | 0.0289 ± 0.012 | -0.0322 ± 0.072 |
| Gaussian | **0.137 ± 0.042** | **0.174 ± 0.045** | **0.988 ± 0.014** | **0.0198 ± 0.0078** | **0.542 ± 0.017** |
| Radial | 0.268 ± 0.029 | 0.187 ± 0.044 | **0.988 ± 0.014** | **0.0198 ± 0.0078** | **0.542 ± 0.017** |
| scGPT-spatial | 0.209 ± 0.037 | 0.222 ± 0.033 | 0.984 ± 0.017 | 0.0787 ± 0.0099 | 0.287 ± 0.094 |
| SpatialFormer | 0.199 ± 0.036 | 0.223 ± 0.032 | 0.987 ± 0.014 | 0.0761 ± 0.011 | 0.293 ± 0.091 |
| scGPT | 0.209 ± 0.036 | 0.222 ± 0.031 | 0.985 ± 0.017 | 0.0795 ± 0.0063 | 0.289 ± 0.09 |
| scFoundation | 0.198 ± 0.047 | 0.222 ± 0.03 | 0.986 ± 0.016 | 0.076 ± 0.0092 | 0.292 ± 0.087 |
| STATE-SE | 0.198 ± 0.047 | 0.221 ± 0.034 | 0.985 ± 0.016 | 0.0798 ± 0.0059 | 0.27 ± 0.068 |

**Table 27:** Benchmark results on DSS-Colitis.

| Baseline | O-W1 ↓ | J-W1 ↓ | Cov. ↑ | MSE ↓ | PCC-delta ↑ |
| --- | --- | --- | --- | --- | --- |
| *All genes* |  |  |  |  |  |
| Resampling | 0.352 ± 0.18 | 0.409 ± 0.22 | **0.887 ± 0.16** | 0.00244 ± 0.0011 | 0.0342 ± 0.057 |
| Gaussian | 0.351 ± 0.18 | **0.407 ± 0.22** | 0.887 ± 0.16 | **0.00222 ± 0.0012** | 0.244 ± 0.14 |
| Radial | 0.503 ± 0.02 | 0.409 ± 0.22 | 0.887 ± 0.16 | **0.00222 ± 0.0012** | 0.244 ± 0.14 |
| scGPT-spatial | 0.341 ± 0.17 | 0.431 ± 0.19 | 0.77 ± 0.19 | 0.00586 ± 0.00072 | 0.361 ± 0.042 |
| SpatialFormer | 0.341 ± 0.17 | 0.43 ± 0.18 | 0.747 ± 0.17 | 0.00614 ± 0.0023 | **0.395 ± 0.099** |
| scGPT | 0.341 ± 0.17 | 0.425 ± 0.2 | 0.75 ± 0.27 | 0.00665 ± 0.0017 | 0.34 ± 0.066 |
| scFoundation | 0.341 ± 0.17 | 0.419 ± 0.19 | 0.806 ± 0.21 | 0.00653 ± 0.0015 | 0.357 ± 0.062 |
| STATE-SE | **0.341 ± 0.17** | 0.422 ± 0.19 | 0.812 ± 0.17 | 0.00617 ± 0.00043 | 0.324 ± 0.061 |
| *Top-100 DEGs* |  |  |  |  |  |
| Resampling | 0.269 ± 0.16 | **0.311 ± 0.21** | **0.86 ± 0.24** | 0.0153 ± 0.0043 | 0.0303 ± 0.094 |
| Gaussian | 0.268 ± 0.16 | 0.311 ± 0.21 | 0.859 ± 0.24 | **0.0134 ± 0.0055** | 0.297 ± 0.23 |
| Radial | 0.452 ± 0.038 | 0.317 ± 0.21 | 0.859 ± 0.24 | **0.0134 ± 0.0055** | 0.297 ± 0.23 |
| scGPT-spatial | 0.263 ± 0.15 | 0.339 ± 0.15 | 0.73 ± 0.25 | 0.0246 ± 0.0075 | 0.411 ± 0.067 |
| SpatialFormer | 0.263 ± 0.15 | 0.327 ± 0.14 | 0.691 ± 0.24 | 0.025 ± 0.013 | **0.456 ± 0.12** |
| scGPT | **0.263 ± 0.15** | 0.32 ± 0.15 | 0.729 ± 0.34 | 0.0279 ± 0.011 | 0.371 ± 0.034 |
| scFoundation | 0.264 ± 0.14 | 0.318 ± 0.16 | 0.77 ± 0.27 | 0.0257 ± 0.011 | 0.42 ± 0.066 |
| STATE-SE | 0.263 ± 0.15 | 0.335 ± 0.17 | 0.795 ± 0.21 | 0.0221 ± 0.0037 | 0.381 ± 0.045 |

**Table 28:**
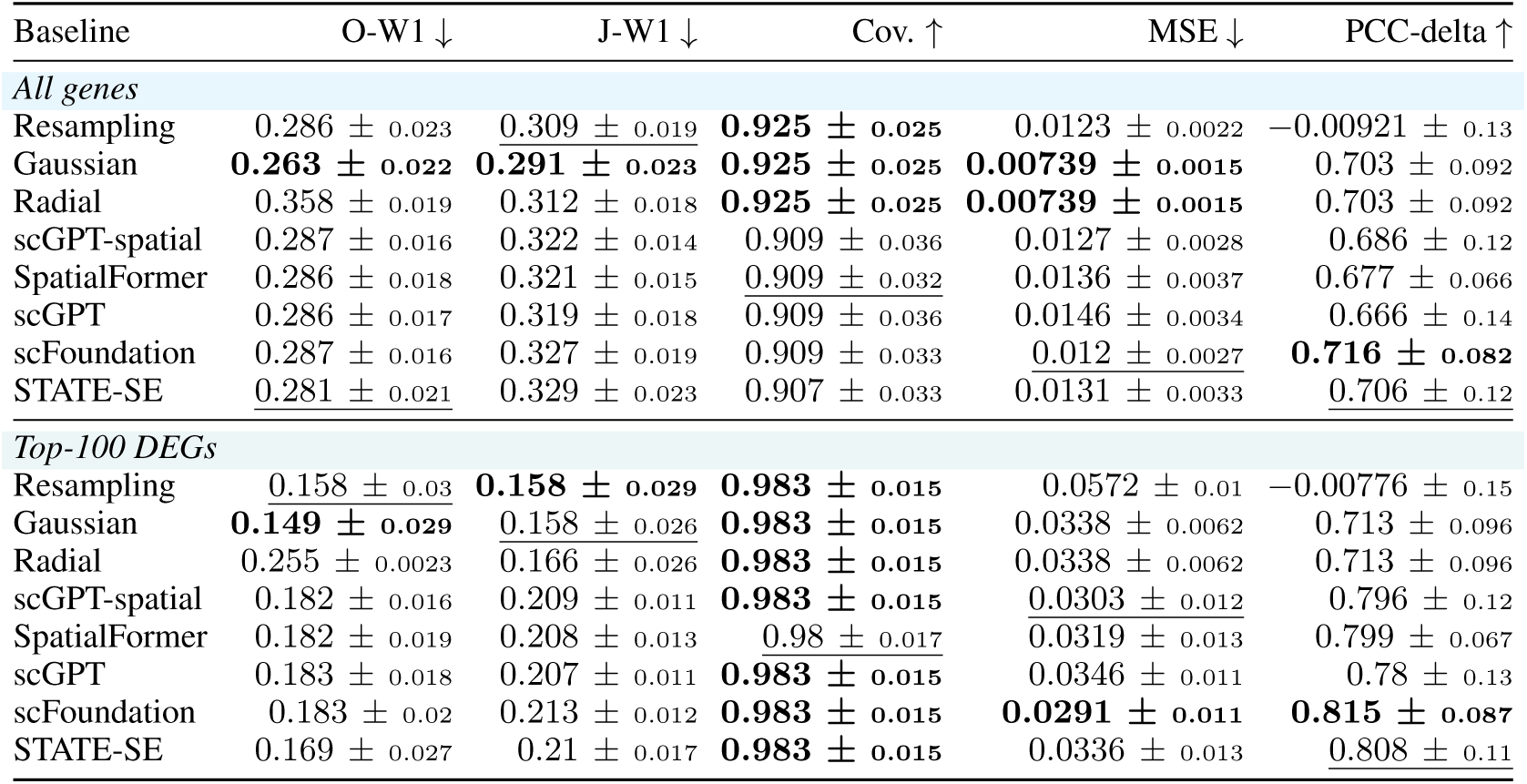
Benchmark results on Adipose.

**Table 29:** Benchmark results on UC-VDZ.

| Baseline | O-W1 ↓ | J-W1 ↓ | Cov. ↑ | MSE ↓ | PCC-delta ↑ |
| --- | --- | --- | --- | --- | --- |
| *All genes* |  |  |  |  |  |
| Resampling | 0.375 ± 0.12 | 0.371 ± 0.14 | **0.997 ± 0.0047** | <u>0.0116 ± 0.0065</u> | -0.0942 ± 0.098 |
| Gaussian | 0.366 ± 0.12 | **0.367 ± 0.14** | **0.997 ± 0.0047** | **0.00708 ± 0.004** | <u>0.538 ± 0.1</u> |
| Radial | 0.452 ± 0.031 | <u>0.37 ± 0.14</u> | **0.997 ± 0.0047** | **0.00708 ± 0.004** | <u>0.538 ± 0.1</u> |
| scGPT-spatial | 0.357 ± 0.12 | 0.488 ± 0.14 | 0.741 ± 0.22 | 0.0157 ± 0.0075 | 0.522 ± 0.11 |
| SpatialFormer | 0.357 ± 0.12 | 0.481 ± 0.14 | 0.75 ± 0.21 | 0.0141 ± 0.0055 | **0.544 ± 0.13** |
| scGPT | **0.357 ± 0.12** | 0.48 ± 0.14 | 0.729 ± 0.23 | 0.0163 ± 0.0069 | 0.52 ± 0.13 |
| scFoundation | <u>0.357 ± 0.12</u> | 0.481 ± 0.13 | 0.771 ± 0.19 | 0.0135 ± 0.0049 | 0.512 ± 0.089 |
| STATE-SE | 0.357 ± 0.12 | 0.471 ± 0.14 | <u>0.778 ± 0.19</u> | 0.0136 ± 0.0042 | 0.519 ± 0.15 |
| *Top-100 DEGs* |  |  |  |  |  |
| Resampling | 0.369 ± 0.14 | <u>0.375 ± 0.17</u> | **0.996 ± 0.0064** | 0.0844 ± 0.051 | -0.224 ± 0.19 |
| Gaussian | 0.36 ± 0.14 | **0.372 ± 0.17** | **0.996 ± 0.0064** | 0.041 ± 0.027 | **0.786 ± 0.078** |
| Radial | 0.435 ± 0.058 | 0.375 ± 0.17 | **0.996 ± 0.0064** | 0.041 ± 0.027 | **0.786 ± 0.078** |
| scGPT-spatial | 0.352 ± 0.13 | 0.443 ± 0.19 | 0.795 ± 0.17 | 0.034 ± 0.017 | 0.743 ± 0.21 |
| SpatialFormer | 0.353 ± 0.13 | 0.413 ± 0.16 | 0.791 ± 0.18 | **0.029 ± 0.015** | 0.755 ± 0.23 |
| scGPT | **0.352 ± 0.13** | 0.42 ± 0.17 | 0.78 ± 0.19 | 0.0321 ± 0.013 | <u>0.758 ± 0.19</u> |
| scFoundation | <u>0.352 ± 0.13</u> | 0.429 ± 0.17 | <u>0.804 ± 0.17</u> | <u>0.031 ± 0.013</u> | 0.744 ± 0.22 |
| STATE-SE | <u>0.353 ± 0.13</u> | 0.399 ± 0.15 | 0.801 ± 0.17 | 0.0321 ± 0.016 | 0.703 ± 0.3 |

**Table 30:** Benchmark results on Spinal-KO.

| Baseline | O-W1 ↓ | J-W1 ↓ | Cov. ↑ | MSE ↓ | PCC-delta ↑ |
| --- | --- | --- | --- | --- | --- |
| *All genes* |  |  |  |  |  |
| Resampling | 0.348 ± 0.022 | 0.375 ± 0.017 | **0.935 ± 0.015** | 0.0201 ± 0.0087 | 0.0392 ± 0.13 |
| Gaussian | **0.33 ± 0.019** | 0.358 ± 0.018 | **0.935 ± 0.015** | **0.0117 ± 0.0028** | **0.511 ± 0.18** |
| Radial | 0.396 ± 0.0059 | 0.377 ± 0.016 | **0.935 ± 0.015** | **0.0117 ± 0.0028** | **0.511 ± 0.18** |
| scGPT-spatial | 0.338 ± 0.02 | 0.315 ± 0.0027 | 0.757 ± 0.024 | 0.0353 ± 0.013 | 0.407 ± 0.22 |
| SpatialFormer | 0.338 ± 0.02 | **0.312 ± 0.0034** | 0.765 ± 0.032 | 0.0372 ± 0.016 | 0.41 ± 0.2 |
| scGPT | 0.338 ± 0.02 | 0.323 ± 0.0056 | 0.723 ± 0.02 | 0.0337 ± 0.012 | 0.401 ± 0.24 |
| scFoundation | 0.338 ± 0.02 | 0.32 ± 0.0051 | 0.723 ± 0.043 | 0.0333 ± 0.011 | 0.396 ± 0.24 |
| STATE-SE | 0.338 ± 0.02 | 0.319 ± 0.0043 | 0.772 ± 0.047 | 0.0345 ± 0.014 | 0.425 ± 0.21 |
| *Top-100 DEGs* |  |  |  |  |  |
| Resampling | 0.294 ± 0.017 | 0.305 ± 0.024 | **0.963 ± 0.03** | 0.0585 ± 0.025 | 0.0563 ± 0.17 |
| Gaussian | **0.283 ± 0.017** | 0.294 ± 0.022 | **0.963 ± 0.03** | **0.031 ± 0.0061** | **0.595 ± 0.19** |
| Radial | 0.342 ± 0.0097 | 0.311 ± 0.022 | **0.963 ± 0.03** | **0.031 ± 0.0061** | **0.595 ± 0.19** |
| scGPT-spatial | 0.288 ± 0.016 | **0.265 ± 0.021** | 0.849 ± 0.011 | 0.0494 ± 0.019 | 0.526 ± 0.24 |
| SpatialFormer | 0.288 ± 0.016 | 0.266 ± 0.026 | 0.865 ± 0.0073 | 0.0505 ± 0.018 | 0.533 ± 0.2 |
| scGPT | 0.288 ± 0.016 | 0.271 ± 0.02 | 0.826 ± 0.041 | 0.0502 ± 0.022 | 0.515 ± 0.25 |
| scFoundation | 0.288 ± 0.016 | 0.271 ± 0.013 | 0.82 ± 0.053 | 0.0498 ± 0.022 | 0.51 ± 0.26 |
| STATE-SE | 0.288 ± 0.016 | 0.272 ± 0.015 | 0.875 ± 0.013 | 0.0475 ± 0.019 | 0.544 ± 0.22 |

**Table 31:** Benchmark results on GBM-oHSV.

| Baseline | O-W1 ↓ | J-W1 ↓ | Cov. ↑ | MSE ↓ | PCC-delta ↑ |
| --- | --- | --- | --- | --- | --- |
| *All genes* |  |  |  |  |  |
| Resampling | 0.209 ± 0.011 | 0.253 ± 0.081 | **0.929 ± 0.083** | 0.0097 ± 0.009 | -0.0846 ± 0.27 |
| Gaussian | **0.202 ± 0.011** | **0.251 ± 0.08** | **0.929 ± 0.083** | **0.00897 ± 0.0091** | **0.29 ± 0.22** |
| Radial | 0.358 ± 0.0048 | 0.255 ± 0.082 | **0.929 ± 0.083** | **0.00897 ± 0.0091** | **0.29 ± 0.22** |
| scGPT-spatial | 0.205 ± 0.012 | 0.254 ± 0.081 | 0.922 ± 0.078 | 0.0272 ± 0.0081 | 0.242 ± 0.5 |
| SpatialFormer | 0.205 ± 0.012 | 0.267 ± 0.069 | 0.903 ± 0.062 | 0.0309 ± 0.0098 | 0.234 ± 0.49 |
| scGPT | 0.207 ± 0.013 | 0.256 ± 0.079 | 0.919 ± 0.076 | 0.0294 ± 0.0081 | 0.237 ± 0.49 |
| scFoundation | 0.205 ± 0.012 | 0.257 ± 0.078 | 0.924 ± 0.08 | 0.0269 ± 0.0086 | 0.249 ± 0.5 |
| STATE-SE | 0.204 ± 0.012 | 0.252 ± 0.082 | 0.923 ± 0.079 | 0.0271 ± 0.0091 | 0.248 ± 0.5 |
| *Top-100 DEGs* |  |  |  |  |  |
| Resampling | 0.147 ± 0.091 | 0.185 ± 0.12 | **0.81 ± 0.33** | 0.0355 ± 0.036 | -0.2 ± 0.34 |
| Gaussian | **0.143 ± 0.086** | 0.187 ± 0.12 | **0.81 ± 0.33** | **0.0324 ± 0.038** | **0.39 ± 0.39** |
| Radial | 0.304 ± 0.078 | 0.19 ± 0.13 | **0.81 ± 0.33** | **0.0324 ± 0.038** | **0.39 ± 0.39** |
| scGPT-spatial | 0.154 ± 0.078 | 0.184 ± 0.13 | **0.81 ± 0.33** | 0.0635 ± 0.016 | 0.237 ± 0.71 |
| SpatialFormer | 0.154 ± 0.078 | 0.192 ± 0.12 | 0.803 ± 0.32 | 0.0678 ± 0.0085 | 0.233 ± 0.71 |
| scGPT | 0.158 ± 0.074 | 0.184 ± 0.13 | **0.81 ± 0.33** | 0.0658 ± 0.011 | 0.237 ± 0.71 |
| scFoundation | 0.154 ± 0.077 | 0.183 ± 0.13 | **0.81 ± 0.33** | 0.0648 ± 0.015 | 0.239 ± 0.71 |
| STATE-SE | 0.153 ± 0.077 | **0.183 ± 0.13** | **0.81 ± 0.33** | 0.0642 ± 0.02 | 0.246 ± 0.72 |

**Table 32:** Benchmark results on Breast-FUS.

| Baseline | O-W1 ↓ | J-W1 ↓ | Cov. ↑ | MSE ↓ | PCC-delta ↑ |
| --- | --- | --- | --- | --- | --- |
| *All genes* |  |  |  |  |  |
| Resampling | 0.526 ± 0.13 | 0.608 ± 0.087 | 0.777 ± 0.21 | **0.000555 ± 0.00017** | 0.0463 ± 0.088 |
| Gaussian | 0.521 ± 0.13 | 0.604 ± 0.086 | 0.777 ± 0.21 | <u>0.000722 ± 8.3×10<sup>-5</sup></u> | **0.162 ± 0.1** |
| Radial | 0.547 ± 0.015 | 0.607 ± 0.086 | 0.777 ± 0.21 | <u>0.000722 ± 8.3×10<sup>-5</sup></u> | **0.162 ± 0.1** |
| scGPT-spatial | 0.505 ± 0.12 | 0.579 ± 0.071 | 0.779 ± 0.2 | 0.00619 ± 0.0037 | 0.0447 ± 0.2 |
| SpatialFormer | **0.505 ± 0.12** | <u>0.576 ± 0.071</u> | <u>0.782 ± 0.2</u> | 0.00681 ± 0.0043 | 0.0114 ± 0.15 |
| scGPT | 0.505 ± 0.12 | **0.574 ± 0.076** | **0.784 ± 0.2** | 0.00799 ± 0.0051 | -0.00783 ± 0.16 |
| scFoundation | 0.505 ± 0.12 | 0.582 ± 0.077 | 0.781 ± 0.19 | 0.00767 ± 0.003 | 0.0257 ± 0.18 |
| STATE-SE | 0.505 ± 0.12 | 0.58 ± 0.074 | 0.778 ± 0.19 | 0.00622 ± 0.0027 | <u>0.0477 ± 0.17</u> |
| *Top-100 DEGs* |  |  |  |  |  |
| Resampling | 0.404 ± 0.13 | 0.491 ± 0.098 | 0.719 ± 0.28 | **0.00353 ± 0.0012** | -0.0405 ± 0.24 |
| Gaussian | 0.401 ± 0.13 | 0.491 ± 0.097 | 0.719 ± 0.28 | <u>0.00369 ± 0.00061</u> | **0.144 ± 0.17** |
| Radial | 0.513 ± 0.016 | 0.491 ± 0.097 | 0.719 ± 0.28 | <u>0.00369 ± 0.00061</u> | **0.144 ± 0.17** |
| scGPT-spatial | <u>0.39 ± 0.12</u> | 0.473 ± 0.1 | 0.73 ± 0.26 | 0.014 ± 0.006 | -0.0305 ± 0.25 |
| SpatialFormer | **0.39 ± 0.12** | 0.471 ± 0.095 | <u>0.738 ± 0.25</u> | 0.0154 ± 0.0077 | -0.061 ± 0.17 |
| scGPT | 0.39 ± 0.12 | **0.464 ± 0.1** | **0.738 ± 0.25** | 0.0179 ± 0.0091 | -0.103 ± 0.2 |
| scFoundation | 0.39 ± 0.12 | <u>0.468 ± 0.1</u> | 0.725 ± 0.25 | 0.0191 ± 0.0066 | -0.0346 ± 0.17 |
| STATE-SE | 0.39 ± 0.12 | 0.472 ± 0.098 | 0.719 ± 0.26 | 0.0155 ± 0.0041 | <u>0.00356 ± 0.15</u> |

**Table 33:** Benchmark results on Gut-Microbiota.

| Baseline | O-W1 ↓ | J-W1 ↓ | Cov. ↑ | MSE ↓ | PCC-delta ↑ |
| --- | --- | --- | --- | --- | --- |
| *All genes* |  |  |  |  |  |
| Resampling | 0.559 ± 0.019 | 0.532 ± 0.0073 | 0.603 ± 0.085 | <u>0.00134 ± 0.00062</u> | -0.08 ± 0.24 |
| Gaussian | **0.523 ± 0.02** | 0.509 ± 0.0097 | 0.602 ± 0.085 | **0.00129 ± 0.00062** | 0.108 ± 0.069 |
| Radial | <u>0.525 ± 0.0051</u> | 0.537 ± 0.0087 | 0.602 ± 0.085 | **0.00129 ± 0.00062** | 0.108 ± 0.069 |
| scGPT-spatial | 0.546 ± 0.018 | 0.515 ± 0.0086 | <u>0.61 ± 0.082</u> | 0.00498 ± 0.0043 | -0.00175 ± 0.3 |
| SpatialFormer | 0.546 ± 0.018 | **0.505 ± 0.0062** | **0.612 ± 0.084** | 0.00654 ± 0.0047 | **0.231 ± 0.1** |
| scGPT | 0.546 ± 0.018 | 0.516 ± 0.01 | 0.605 ± 0.071 | 0.00338 ± 0.0025 | -0.104 ± 0.26 |
| scFoundation | 0.546 ± 0.018 | <u>0.508 ± 0.01</u> | 0.608 ± 0.08 | 0.00347 ± 0.0003 | <u>0.194 ± 0.093</u> |
| STATE-SE | 0.546 ± 0.018 | 0.515 ± 0.013 | 0.608 ± 0.09 | 0.00493 ± 0.0031 | 0.19 ± 0.17 |
| *Top-100 DEGs* |  |  |  |  |  |
| Resampling | 0.234 ± 0.099 | 0.312 ± 0.091 | 0.843 ± 0.13 | <u>0.0194 ± 0.01</u> | -0.0771 ± 0.33 |
| Gaussian | 0.232 ± 0.095 | 0.309 ± 0.086 | 0.843 ± 0.13 | **0.0186 ± 0.01** | 0.212 ± 0.06 |
| Radial | 0.468 ± 0.016 | 0.327 ± 0.094 | 0.843 ± 0.13 | **0.0186 ± 0.01** | 0.212 ± 0.06 |
| scGPT-spatial | **0.23 ± 0.096** | 0.291 ± 0.077 | 0.85 ± 0.13 | 0.0269 ± 0.015 | 0.00336 ± 0.39 |
| SpatialFormer | 0.23 ± 0.096 | 0.291 ± 0.084 | **0.853 ± 0.13** | 0.0333 ± 0.0068 | **0.291 ± 0.087** |
| scGPT | 0.23 ± 0.096 | 0.298 ± 0.073 | <u>0.852 ± 0.12</u> | 0.0274 ± 0.015 | -0.106 ± 0.34 |
| scFoundation | 0.23 ± 0.096 | **0.288 ± 0.082** | 0.85 ± 0.13 | 0.026 ± 0.008 | <u>0.286 ± 0.061</u> |
| STATE-SE | 0.23 ± 0.096 | 0.298 ± 0.1 | 0.837 ± 0.12 | 0.0316 ± 0.0078 | 0.283 ± 0.12 |

**Table 34:** Benchmark results on Cardiac-MI.

| Baseline | O-W1 ↓ | J-W1 ↓ | Cov. ↑ | MSE ↓ | PCC-delta ↑ |
| --- | --- | --- | --- | --- | --- |
| *All genes* |  |  |  |  |  |
| Resampling | 0.333 ± 0.061 | 0.392 ± 0.064 | **0.872 ± 0.13** | 0.0177 ± 0.015 | −0.152 ± 0.27 |
| Gaussian | **0.302 ± 0.063** | 0.372 ± 0.069 | **0.872 ± 0.13** | **0.0175 ± 0.015** | **0.145 ± 0.22** |
| Radial | 0.429 ± 0.017 | 0.395 ± 0.065 | **0.872 ± 0.13** | **0.0175 ± 0.015** | **0.145 ± 0.22** |
| scGPT-spatial | 0.32 ± 0.058 | 0.325 ± 0.014 | 0.836 ± 0.12 | 0.066 ± 0.012 | −0.023 ± 0.23 |
| SpatialFormer | 0.32 ± 0.058 | **0.322 ± 0.017** | 0.832 ± 0.14 | 0.0648 ± 0.011 | −0.0156 ± 0.23 |
| scGPT | 0.32 ± 0.058 | 0.325 ± 0.019 | 0.832 ± 0.13 | 0.0657 ± 0.0094 | −0.00718 ± 0.21 |
| scFoundation | 0.32 ± 0.058 | <u>0.324 ± 0.019</u> | 0.833 ± 0.13 | 0.0656 ± 0.0098 | <u>−0.00216 ± 0.17</u> |
| STATE-SE | <u>0.32 ± 0.058</u> | 0.324 ± 0.018 | <u>0.844 ± 0.13</u> | 0.0607 ± 0.0072 | −0.00681 ± 0.2 |
| *Top-100 DEGs* |  |  |  |  |  |
| Resampling | 0.165 ± 0.067 | 0.231 ± 0.083 | 0.961 ± 0.035 | <u>0.0789 ± 0.067</u> | −0.188 ± 0.32 |
| Gaussian | **0.163 ± 0.069** | 0.231 ± 0.084 | 0.961 ± 0.035 | **0.0786 ± 0.069** | **0.108 ± 0.2** |
| Radial | 0.348 ± 0.025 | 0.238 ± 0.084 | 0.961 ± 0.035 | **0.0786 ± 0.069** | **0.108 ± 0.2** |
| scGPT-spatial | 0.173 ± 0.058 | 0.16 ± 0.039 | **0.978 ± 0.013** | 0.233 ± 0.044 | −0.127 ± 0.45 |
| SpatialFormer | 0.174 ± 0.058 | 0.16 ± 0.044 | <u>0.976 ± 0.013</u> | 0.226 ± 0.037 | −0.11 ± 0.45 |
| scGPT | 0.174 ± 0.058 | <u>0.159 ± 0.043</u> | <u>0.973 ± 0.016</u> | 0.228 ± 0.033 | <u>−0.098 ± 0.43</u> |
| scFoundation | 0.173 ± 0.058 | 0.16 ± 0.041 | 0.974 ± 0.018 | 0.226 ± 0.034 | −0.0987 ± 0.37 |
| STATE-SE | 0.173 ± 0.058 | **0.158 ± 0.042** | 0.971 ± 0.017 | 0.215 ± 0.025 | −0.108 ± 0.41 |

**Table 35:** Benchmark results on Kidney-IRI.

| Baseline | O-W1 ↓ | J-W1 ↓ | Cov. ↑ | MSE ↓ | PCC-delta ↑ |
| --- | --- | --- | --- | --- | --- |
| *All genes* |  |  |  |  |  |
| Resampling | 0.18 ± 0.036 | 0.256 ± 0.13 | **0.941 ± 0.099** | 0.0403 ± 0.025 | −0.0751 ± 0.19 |
| Gaussian | **0.167 ± 0.033** | 0.247 ± 0.13 | **0.941 ± 0.099** | 0.0287 ± 0.023 | 0.569 ± 0.095 |
| Radial | 0.427 ± 0.0049 | 0.262 ± 0.13 | **0.941 ± 0.099** | 0.0287 ± 0.023 | 0.569 ± 0.095 |
| scGPT-spatial | 0.176 ± 0.033 | 0.242 ± 0.048 | 0.852 ± 0.11 | 0.0286 ± 0.0066 | <u>0.664 ± 0.08</u> |
| SpatialFormer | 0.176 ± 0.033 | **0.236 ± 0.038** | 0.854 ± 0.11 | <u>0.0284 ± 0.0066</u> | **0.665 ± 0.074** |
| scGPT | 0.176 ± 0.033 | 0.244 ± 0.05 | 0.869 ± 0.086 | **0.0275 ± 0.0058** | 0.664 ± 0.095 |
| scFoundation | <u>0.176 ± 0.033</u> | 0.238 ± 0.041 | 0.866 ± 0.11 | 0.0285 ± 0.0069 | 0.656 ± 0.086 |
| STATE-SE | 0.176 ± 0.033 | <u>0.236 ± 0.042</u> | <u>0.875 ± 0.096</u> | 0.0302 ± 0.0081 | 0.648 ± 0.081 |
| *Top-100 DEGs* |  |  |  |  |  |
| Resampling | <u>0.111 ± 0.057</u> | 0.169 ± 0.098 | 0.871 ± 0.22 | 0.109 ± 0.058 | −0.0588 ± 0.19 |
| Gaussian | **0.104 ± 0.05** | 0.16 ± 0.088 | 0.871 ± 0.22 | 0.0762 ± 0.054 | 0.549 ± 0.1 |
| Radial | 0.382 ± 0.012 | 0.18 ± 0.093 | 0.871 ± 0.22 | 0.0762 ± 0.054 | 0.549 ± 0.1 |
| scGPT-spatial | 0.117 ± 0.049 | 0.129 ± 0.026 | 0.874 ± 0.17 | 0.0571 ± 0.01 | <u>0.689 ± 0.079</u> |
| SpatialFormer | 0.117 ± 0.049 | <u>0.128 ± 0.022</u> | 0.881 ± 0.17 | <u>0.0565 ± 0.01</u> | <u>0.688 ± 0.073</u> |
| scGPT | 0.117 ± 0.049 | 0.129 ± 0.021 | 0.881 ± 0.17 | **0.0546 ± 0.0091** | **0.69 ± 0.09** |
| scFoundation | 0.117 ± 0.049 | 0.131 ± 0.018 | **0.888 ± 0.18** | 0.0569 ± 0.011 | 0.682 ± 0.082 |
| STATE-SE | 0.117 ± 0.049 | **0.128 ± 0.021** | <u>0.885 ± 0.16</u> | 0.0604 ± 0.013 | 0.673 ± 0.083 |

### H.1 Dependence on the evaluated gene panel

#### Purpose and comparison

We ask whether an Oracle spatial improvement averaged over all measured genes also holds for genes ranked highly by their abundance changes. The five foundation-model adapters on 14 datasets give 70 model–dataset comparisons. Each comparison uses the same saved predictions and responses under both panels; only the evaluated genes change. Thus this is a scoring sensitivity check, not additional model training.

#### Reading the figure

In Figure 10, the horizontal axis is split-paired percentage improvement over Resampling on all genes, and the vertical axis is the same quantity on the top-100 abundance-ranked panel. Positive values mean improvement. A point in the lower-right quadrant therefore represents a model that improves the all-gene score but worsens the score on the ranked panel.

**Figure 10:**
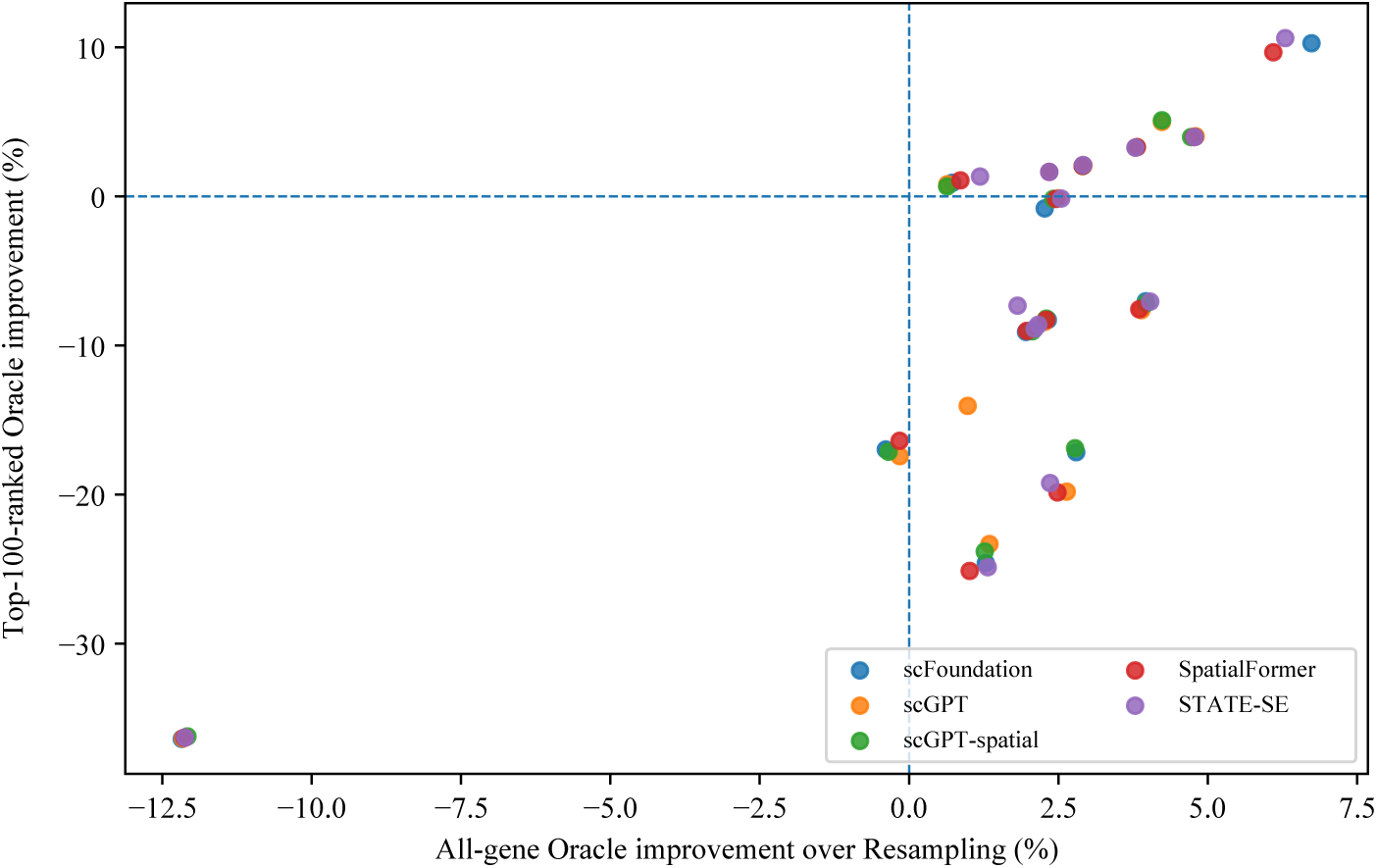
Spatial gains depend on the evaluated genes. Each point is one foundation-model–dataset combination. Both axes show split-paired percentage improvement in Oracle W1 over Resampling, with positive values indicating lower error. Lower-right points improve on all genes but worsen on the top-100 abundance-ranked panel. The comparisons reuse the same datasets, responses, and saved predictions.

#### Results and aggregation sensitivity

Of 61 model–dataset comparisons with positive all-gene improvement, 31 become negative on the top-100 panel. An all-gene spatial gain therefore does not uniformly extend to abundance-ranked genes. The ranking is based on abundance after predictions are fixed; it is not a discovery test for spatial-response genes.

The order of averaging also matters. Our primary summary computes the percentage improvement within each split and then averages those percentages, as defined in Equation (16). An alternative first averages each method’s raw errors across splits and then takes their ratio. That alternative implicitly gives more weight to splits with larger Resampling error and yields 26 reversals instead of 31. All five differences arise from DSS-Colitis. The two counts describe different summaries of the same predictions and are not independent experimental replications.

